# Synthetic genomes unveil the effects of synonymous recoding

**DOI:** 10.1101/2024.06.16.599206

**Authors:** Akos Nyerges, Anush Chiappino-Pepe, Bogdan Budnik, Maximilien Baas-Thomas, Regan Flynn, Shirui Yan, Nili Ostrov, Min Liu, Meizhou Wang, Qingmei Zheng, Fangxiang Hu, Kangming Chen, Alexandra Rudolph, Dawn Chen, Jenny Ahn, Owen Spencer, Venkat Ayalavarapu, Angela Tarver, Miranda Harmon-Smith, Matthew Hamilton, Ian Blaby, Yasuo Yoshikuni, Behnoush Hajian, Adeline Jin, Balint Kintses, Monika Szamel, Viktoria Seregi, Yue Shen, Zilong Li, George M. Church

## Abstract

Engineering the genetic code of an organism provides the basis for (i) making any organism safely resistant to natural viruses and (ii) preventing genetic information flow into and out of genetically modified organisms while (iii) allowing the biosynthesis of genetically encoded unnatural polymers^1–4^. Achieving these three goals requires the reassignment of multiple of the 64 codons nature uses to encode proteins. However, synonymous codon replacement—recoding—is frequently lethal, and how recoding impacts fitness remains poorly explored. Here, we explore these effects using whole-genome synthesis, multiplexed directed evolution, and genome-transcriptome-translatome-proteome co-profiling on multiple recoded genomes. Using this information, we assemble a synthetic *Escherichia coli* genome in seven sections using only 57 codons to encode proteins. By discovering the rules responsible for the lethality of synonymous recoding and developing a data-driven multi-omics-based genome construction workflow that troubleshoots synthetic genomes, we overcome the lethal effects of 62,007 synonymous codon swaps and 11,108 additional genomic edits. We show that synonymous recoding induces transcriptional noise including new antisense RNAs, leading to drastic transcriptome and proteome perturbation. As the elimination of select codons from an organism’s genetic code results in the widespread appearance of cryptic promoters, we show that synonymous codon choice may naturally evolve to minimize transcriptional noise. Our work provides the first genome-scale description of how synonymous codon changes influence organismal fitness and paves the way for the construction of functional genomes that provide genetic firewalls from natural ecosystems and safely produce biopolymers, drugs, and enzymes with an expanded chemistry.

## Introduction

The shared language of the genetic code uses 64 codons to encode the canonical 20 amino acids on the course of protein production. However, 18 of these amino acids and the translational start and stop signals are encoded by multiple synonymous codons^5,6^. This degeneracy of the genetic code allows coding sequences—through codon selection—to define protein production simultaneously while maintaining cellular homeostasis by regulating gene expression, translation, and protein folding^7,8^. Synonymous codon selection also influences cellular homeostasis by altering messenger (m)RNA structure, translation efficiency and speed, and accuracy^9^. Furthermore, RNA and DNA motifs provide additional noncoding functions such as regulating DNA and mRNA localization, genome replication, DNA repair, and chromosome segregation^7,10^.

The nearly universal nature and degeneracy of the genetic code spurred the exploration of synthetic genomes with artificial genetic codes to enhance biological functions. Studies demonstrated that reducing the number of genetic codons in an organism’s genetic code is feasible—by extensive genome editing or whole genome synthesis—paving the way for making organisms safely resistant to natural viruses, preventing genetic information flow into and out of Genetically Modified Organisms (GMOs), and enabling the biosynthesis of genetically encoded non-canonical polymers^1–4,11,12^.

However, despite the wide-ranging benefits of organisms with modified genetic codes, studies highlighted the collateral impact of genetic code engineering on organismal fitness. The creation of Genomically Recoded Organisms (GROs), whose genomes have been systematically redesigned to confer an alternate genetic code, in all cases induced simultaneous fitness reduction, decreased growth rate, and reduced biomass yield. The creation of the first GRO by our group through the genome-wide removal of 321 TAG stop codons and release factor 1 (RF1, encoded by *prfA*) from *Escherichia coli* K-12 MG1655 resulted in a 60% increase in cell doubling time compared to the parental strain^1^. Follow-up projects from our lab demonstrated that not only the rarest TAG stop codon but sense codons are also malleable—by attempting to recode 62 codons and eliminate 13 codons in 42 highly expressed prokaryotic genes and, in a separate study, by removing the rarest AGA and AGG arginine codons from growth-essential genes of *E. coli*—paving the way for sense codon recoding on whole genomes. This latter study revealed the occasionally lethal effect of synonymous recoding due to the role of rare codons in mRNA folding and translation initiation in natural transcripts. In that study, the replacement of only 2.9% of all target codons, *i.e.*, 123 arginine codons from the total 4,228 instances in the entire genome, resulted in a 28% increase in doubling time despite rounds of troubleshooting, including the selection of codons with the least impact on mRNA folding.

These early studies suggested that the compression of the genetic code in a living organism beyond the complete removal of rare stop codons might be feasible. Driven by this possibility, we initiated the design and synthesis of a complete *E. coli* genome, that we named Ec_Syn57, designed to rely on only 57 genetic codons^13^. Our early tests using a few select genomic regions from this strain, spanning approximately 1.1% of the entire genome, confirmed the viability of 63% of the tested recoded genes despite removing seven codons from the genetic code (**Figure 1a**). However, these small-scale tests also indicated frequent deleterious effects and our efforts to replace multiple recoded sections in a single strain sequentially remained unsuccessful. In parallel, a strain of *E. coli*, Ec_Syn61Δ3, was created with a synthetic recoded genome from which all annotated instances of two serine codons, TCG and TCA, and the TAG stop codon were eliminated to confer a 61-codon genetic code^14^. Successful genome recoding required the stepwise troubleshooting of four loci where codon swaps induced lethality, and despite correcting these, the final strain displayed a 73-153% increase in doubling time compared to its parental strain, depending on growth conditions^14^. Decreased fitness subsequently prevented the immediate deletion of the corresponding serine transfer (t)RNA genes (*serU* and *serT*) and RF1^3,14^, necessitating two rounds of additional troubleshooting before the first sense-codon-recoded strain could be completed.

**Figure 1.**
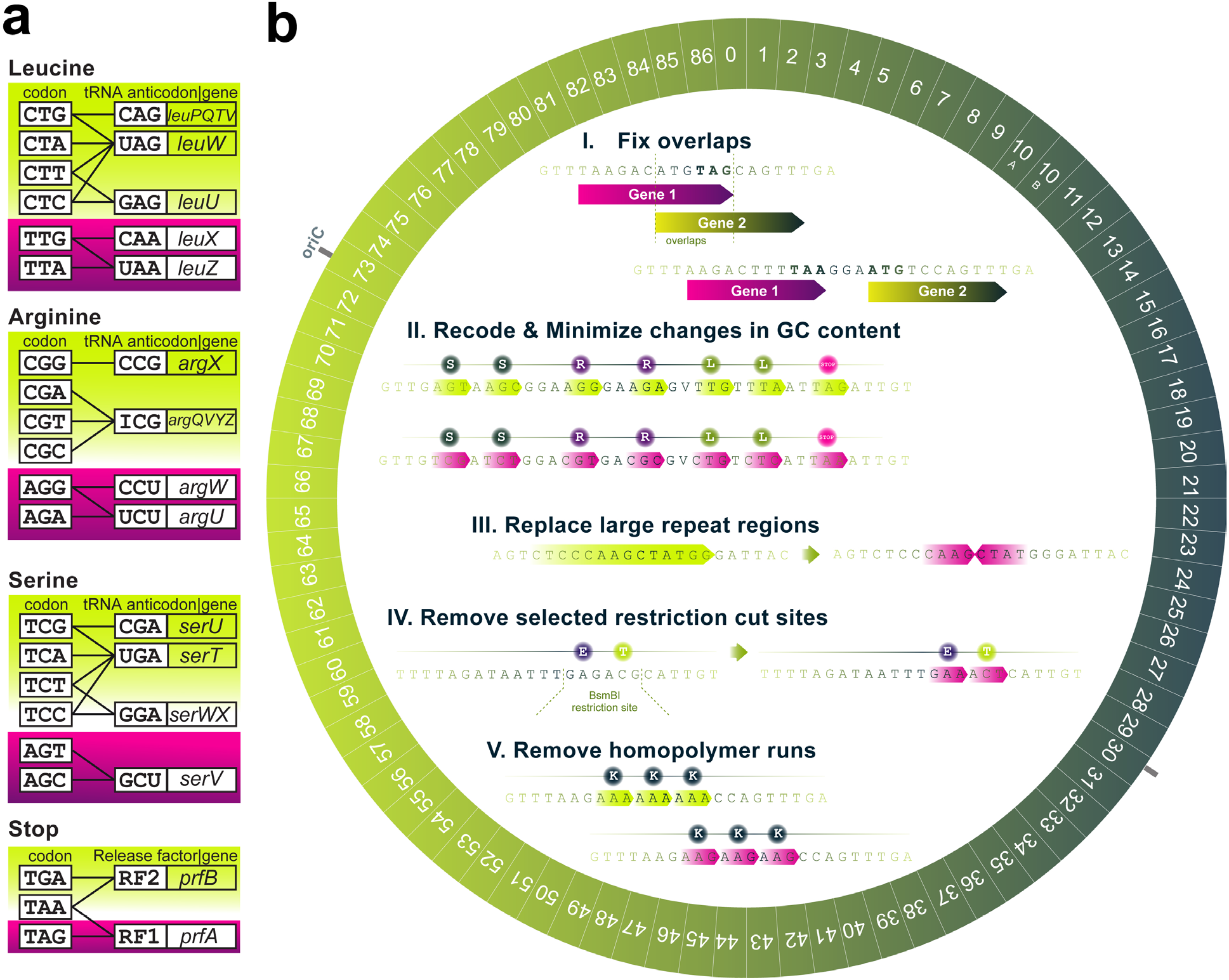
Design of a synthetic 57-codon *Escherichia coli* genome. (**a**) The design of Ec_Syn57’s compressed genetic code. Magenta marks codons and their corresponding tRNA genes and release factor I (RF1, encoded by *prfA*) selected for elimination during genome design. (**b**) The computational genome design of Ec_Syn57 synonymously recoded all known instances of the seven target codons highlighted in Figure 1a and streamlined the synthetic chromosome. Prior to synonymous recoding, (**I.**) overlapping genes that contain forbidden codons within their overlapping region were disentangled while preserving the downstream gene’s ribosomal binding site; next, (**II.**) protein-coding genes were recoded to confer the 57-codon genetic code of Figure 1a while minimizing local changes in GC% and mRNA folding differences near the 5’ end of genes. Next, we streamlined DNA synthesis and subsequent genome assembly steps by eliminating unstable repeats (**III.**), removing cut sites of AarI, BsaI, and BsmBI restriction enzymes (**IV.**), and eliminating sequences containing >8 consecutive As, Cs, Ts, or >5 consecutive Gs (**V.**). Finally, the refactored, recoded genome was divided into 86 ∼50 kbp segments, and the entire genome was synthesized.

Prior studies highlighted two fundamental challenges in front of our ability to construct synthetic genomes with custom functions and genetic codes^15–32^:

1. The precise *in vivo* assembly of mega-to-gigabase-scale genomes from short, synthetic DNA precursors; and
2. The rapid, rational, and parallelizable discovery and troubleshooting of fitness-decreasing design issues on synthetic genomes.

While multiple studies addressed necessary steps in genome assembly^15–33^, the identification of genome design issues and their troubleshooting remained unsolved. This, in turn, limits our ability to generate synthetic genomes distant from naturally occurring templates, hindering our capability to generate intrinsically safe biological systems for therapy, bioproduction, and bioremediation.

In this paper, we address these challenges by developing a data-driven genome synthesis and troubleshooting technology that rapidly identifies and corrects errors in synthetic genomes. We demonstrate that—by integrating multi-omics analyses along all steps of the central dogma (*i.e.*, genome, primary- and total-transcriptome, translatome, and proteome profiling) to discover fitness-decreasing errors and correct these using multiplexed genome editing—the complete synthesis of a functional *Escherichia coli* genome is possible in seven strains in which all 62,007 instances of seven codons have been synonymously replaced. We discover how synonymous codon replacement impacts gene expression and cellular protein production by uncovering the transcriptional, translational, and proteomic effects of a 61- and 57-codon genetic code. Finally, based on newly discovered genome design rules, we describe a computational and experimental workflow for designing high-fitness synthetic genomes.

This work provides a universal technology for constructing and troubleshooting genomes with user-defined genetic codes and the first functional example of a synthetic genome that uses 57 genetic codons for protein synthesis and liberates seven codons for subsequent reassignment. We expect that this study will pave the way toward organisms with custom functionalities, including multi-virus-resistant cells capable of producing biopolymers with an enhanced chemical repertoire and intrinsically safe GMOs by eliminating horizontal gene transfer via dependence on genetic codes and amino acids not found in nature^34^.

## Results

### Design and synthesis of a radically recoded genome

We previously designed a genome of *E. coli* MDS42 in which all 62,007 annotated instances of seven codons (5.35% of all codons) were replaced with synonymous alternatives across all protein-coding genes^13^. Based on our previously described design principles (**Figure 1a-b**)^13^, we installed an additional 11,108 modifications, resulting in a total of 162,521 basepair (bp) difference compared to the 3.98 Mbp parental genome. Codons were chosen for replacement based on their low frequency in protein-coding sequences to minimize the changes required (*i.e.,* the TAG stop, and the AGA and AGG arginine codons) and their orthogonality within the cellular tRNA pool (*i.e.,* the TTG and TTA leucine, and AGT and AGC serine codons) to ensure that their cognate tRNAs and Release Factor I could be later eliminated and all seven codons reassigned to new function without perturbing the cellular proteome (**Figure 1a**). Following computational design, we disconnected the synthetic genome at operon borders into 87 segments between 31 kbp and 52 kbp in length. Sixty segments were assembled previously into bacterial artificial chromosomes (BACs) using chemical DNA synthesis and homologous recombination in *Saccharomyces cerevisiae*^13^. For the remaining 27 segments in this work, we updated our genome synthesis workflow by i.) utilizing a new assembly strategy that utilizes 10-times longer overlaps (that is, 500 bp compared to the previous 50 bp) and sequence-validated clonal DNA fragments, allowing us to reduce synthesis error rate 19-fold and achieve an average error rate of 0.8 compared to 15.4 errors per segment before, and ii.) increase segment stability following yeast assembly. In line with previous reports showing that endogenous bacterial mobile genetic elements (MGEs)— insertion sequences (ISes) and transposons—frequently invade synthetic DNA constructs^35–37^, we detected extensive MGE transposition into newly synthesized chromosomal regions following segment delivery into *E. coli.* These transposition events were mainly driven by the Tn1000 transposon of *E. coli* DH10B^38,39^, a widely used cloning host designed to propagate large BACs (**Supplementary Figure 1a**). Transpositions occasionally disrupted growth-essential genes in synthetic constructs, preventing subsequent genome construction (**Supplementary Figure 1b**). We therefore eliminated transposon mutagenesis from strains carrying parts of our synthetic genome using CRISPR/Cas9-assisted MAGE to delete all 24 instances of transposed mobile genetic elements, and utilized mobile-genetic-element-free cloning hosts to maintain recoded segments^40,41^. Our updated workflow allowed us to assemble all but one of the 87 segments smoothly. Segment 10 was challenging to synthesize, and we therefore constructed it by splitting its sequence into two fragments, a 21 kbp (part A) and 24 kbp (part B). Finally, we also constructed a 90.1 kbp neochromosomal region carrying genes and operons that were disrupted by DNA synthesis errors on 24 previously synthesized segments, under the control of their native promoter.

Using the updated genome synthesis workflow, we completed the synthesis of an *E. coli* genome in which 62,007 instances of seven codons (*i.e.*, the AGT, AGC (Serine), TTG, TTA (Leucine), AGG, AGA (Arginine), and TAG (Stop)) were recoded to synonymous alternatives and 11,108 additional modifications were installed to maintain viability, ease DNA synthesis, and streamline the final synthetic chromosome.

### Genome construction with frequent design issues

With the 89 BACs containing our synthetic genome, we initiated genome assembly in eleven strains using a method that relies on CRISPR/Cas9-assisted-recombineering, multiomics-based rational editing- and adaptive-laboratory-evolution-based troubleshooting. The development of a combined genome construction and data-driven troubleshooting method was necessary based on our previous estimate of approximately one lethal and multiple fitness-decreasing issues caused by synonymous recoding in each ∼50-kbp segment^13^. Existing prokaryotic genome construction methods (*i.e.*, REXER, GENESIS, SIRCAS, CONEXER, and CGS) require tedious, stepwise troubleshooting consisting of the experimental detection of the locus containing a lethal design error and then troubleshooting these errors one-by-one (**Supplementary Table 1**)^13,14,21,42^. Furthermore, existing methods do not employ counterselection against the parental sequence and rely on the induced crossover between synthetic and wild-type chromosomal regions, rendering them unsuitable for the rapid, sequential construction and troubleshooting of synthetic genomic regions containing multiple closely spaced fitness-decreasing errors. Without sequence-based counterselection against the wild-type genomic locus, induced crossovers frequently result in chimeric genomes and the complete reversion of fitness-impairing codon changes, preventing the construction of genomes with less-tolerated recoding schemes^21,43^ (**Figure 2a**). For example, previous REXER-based efforts to replace all TTG and TTA (Leu) codons within *E. coli*’s *mraZ*-*ftsZ* locus, encoding growth-essential cell wall components, using a synonymous recoding scheme that closely follows our 57-codon design failed catastrophically, yielded no recoded variants, and the reasons behind nonviability remained undiscovered^43^ (**Supplementary Table 2**). Therefore, we sought to develop a data-informed genome synthesis strategy that relies on the standardized, multiomics-based, high-throughput discovery of fitness-decreasing errors and their subsequent parallelized troubleshooting. We hypothesized that the stringent elimination of the parental copy of each synthetic genomic segment, without induced crossover between the parental and synthetic genome, would allow us to eliminate the parental copy even in the presence of near-lethal design errors (**Figure 2b**). The deletion of the parental copy would, in turn, pose a fitness-based selection pressure on correcting design issues in the synthetic chromosomal region, and this feature could then be exploited to troubleshoot design errors in a multiplexed manner via rational editing and Adaptive Laboratory Evolution (ALE). Once variants with improved fitness are identified, the induction of homologous recombination without the presence of the parental copy would allow the scarless implantation of synthetic genomic regions.

**Figure 2.**
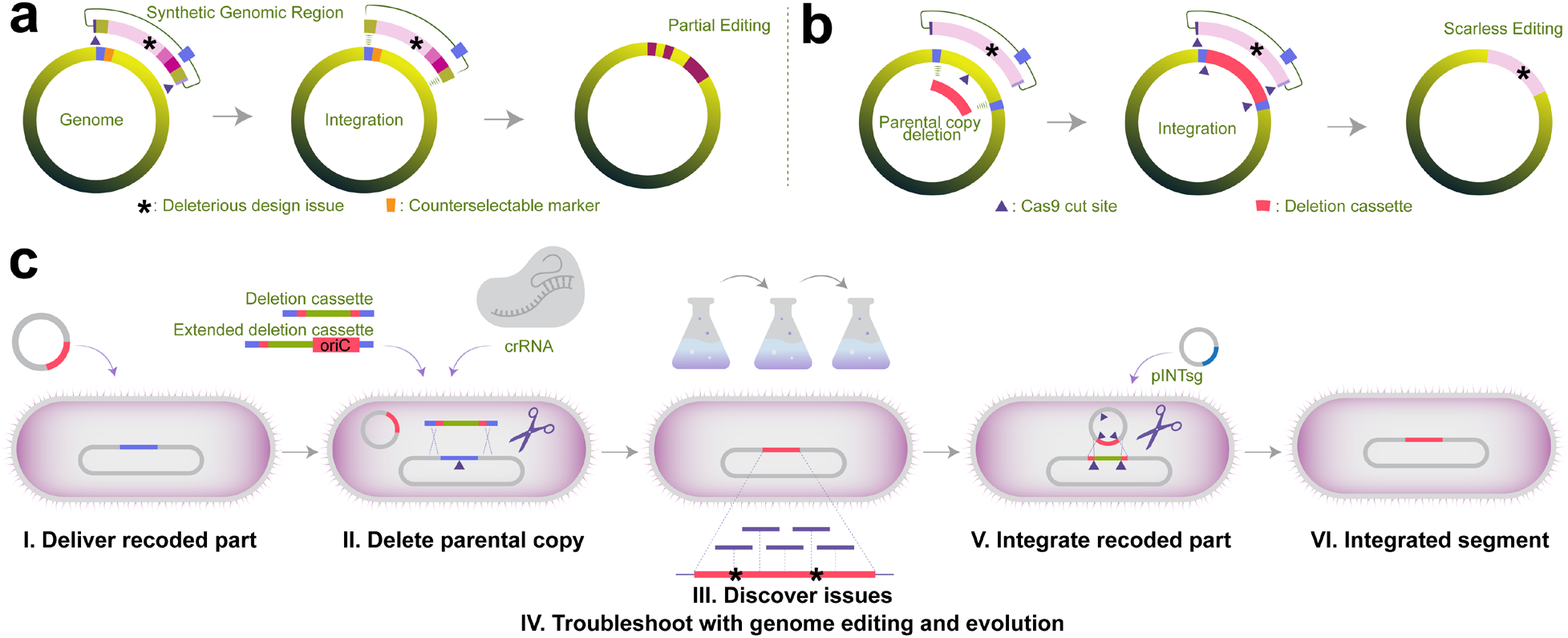
Genome construction with challenging recoding schemes. (**a**) Without sequence-based counterselection against the parental genome (*i.e.*, CRISPR/Cas9-cut within the parental copy, marked as ▴), genome construction methods that rely on induced crossover and the use of only counterselectable marker genes (marked in orange) to eliminate the parental copy frequently result in chimeric genomes and the reversion of challenging modifications and deleterious genome-design issues (marked as **\***), preventing the construction and troubleshooting of genomes with less-tolerated recoding schemes. (**b**) Sequential integration coupled with CRISPR/Cas9-cut assisted sequence-based counterselection against the parental locus prevents unwanted chimera formation and allows the generation of fitness-decreasing recoding schemes. (**c**) The workflow of SynOMICS-based genome construction and troubleshooting. SynOMICS utilizes recombination deficient (*i.e.*, Δ*recA*) parental strains, leading to increased CRISPR/Cas9 selection stringency and preventing unwanted recombination between the parental and recoded copies. Recoded chromosomal segments are delivered using electroporation or conjugation (**I.**), followed by CRISPR/Cas9-assisted Lambda-Red recombineering-based deletion of the parental segment copy using an antibiotic-resistance-conferring deletion cassette and a genome-targeting crRNA plasmid (**II.**). Fitness-decreasing synonymous codons and genomic design errors are discovered using multi-omics analyses (**III.**) and troubleshot using multiplexed genome editing and laboratory evolution (**IV.**). Finally, the extrachromosomal recoded segment is integrated by delivering a 6-plex sgRNA expression plasmid (**V.**) that site-specifically integrates the synthetic part (**VI.**) and eliminates the genome-targeting crRNA plasmid of Step II. SynOMICS cycles are repeatable without interruption due to the inducible elimination of its pINTsg integration plasmid.

We addressed this challenge by developing **Syn**thetic genomes by multi**OM**ics-based **I**terative **C**onstruction and trouble**S**hooting or **SynOMICS** in short, a genome construction and data-driven troubleshooting method relying on CRISPR/Cas9-counterselection, recombineering, multi-omics-based rational troubleshooting, and ALE. In SynOMICS, following the delivery of a synthetic-segment-containing BAC (**Figure 2c**), the elimination of the corresponding parental chromosomal region is induced by CRISPR/Cas9-assisted recombineering. This step is initiated in cells harboring a constitutive Cas9 expression plasmid (termed pRedCas2, conferring chloramphenicol resistance (R)) by transiently overexpressing recombineering proteins and delivering a genome-targeting CRISPR-RNA-(crRNA)-expressing plasmid (conferring kanamycin-R) and a linear antibiotic marker cassette that confers gentamicin resistance. The antibiotic marker cassette serves three purposes. 1.) It eliminates—in combination with a Cas9 cut—the parental genomic region via homologous recombination, 2.) prepares the genomic locus for the subsequent integration of the extrachromosomal recoded region by integrating terminal homologies that match the last 200 basepairs of the recoded part, and 3.) allows the antibiotic-resistance-based identification of clones that lost the parental copy of the synthetic chromosomal segment. The use of recombination-deficient host strain^44^ and the development of pRedCas2—an all-in-one CRISPR/Cas9 and heat-inducible Lambda-Red expression plasmid—allowed us to achieve stringent selection and identify even rare variants that lost the parental segment copy. To demonstrate the stringency of the deletion step, we explored whether SynOMICS could generate recoding schemes that previously seemed nonviable based on a prior genome construction method^43^. The *mraZ-ftsZ* locus in Segment 2 served as an ideal target, as i.) it harbors the entire, previously tested *mraZ-ftsZ* locus, ii.) attempts the elimination of the same 157 instances of TTG, TTA (Leu) codons that failed in earlier recoding efforts^43^, and iii.) contains a total number of 585 codon changes, including the synonymous recoding of AGT, AGC (Ser), AGG, AGA (Arg), and TAG (Stop) codons besides leucine codon recoding, 273% more codon changes than what was previously attempted (**Supplementary Table 2**). SynOMICS-based deletion of the parental copy of Segment 2 yielded approximately 1000 colonies. Edited clones after the SynOMICS deletion step were easily identified based on their antibiotic resistance profile (*i.e.,* chloramphenicol-R, kanamycin-R, gentamicin-R) and colony PCR and confirmed using Multiplex Allele-Specific Colony (MASC) PCR^13,45^. 92% of all clones surviving antibiotic selection contained only the recoded allele (**Supplementary Table 2**), demonstrating SynOMICS’s ability to achieve challenging recoding schemes. As the elimination of the parental copy poses a growth-based, continuous selection pressure on the synthetic chromosomal region, we hypothesized that multiplexed genome editing and ALE would allow us to troubleshoot errors and identify improved variants after the SynOMICS deletion step (**Figure 2c**).

Based on the stringent selection posed by CRISPR/Cas9 and the multi-kbp payload capacity of the marker cassette, we expected that SynOMICS would also allow us to simultaneously correct design flaws and DNA synthesis errors within recoded segments, while testing the functionality of the recoded design. We demonstrated that the deletion cassette of SynOMICS (**Figure 2c**) is extendable to correct design and synthesis errors within BAC-carried segments. We utilized this feature to insert and replace genes and operons up to 6.7 kbp, correct DNA synthesis errors in eight segments, and complement lethal design flaws at four additional loci (Supplementary Methods). Temporary insertion at the deletion step allowed us to delete and recode the growth-essential genomic origin-of-replication (oriC) in Segment 73 and the dif (deletion-induced filamentation) site in Segment 28. This feature of SynOMICS is especially important, as it allows both the replacement of essential genomic motifs (*e.g.*, oriC) and the parallelized correction of errors by varying only the sequence of a short double-stranded (ds)DNA deletion cassette.

Following the deletion of the parental genomic locus, the scarless integration of the recoded segment—directed by the deletion-cassette-integrated 200 bp terminal homologies—is initiated by the delivery of a compact, CRISPR/Cas9 guide (g)RNA-expressing plasmid, pINTsg. pINTsg confers carbenicillin-R to allow selection in SynOMICS-deleted clones. pINTsg expresses six single guide RNAs (sgRNAs) simultaneously, targeting the two ends of the SynOMICS deletion cassette, two ends of the recoded segment, and the backbone of the BAC vector. At the same time, the sixth sgRNA cuts the deletion step’s crRNA plasmid (**Figure 2c**), preparing the cell for the next SynOMICS cycle. pINTsg also carries a tightly repressed bacterial toxin expression module for self-elimination following the integration step. As the selection stringency of CRISPR/Cas9-based selection is frequently limited by intramolecular recombination due to the sequence similarity between neighboring sgRNA expression constructs resulting in the loss of targeting sgRNAs^46,47^ (**Supplementary Figure 2a**), we maximized the efficiency of SynOMICS by utilizing a recently developed set of nonrepetitive sgRNAs and by eliminating unstable motifs and direct repeats longer than 11 bp^47–50^ (**Supplementary Figure 2b-c**). The elimination of unstable sites boosted SynOMICS’s integration efficiency, frequently approaching 100% (**Supplementary Figure 3**). Finally, once edited clones are identified based on their antibiotic resistance profile (chloramphenicol-R and carbenicillin-R) and verified by colony PCR (**Figure 2c, Supplementary Figure 3**), the induction of pINTsg’s toxin module prepares the cells for a new integration cycle.

Using SynOMICS, we initiated iterative single-segment replacements but observed drastic fitness impairment after recoding one to five neighboring ∼50-kbp segments, indicating the fitness-decreasing effects of synonymous recoding (for the detailed description of genome construction steps and the observed effects, see Supplementary Methods). As these fitness-decreasing errors prevented the replacement of additional genomic regions, we next investigated the sources of fitness-decreasing issues and corrected them using SynOMICS.

### Identification of design issues

We identified potential fitness-decreasing synonymous codon changes using simultaneous genome, transcriptome, translatome, and proteome profiling (**Figure 3a**) on multiple recoded genomes and corrected the identified issues using genome editing and ALE. We hypothesized that synonymous codon swaps that overlap with i.) promoter motifs and decrease promoter activity or impact translation initiation rate via ii.) changes in mRNA folding or due to iii.) changes in ribosomal binding site strength can result in expression changes. Furthermore, as our previous computational analysis highlighted the presence of unassigned serine TCA and TCG codons in Ec_Syn61Δ3’s genome^2^, which are known to arrest translation^51^, iv.) we sought to comprehensively explore the presence of unassigned codons on recoded chromosomes.

**Figure 3.**
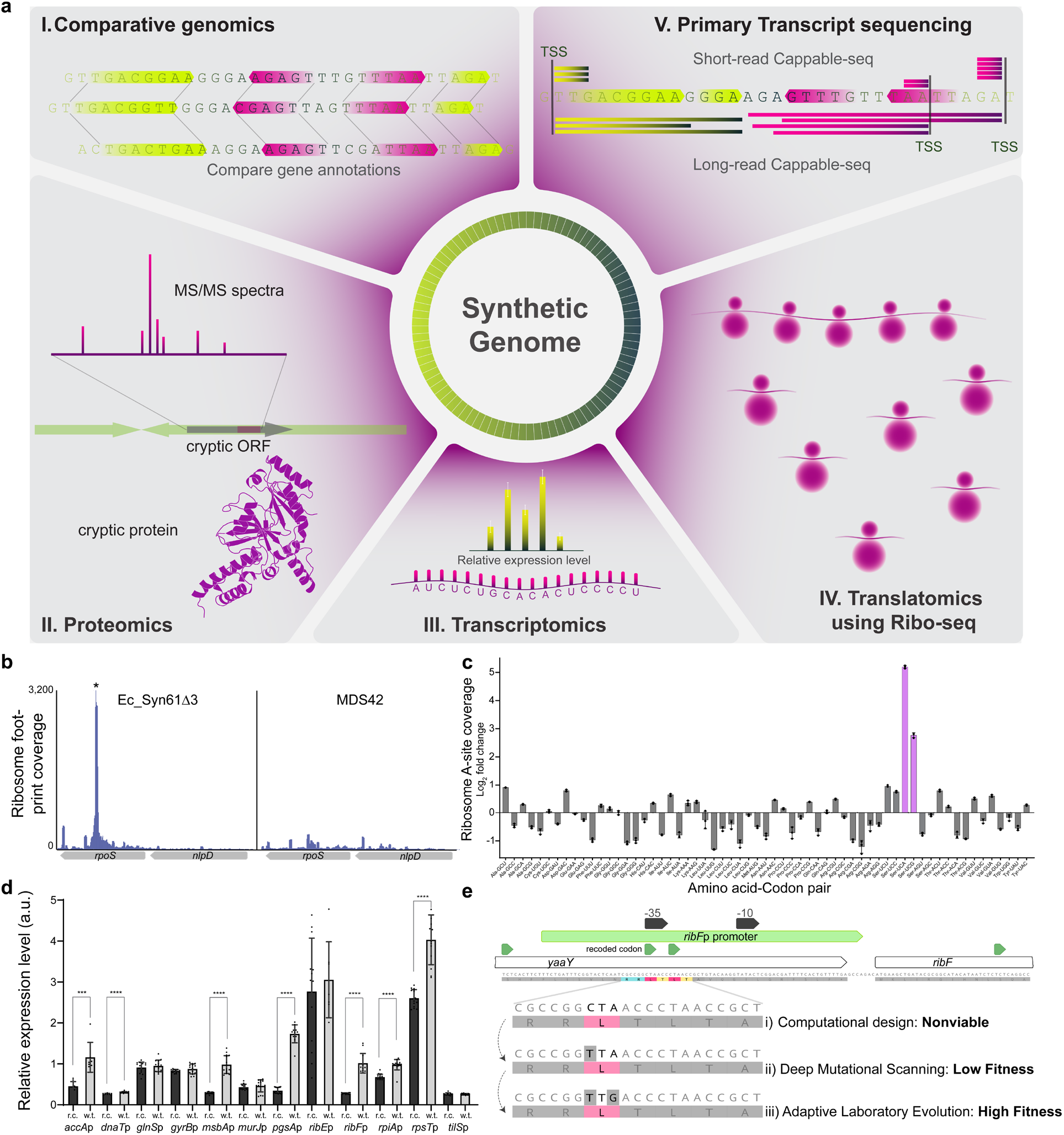
Multi-omics-based detection and troubleshooting of genome design issues. (**a**) Overview of multi-omics analyses utilized in this study to identify fitness-impacting changes on synthetic genomes. (**b**) Ribosomes stall at forbidden codons present in Ec_Syn61Δ3. The figure shows ribosome footprint coverage based on Ribo-seq along *rpoS* containing a TCA codon (marked with *) in Ec_Syn61Δ3 lacking tRNA^Ser(UGA)^ and tRNA^Ser(CGA)^ needed to translate TCA and TCG codons, respectively, and in *E. coli* MDS42 bearing the canonical genetic code. (**c**) Ribosome A-(aminoacyl)-site coverage of sense codons in Ec_Syn61Δ3. Ribosomes in the compressed genetic code accumulate at TCA (mRNA UCA) and TCG (mRNA UCG) codons—marked in magenta—present due to genome annotation errors and hypermutagenesis-induced mutations. Ribo-Seq data was collected in *n*=3 independent replicates, error bars indicate standard deviation. (**d**) Synonymous recoding frequently reduces promoter activity. Figure shows the mRNA output of intragenic promoters that overlap with synonymous codon swaps in Ec_Syn57. r.c. marks recoded promoter variant from Ec_Syn57, while w.t. marks the parental promoter variant. Experiments were performed in *n*=10 replicates. Error bars denote standard deviation; *** indicates a P value ≤ 0.001, while **** indicates P ≤ 0.0001 based on unpaired two-sided Student’s t-test. (**e**) Troubleshooting synonymous recoding-induced lethality using multiplexed genome editing and laboratory evolution. The nonviable computational genome design (i) was troubleshot using a combination of (ii) targeted DIvERGE mutagenesis followed by (iii) ALE to identify optimal variants within the growth-essential promoter of the *ribF-ispH* operon.

Unassigned and, therefore, forbidden codons can originate from annotation errors on bacterial chromosomes and—following the elimination of tRNAs or release factor recognizing these codons—can impact growth by arresting protein synthesis due to ribosome stalling^51^. Therefore, we first revised the gene annotations of our 57-codon genome using the most recent genome of *E. coli* K-12 MG1655 (GenBank accession number U00096.3), the parental strain of the genome-reduced MDS42 strain^37^. Genome reannotation revealed the presence of 109 new protein-coding genes and an additional 100 genes with altered coordinates, containing 644 forbidden codons. Performing the same analysis on Ec_Syn61’s genome indicated the presence of 105 new protein-coding genes and 104 genes with altered coordinates containing 186 forbidden codons. We also performed the same analysis on its evolved derivative, Ec_Syn61Δ3, indicating the presence of 217 unassigned forbidden codons. The large number of changes is not surprising as both Ec_Syn57’s and Ec_Syn61’s design are based on MDS42’s genome, (GenBank accession number AP012306.1) that, since its publication on December 3, 2011, remained essentially unchanged except for a minor annotation update during the construction of Ec_Syn61^14^. In the same timeframe, however, the genome of MG1655 underwent multiple updates, resulting in 79 newly annotated genes and 104 genes with altered coordinates^52^. Genome annotation updates also affect the number of forbidden codons in *E. coli* C321.ΔA, the first synonymously recoded organism, whose genome harbors eight unassigned TAG stop codons. Finally, the 17% increase in the number of forbidden codons between Ec_Syn61 and its Δ3 variant is due to the untargeted hypermutagenesis-based troubleshooting^3^ used to construct the fully recoded strain, indicating that off-target mutations during troubleshooting can increase the number of forbidden codons.

We next explored the consequences of unassigned codons by performing genome-wide ribosome occupancy analysis using ribosome profiling (Ribo-seq) on MDS42 and Ec_Syn61Δ3. As Ec_Syn61Δ3 lacks tRNA^Ser(UGA)^ and tRNA^Ser(CGA)^ needed to translate TCA and TCG codons, stalled ribosomes are expected to accumulate at these codons. These translational arrests would manifest as peaks centered around the forbidden codon in Ribo-seq data. In line with this hypothesis, we detected ribosome A-(aminoacyl)-site footprint peaks centered at forbidden codons in translated genes (**Figure 3b, Supplementary Figure 4**), and the analysis of ribosome occupancy across the genetic code table highlighted the arrest of ribosomes at the TCA and TCG serine codons (**Figure 3c, Supplementary Figure 5**). Based on the detected adverse effect of unassigned codons on bacterial translation, we selected all protein-coding genes that are actively transcribed in MDS42 and recoded these 73 genes containing 311 forbidden codons to match Ec_Syn57’s genetic code (**Supplementary Data 1**).

Due to the extent of genome annotation-induced recoding errors and recent reports indicating the presence of multiple cryptic translated open reading frames (ORFs) on the genome of *E. coli* MG1655^53–57^, we next performed a proteomics-based screen to identify highly expressed cryptic translated sequences in the genome of *E. coli* MDS42. We hypothesized that a high-resolution proteomics-based screen would allow us to detect abundant cryptic proteins, identify previously unannotated target codons for recoding, and eliminate these codons from the final recoded genome. The identification of these cryptic proteins is especially important as these translated ORF, if they exist and are sufficiently highly expressed, will induce fitness-decreasing ribosome stalling—as we showed earlier in the case of Ec_Syn61Δ3 (**Figure 3b-c**). To detect translated cryptic ORFs, we followed a three-step workflow. We first computationally identified all potential ORFs on the genome, including ORFs with alternative start codons^58^. Next, we performed deep proteomics and identified peptides matching cryptic ORFs but not known genes on the reference genome. Proteomic analysis detected translation from dozens of unannotated ORFs in both strains (**Supplementary Data 8**). For example, we detected translation from an alternative CTG start codon^58^ in antisense orientation within *lplA*, resulting in the extension of a 169 amino acid-long cryptic product and two unassigned TTG leucine, an AGC serine, and a TAG stop codon (**Supplementary Figure 6a**). Our screen revealed the presence of 48 novel translated sequences on the parental strain’s genome, containing 263 forbidden codons not conforming to the 57-codon genetic code of Ec_Syn57. The same analysis on Ec_Syn61 revealed 46 novel translated sequences containing 128 potential forbidden codons in the 61-codon genetic code of Ec_Syn61 (**Supplementary Figure 7**). In line with prior studies^53–57^, detected cryptic peptides mainly originated from intragenic ORFs and from noncanonical start codons upstream of known translational start sites (**Supplementary Data 8**). Finally, using independent Ribo-seq data, we also confirmed ribosome coverage at the *lplA*-derived cryptic ORF (**Supplementary Figure 6b**). In future experiments, we aim to more accurately map cryptic start codons and translated ORF using the combination of start-codon-arrested Ribo-seq^53^ and deep proteomics screens. Genomic, translatomic, and proteomic experiments jointly highlighted that existing recoded organisms harbor and tolerate hundreds of forbidden codons, and these codons might be responsible for the observed fitness decrease of previous recoded strains by sequestering ribosomes and interfering with global translation.

Next, we explored the effects of synonymous recoding on translation initiation. Although our genome design aimed to minimize changes in mRNA folding around the ribosomal binding site (RBS) by rational codon selection and by preserving the upstream 15-20 base pairs of the 5’ untranslated region (UTR) for overlapping genes (**Figure 1b**), the *in vivo* effects of our design on translation remained unexplored. Therefore, to explore potential changes in gene expression due to perturbations in translation initiation rates (TIR), we computationally predicted TIR for every protein-coding gene on the 57-codon genome and compared them to the TIR of the parental gene in MDS42. Our analysis indicated 816 genes with reduced and 507 genes with increased translation initiation rates (reaching at least a 2-fold TIR difference), representing 36.3% of all protein-coding genes of the host (**Supplementary Figure 9**). Comparative TIR analysis also revealed multiple potentially lethal design errors. In Segment 49, due to the erroneous recoding of the growth-essential *tadA*’s start codon, encoding tRNA adenosine-deaminase whose genomic coordinates were updated during our genome reannotation, our analysis predicted no expression for *tadA*. In line with this hypothesis, eliminating Segment 49’s parental copy resulted in drastic fitness loss and prevented segment integration. By reverting the accidentally introduced CTA start codon to the canonical ATG start codon via ssDNA-mediated recombineering, and this restoring gene expression^58^, we could quickly restore the function and integratability of this segment. In sum, the prediction of translational effects of synonymous recoding indicated 1,323 genes for troubleshooting.

Finally, we sought to comprehensively explore the effect of synonymous recoding on promoters across our synthetic genome. Using deep primary transcript sequencing, we first identified all active promoters of MDS42 in lysogeny broth, a commonly used rich bacterial medium. We performed Cappable-seq, a transcriptomic method that directly captures the 5’ end of primary transcripts, and determined transcription start sites (TSS) at single base resolution. Consequently, the location of TSS allows us to identify the location of all promoters on the analyzed genome^59,60^. Ultradeep Illumina short-read Cappable-seq analysis identified 39,140 TSS, from which 17,504 resided in protein-coding genes (*i.e.*, intragenic TSS). We also identified the principal promoters for all operons on MDS42’s genome using long-read Pacific Biosciences SMRT-Cappable seq^60^. This latter method extends the Cappable-seq workflow by performing long-read sequencing on entire primary transcripts, allowing the identification of all genes driven by a single promoter. Using Cappable-seq data, we identified promoter regions that overlap with codon changes on the recoded genome and explored the effects of synonymous recoding on a subset of these promoters.

Next, we individually synthesized 12 promoters in their wild-type and 57-codon recoded form and cloned them into a single-copy BAC vector. We then analyzed the mRNA output of these Cappable-seq-identified promoter variants using transcriptomics. Transcriptomic profiling revealed that synonymous recoding significantly decreased the activity of 7 out of the 12 assayed promoter pairs (**Figure 3d**). Notably, promoters that drive growth-essential genes and conferred decreased mRNA output (*i.e.*, *accA*p, *dnaT*p, *msbA*p, *pgsA*p, *ribF*p, *rpsT*p) were all located in recoded chromosomal regions that had previously failed to replace their corresponding parental copy, further confirming these promoters’ impact on the viability of recoded chromosomal regions. Based on this observation, we validated promoter mutations’ *in vivo* fitness effects using Segment 21’s *fabH-ycfH* region and attempted the multiplexed troubleshooting of the Cappable-seq predicted promoter issues. The deletion of Segment 21’s parental copy induced a drastic fitness drop, *i.e.*, a doubling time of 253 minutes, compared to the 35-minute doubling time of MDS42 Δ*recA* (**Supplementary Table 3**). Our Cappable-seq-based promoter analysis predicted the intragenic promoters of the growth-essential *fabH-acpP* and the *tmk-holB* operons as the cause of fitness drop due to the presence of codon swaps within these promoters (**Supplementary Figure 10**). We attempted to restore the expression of these operons by performing MAGE^41,61^ to insert a library of constitutive promoters in front of each target gene. Following three editing cycles with a mixture of 24 oligonucleotides targeting all three loci simultaneously, followed by growth-based selection, we identified fast-growing variants. Increased fitness required the insertion of three promoters, resulting in a doubling time of 83 minutes (**Supplementary Table 3**). The fitness benefit of newly inserted promoters in front of these growth-essential genes validated our transcriptomics-based predictions and suggested that the multiplexed troubleshooting of promoter issues is feasible during SynOMICS cycles.

Using genome, transcriptome, translatome, proteome profiling, and computational analyses, we comprehensively identified potential design errors and fitness-decreasing synonymous codon changes on existing recoded genomes and validated these using multiplexed genome editing.

### Troubleshooting design errors

We initiated troubleshooting the synthetic 57-codon genome based on the list of multi-omics-identified design errors using multiplexed genome editing and adaptive laboratory evolution. We first focused on troubleshooting design issues that we identified using multi-omics screens (**Figure 3a**). Using the list of recoding-impacted promoters and RBSes, these regulatory elements’ expression and translation level, respectively, and the operon structure of *E. coli* MDS42 based on Cappable-seq, we troubleshot predicted issues using three strategies: I.) in Segments 4 and 7, we utilized CRISPR/Cas9 assisted MAGE to simultaneously insert a promoter and an optimized RBS in front of impacted operons before attempting to delete the segment’s parental copy. As expected, promoter insertions restored viability. II.) In the case of Segments 5, 22, 61, 64, and 86, in which multiple distant changes were necessary, we resynthesized the entire recoded segment with a modified design that corrected all potential errors (for a detailed description of modifications made to troubleshoot each segment, see Supplementary Methods). In Segment 64, resynthesis also allowed us to correct a design issue leading to the separation of a Cappable-seq-discovered promoter region and the corresponding growth-essential *rpsM-rplQ* operon in the neighboring Segment 63. Segment resynthesis restored the viability of all but one segment (*i.e.*, Segment 64, which we later troubleshot using a combination of all three methods).

III.) For all remaining design issues that induced fitness defects but did not prevent the deletion of the parental copy, we developed a parallelizable, multiplexed approach. We took advantage of the fitness-based selection pressure posed by deleting the parental chromosomal region in SynOMICS and performed targeted mutagenesis to repair errors and increase fitness. We hypothesized that targeted mutagenesis would allow us to identify beneficial mutation combinations at multiple loci simultaneously while minimizing off-target effects. We utilized DIvERGE, an ssDNA-recombineering-based targeted mutagenesis method that allows an up to million-fold increase in mutation rate along the entire lengths of multiple predefined loci—ranging from promoters up to entire operons—without inducing off-target mutations, and capable of targeting hundreds of sites simultaneously^62,63^. We synthesized 324 DIvERGE oligonucleotides targeting promoter issues together with an additional 442 MAGE oligonucleotides to correct 447 DNA synthesis errors. Next, using a pooled mixture of all DIvERGE and MAGE oligos targeting all potential promoter issues and DNA synthesis errors in each partially recoded strain, we performed one to seven rounds of diversification followed by growth-based selection. Whole genome sequencing of the resulted clones indicated the reversion of multiple DNA synthesis errors, up to ten in a single strain, and newly emerged promoter mutations within DIvERGE oligo targets (Supplementary Data 1 and 7). In Segment 0, where promoter transcriptomics identified the recoded promoter of the *ribF-ispH* operon in the upstream *yaaY* as the cause of nonviability (**Figure 3d**), mutagenesis identified a single point mutation. In *yaaY*, two neighboring recoded leucine codons overlap with the −35 region of the growth-essential *ribF-ispH* promoter and DIvERGE mutagenesis resulted in the reversion of one of the recoded leucine codons restoring viability (**Figure 3e**). Similarly, diversifying the damaged promoter of the growth-essential *msbA-ycaQ* operon in Segment 18 and the *pgk-fbaA* operon in Segment 56 identified beneficial mutations (**Supplementary Figure 11a-b**).

Next, following targeted editing and using the fastest-growing variants from each screen, we initiated Adaptive Laboratory Evolution (ALE) experiments. ALE is a powerful method that allows the discovery of compensatory mutations in a nontargeted manner and the identification of design-issues that remained hidden in our previous analyses. To perform ALE, we relied on natural mutagenesis and growth-based selection using a large population size (5×10^9^ cells per transfer and a total population size of approximately 10^12^ cells per ALE step) without inducing hypermutagenesis. Based on prior studies^64,65^, we hypothesized that relying on the natural mutation rate of *E. coli*—approximately 10^-3^ mutations per genome per generation^66,67^—and using a population size of ∼10^12^ cells would allow us to explore all possible single-step mutations across the entire genome at every ALE step and thus identify rare beneficial mutations. This approach is also expected to minimize the accumulation of fitness-decreasing hitchhiker mutations, including frameshifts leading to forbidden codons. In line with this hypothesis, strain-fitness rapidly improved during ALE experiments. Within only 140–450 generations performed in 16–50 days, we identified improved variants (**Figure 3e**). As expected, based on the low mutation rate applied during evolution, whole genome sequencing of the ALE-evolved variants indicated the accumulation of only a few mutations within the recoded region, ranging from one to six across evolved strains (Supplementary Data 7). The only exception was the strain carrying Segments 82–0, which accumulated 26 mutations within the 283 kbp recoded region, likely due to the emergence of a variant with an elevated mutation rate during ALE.

Using SynOMICS, we replaced the parental copy of all synthetic segments, up to twelve in a single genome. During SynOMICS cycles, we iteratively applied rational genome editing and ALE to maintain the viability of the partially recoded genome while deleting the parental copy and integrating the corresponding recoded segment. In total, we performed 89 SynOMICS integration rounds, 113 genome editing cycles, and 680 cumulative days of ALE to construct and troubleshoot partially recoded strains (Supplementary Methods). Iterative genome editing and troubleshooting during SynOMICS cycles in eleven *E. coli* strains overcame lethal fitness issues, restored the integrability of all tested segments, and allowed us to complete the 57-codon synthetic genome.

### Genome assembly

Following the assembly and testing of the 3.98 Mbp genome in eleven sections, we utilized an extended version of SynOMICS to move and merge the troubleshot recoded chromosomal regions into a single strain. We merged chromosomal regions by reversing the integration step of SynOMICS using a modified version of our pINTsg plasmid termed pFISSIONsg, which liberated the entire recoded chromosomal region from the parental genome. The liberated linear chromosomal region (150–451 kbp in size, containing 4–12 segments) could then be transposed into the empty BAC within the same cells using Lambda-Red recombineering based on matching terminal homologies (**Figure 4a**). Simultaneously, the delivery of an antibiotic marker cassette, with flanking genomic homologies to seal the genomic Cas9-cut, allowed us to select clones with the expected genome topology based on antibiotic resistance profile. The combination of our previously optimized SynOMICS plasmid system with pFISSIONsg resulted in frequent chromosome fission on all tested strains (Supplementary Methods), usually approaching 100% efficiency and generating thousands of clones. Following chromosome fission, we utilized colony PCR to confirm the genotype of selected clones, followed by Illumina and Oxford Nanopore whole-genome sequencing and *de novo* genome assembly to ensure the sequence and topology of the fissioned genome. Once error-free clones are identified, we electroporated or conjugated the 160–460 kbp fission BACs into a recipient cell containing a partially recoded genome. Following the delivery of the fission BAC, the standard workflow of SynOMICS without any additional modifications allowed us to combine recoded regions. We accelerated the delivery of the fissioned chromosomes by generating a version of the F plasmid that does not share replication origin (ORI) with the commonly used F-plasmid-based BAC vectors, including our fission BACs. This is because, in our experiments, the competing replication origins of the two BACs frequently resulted in slow growth and prevented the conjugation of the fissioned genome into our recipient strain. We overcame this issue by replacing the F plasmid ORI with the low-copy-number ORI of the RK2 plasmid^68^, thereby preventing plasmid incompatibility and restoring conjugatability.

**Figure 4.**
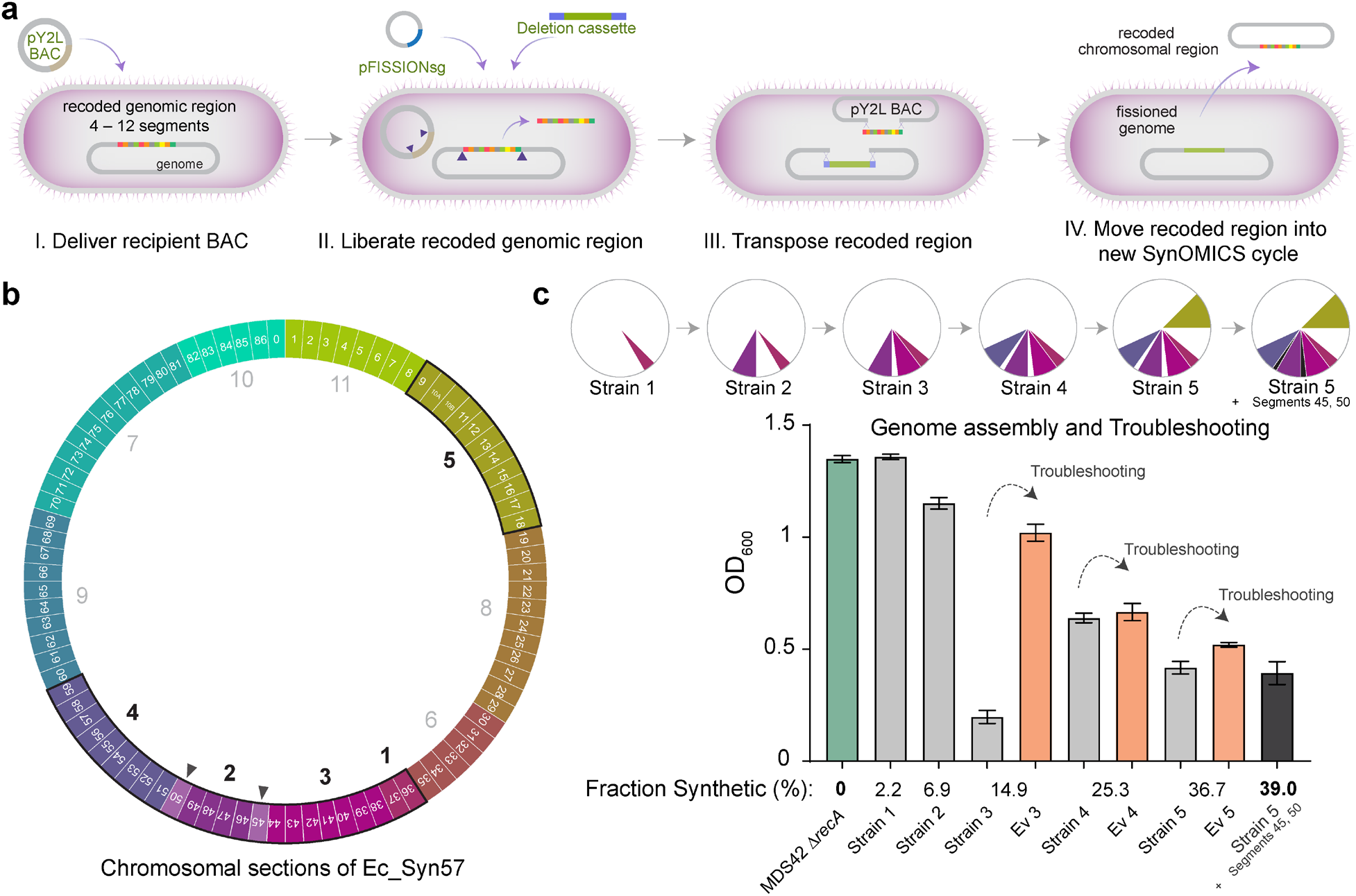
Assembly of recoded chromosomal regions to create Ec_Syn57. (**a**) The reversal of SynOMICS’s steps and its direction allows genome fission, fusion, and assembly from separately constructed synthetic chromosomal regions. Chromosome fission is achieved by delivering recipient BAC (*i.e.*, the pYES2L fission BAC, abbreviated as pY2L) into a cell line bearing a partially recoded chromosome (**I.**), and (**II.**) following the expression of Lambda-Red and Cas9 from pRedCas2 (not shown), chromosome fission is initiated by delivering pFISSIONsg—a four-plex nonrepetitive sgRNA expression plasmid—and a version of the SynOMICS deletion cassette that only contains terminal genomic homologies. (**III.**) CRISPR/Cas9-cuts liberate the recoded chromosomal region from the genome and linearize the recipient BAC allowing the Lambda-Red-mediated terminal homology-directed transposition of the recoded chromosomal region into the recipient BAC while simultaneously sealing the genomic cut. Finally, (**IV.**) the fissioned chromosomal region is delivered into a new recipient cell where it is integrated using the standard SynOMICS workflow, depicted on Figure 2c. (**b**) Construction of the synthetic genome of Ec_Syn57 from 11 simultaneously constructed synthetic sections. Numbers indicate the steps in which genomic sections are merged to assemble the final, fully synthetic genome of Ec_Syn57. Segments 45 and 50 (marked as ▲) were added following the merging of sections 1 to 5. To date, sections 1–5 have been combined, as indicated on Figure 4c, yielding 7 strains containing the synthetic genome of Ec_Syn57. (**c**) SynOMICS-based sequential assembly and troubleshooting of recoded chromosomal sections to generate Ec_Syn57. Following the construction of recoded chromosomal regions, we utilized SynOMICS (Figure 4a) to transfer and integrate recoded chromosomal regions in *E. coli* MDS42 Δ*recA* containing Segments 36 and 37. Following the assembly of three chromosomal regions, we initiated genome-editing- and ALE-based troubleshooting to increase the fitness of partially recoded strains before the next chromosomal region was transferred. Pie chart displays the steps of genome assembly, colored sections mark synthetic recoded chromosomal regions transferred to obtain *E. coli* MDS42 Δ*recA* containing Segments 9–18 and 36–59. For the detailed description of assembly steps, see Supplementary Methods. Bar graph shows final optical density at 600 nm (OD600) following aerobic growth in 2×YT broth, a rich bacterial growth medium, at 37 °C. Source data is available in this paper.

SynOMICS-based genome synthesis allowed us to assemble recoded chromosomal regions from distinct donor strains sequentially (**Figure 4b**). Despite transferring recoded chromosomal segments to a new genomic background and thus eliminating potential compensatory mutations from previous troubleshooting steps on the genomic background, recoded regions remained viable in the recipient strain. We observed a strong correlation between the extent of genome recoding and fitness. Strains with increasingly recoded genomes displayed lower overall fitness, quantified as the maximal attainable optical density in rich bacterial growth medium (*i.e.*, OD at 600 nm, measured in 2×YT broth; **Figure 4c, Supplementary Table 3**). Therefore, following the assembly of multiple chromosomal regions, we performed ALE-based troubleshooting and identified variants with increased fitness. Iterative genome replacement and troubleshooting using SynOMICS allowed us to sequentially combine 35 segments from five donor strains, representing 1.55 Mb, 39% of the entire genome. During troubleshooting cycles, we observed only 38 ALE-derived mutations in the recoded region of this strain, demonstrating the benefits of using rational genome editing and ALE with low mutation rate to troubleshoot synthetic genomes. As a reference, the hypermutagenesis-based troubleshooting of Ec_Syn61Δ3 in an earlier study, which utilized a mutation rate approximately 8,000 times higher than our ALE experiments, led to the accumulation of 482 off-target mutations^3^.

Using iterative genome synthesis and troubleshooting, in 1120 days of ALE-based troubleshooting and 210 genome editing cycles, we successfully constructed the complete Ec_Syn57 genome across seven strains and consolidated five synthetic chromosomal segments into a single strain harboring 39% of the entire genome (**Figure 4b**). We hypothesized that the resulted collection of partially recoded strains together with currently available recoded organisms could serve as a unique opportunity to comprehensively explore the consequences of large-scale synonymous recoding and compare these effects across synthetic and parental genomes.

### Multi-omics-based discovery of recoding effects

Finally, we explored the transcriptome, translatome, and proteome-level consequences of synonymous recoding by performing simultaneous genome-transcriptome-translatome profiling on a wide array of recoded strains. We expected that exploring these effects along the steps of the central dogma would comprehensively reveal the consequences of synonymous recoding, highlight targets for future troubleshooting, and potentially uncover previously unseen effects. Therefore, we characterized changes in mRNA synthesis, promoter distribution, ribosomal translation, and protein synthesis by first performing Illumina and Nanopore whole genome sequencing and *de novo* genome assembly on all strains to obtain accurate genomic information and then performed RNA-seq, RIBO-seq, and Cappable-seq analysis on recoded strains and their corresponding wild-type parents.

Comparative transcriptome profiling using RNA-seq revealed large-scale changes in all recoded strains compared to their parental strain. Strains harboring the replacement of 33%, 6.9%, 11.7% of their genome with the 57-codon version contained 1156, 710, and 1220 differentially expressed genes (*i.e.*, showing at least two-fold relative transcript level difference and P<0.05, based on *n*=3), respectively. The comparison of Ec_Syn61Δ3’s transcriptome to MDS42, in the same manner, indicated 1366 differentially expressed genes (**Figure 5a**).

**Figure 5.**
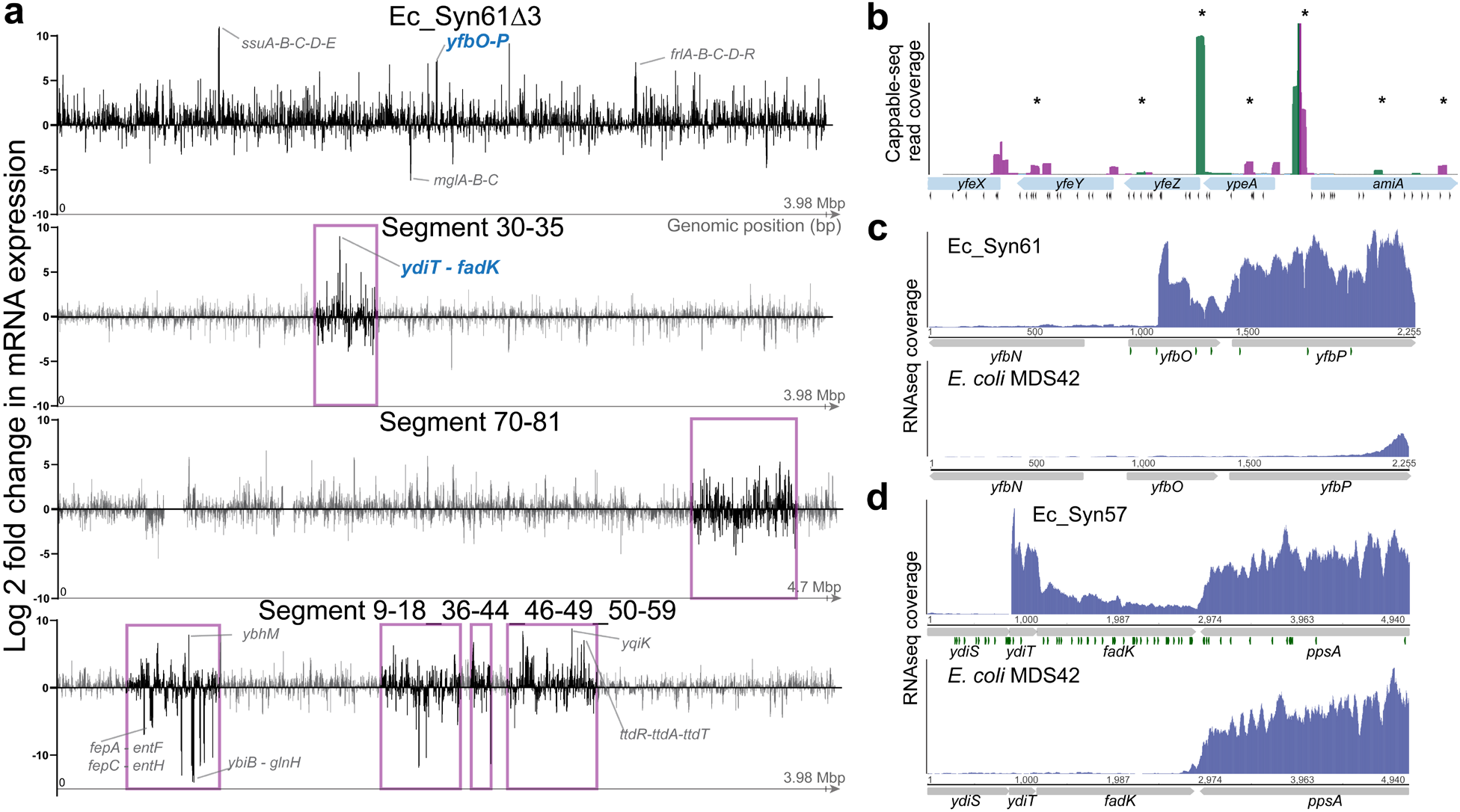
Synonymous recoding induces widespread changes in gene expression. (**a**) The recoded genome of Ec_Syn61Δ3 and recoded chromosomal regions of Ec_Syn57 display marked differences in mRNA expression. The figure shows differential mRNA expression for all protein-coding genes along the recoded genome of Ec_Syn61Δ3 and partially recoded genomes of Ec_Syn57, compared to their parental variant. Recoded regions on partially recoded genomes are marked in magenta. Example operons displaying the most significant changes in mRNA expression are indicated. Bars indicate mean differential expression. Differential expression values were calculated using the EdgeR algorithm using three independent replicates (*n*=3) Source data is provided in Supplementary Data 3. (**b**) Synonymous recoding induces the widespread appearance of cryptic promoters. Figure shows the position and direction of primary transcripts based on primary transcriptome sequencing using Cappable-seq. Recoding-induced novel transcriptional start sites are marked with a star (*). Cappable-seq reads in magenta marks reverse transcript orientation, while green marks forward transcript orientation. Black triangles indicate recoded codons. Cappable-seq experiments are based on *n*=2 independent replicates. For the analysis of the same locus in the parental MDS42, see **Supplementary** Figure 14. (**c-d**) Synonymous recoding-induced promoters behind transcriptome changes of Ec_Syn61Δ3 and Ec_Syn57, highlighted in Figure 5a. The figure shows RNA-seq coverage within the *ybfO-P* locus of Ec_Syn61Δ3 (**c**) and the *ydiT-fadK* locus in Ec_Syn57 (**d**) and their corresponding parental copy. Green triangles indicate recoded codons. RNA-seq experiments were performed in *n*=3 independent replicates.

Comparative translatome profiling using Ribo-seq confirmed the extent of transcriptomics-indicated effects. Ribosome profiling indicated 1133, 638, and 719 differentially translated genes (*i.e.*, showing at least two-fold relative ribosome occupancy difference and P<0.05, based on *n*=3) on the same set of genomes, respectively. The comparison of Ec_Syn61Δ3’s translatome to MDS42 indicated 1148 differentially translated genes (**Supplementary Figure 12**). Based on Ribo-seq results, we selected all genes in recoded regions that display significant changes in translation rate compared to their parental copy. In follow-up troubleshooting steps, we will utilize our multiplexed MAGE- and DIvERGE-based troubleshooting workflow (**Figure 2c**) to troubleshoot gene expression and identify optimal alleles in a massively parallel manner.

Next, we investigated the correlation between our previously predicted gene expression changes and the ribosome profiling-based gene expression results to aid genome troubleshooting. The availability of RNA- and Ribo-seq data from the same samples allowed us to measure changes in relative translation efficiency for every gene between their parental and recoded variant and compare the resulting dataset to our previous computational TIR prediction. Unexpectedly, *in vivo* translation efficiency differences between the recoded and parental genome showed no correlation with *in silico* predictions (R^2^=4.86x10^−3^, **Supplementary Figure 13**).

As translation efficiency measurements assess the correlation between the number of mRNA reads and ribosome footprints on a single gene, we hypothesized that promoter location and activity changes could influence translation efficiency results and account for the lack of correlation with *in silico* results. According to this hypothesis, newly generated intragenic promoters could generate artifacts in transcriptomic data by increasing transcript levels without upregulating translation, while antisense transcripts could downregulate translation through dsRNA formation, increased mRNA degradation, or the inaccessibility of RBS sequences^69,70^. We tested this hypothesis by repeating Cappable-seq on our partially recoded 57-codon strain carrying Segments 9–18, 36–44, 46–49, and 51–59, *i.e.*, 36.7% of the entire recoded chromosome. Next, we analyzed the location and expression intensity of transcriptional start sites (TSS) and compared the parental and recoded genome. In line with our expectation that synonymous recoding only impacts intragenic promoters, from intergenic TSS we detected no difference in total mRNA output between the parental and recoded genomic variant (that is, a total mRNA output of 303,477 transcripts per million (TPM) in the case of the recoded strain, compared to 292,890 TPM in the case of the parental MDS42). In contrast to intergenic TSS, intragenic TSS were markedly affected. The total sense transcript output from intragenic TSS—that are producing transcripts with matching direction of the gene they reside in—increased 2.23-fold in the recoded strain, while the transcript output from antisense intragenic TSS increased 4.01-fold compared to the antisense transcription of the parental MDS42. Most of these recoding-induced antisense promoters’ mRNA output was comparable to sense transcription for the same gene (**Figure 5b, Supplementary Figure 14**). Due to the widespread presence of these antisense promoters in recoded genes—exceeding one newly generated antisense TSS per gene—and as antisense transcription is known to downregulate gene expression by affecting mRNA stability and ribosome binding^69–71^, we hypothesize that recoding-induced antisense transcription might be responsible for a significant fraction of synonymous-recoding-induced fitness defects, jointly with direct perturbations in natural intragenic promoters (**Figure 3d**)

Consequently, we quantified the extent of antisense transcription in four isogenic *E. coli* strains with three distinct genetic codes. We selected MDS42 as wild-type parental, Ec_Syn61Δ3 as a 61-codon, and two strains bearing distinct 57-codon recoded chromosomal regions of Ec_Syn57.

These two latter strains harbored 33% and 6.9% of the synthetic 57-codon genome. Strand-specific (stranded) RNA-seq and subsequent identification of sense and antisense transcripts allowed us to quantify antisense transcription in each analyzed strain. Stranded RNA-seq results confirmed our previous Cappable-seq-based results. Chromosomal regions of Ec_Syn57 displayed 604% and 715% of the antisense transcription of their parental copy, respectively. In comparison, the 61-codon code of Ec_Syn61Δ3 resulted in 301% of the antisense transcription of the parental MDS42 (**Supplementary Figure 15**). Our RNA- and Ribo-seq experiments further supported the negative impact of newly generated promoters. Recoding-induced promoters frequently resulted in drastic expression changes in neighboring genes in both Ec_Syn61 and Ec_Syn57 (**Figure 5a, c-d**), indicating the severe impact of these promoters on gene expression. As our analysis revealed a correlation between the number of recoded codons and the extent of antisense transcription, these results suggest that creating genomes with highly compressed genetic codes requires the active elimination of *de novo* promoters during genome design.

Using multi-omics analyses on multiple recoded bacterial strains, we discovered that synonymous codon replacements perturb noncoding sequence motifs responsible for gene expression and induce transcriptional and translational noise, leading to drastic transcriptome and proteome perturbation. Our analysis revealed that altering the natural codon choice results in the widespread emergence of new intragenic promoters, inducing transcriptional and translational noise that might significantly contribute to the observed fitness effects of synonymous recoding. In the follow-up six steps of genome construction (**Figure 4b**), we focus on troubleshooting gene expression changes indicated by our analyses.

## Discussion

We have explored the transcriptional, translational, proteomic, and phenotypic effects of synonymous genome recoding using an array of *E. coli* strains bearing synthetic genomes with modified genetic codes, and based on these results, constructed a viable 4 Mbp genome of *E. coli* that relies on 57 genetic codons only. The construction of this 57-codon genome, which required 7.3×10^4^ edits—more than 4× what was previously attempted (**Supplementary Figure 8**)—demonstrates that both the genome and genetic code of living organisms are malleable and organismal genomes are designable for functions not available in current living systems.

The strategy we have developed for genome construction—the multi-omics-based identification of fitness-impacting changes on computationally designed genomes and our scalable, universal approach to troubleshoot these design errors—provides a general workflow for future prokaryotic and eukaryotic genome synthesis projects. Our validation of this workflow in hundreds of experiments demonstrates its robust performance in troubleshooting diverse design issues (Supplementary Methods).

We have shown that synonymous codon replacements induce widespread promoter perturbations in recoded organisms, resulting in frequent changes in gene expression and the generation of new promoters. As the elimination of seven codons from the canonical genetic code resulted in the appearance of ∼1 new promoter in each recoded gene, providing an up to 715% increase in global antisense transcription in a 57-codon genetic code and a 301% increase in a 61-codon code, compared to the parental wild-type variant, our results confirm a prior hypothesis^72,73^ and show that genomic codon preference is naturally evolved to reduce transcriptional noise.

Furthermore, we revealed that all previously constructed recoded organisms harbor non-recoded, forbidden codons due to frequent annotation errors in template genomes, leading to the presence of unassigned codons and, consequently, ribosome stalling at these codons (**Figure 3b-c**). As our profiling of translating ribosomes highlighted deleterious effects due to the widespread presence of hundreds of forbidden codons in a previously recoded synthetic *E. coli* genome^3^, we developed a generalizable genomics- and proteomics-based workflow for detecting and correcting these issues. We expect that these results will soon allow us to achieve, to our knowledge, the first 57-codon organisms and recoded genome with the lowest fraction, less than 1%, of unassigned codons (**Supplementary Figure 8**). Once complete, at an estimated cost of ∼$3.2 million (**Supplementary Table 4**), the resulting Ec_Syn57Δ7 strain will provide a broadly applicable and unique chassis for fundamental research and biotechnology.

We anticipate that our workflow of genome construction and the multi-omics data generated in this work will significantly reduce the cost and timescale of heavily modified genomes’ construction and troubleshooting. We estimate that at the current $0.13/bp synthesis cost of clonal dsDNA, the resynthesis of Ec_Syn57Δ7 would cost ∼$800,000 (**Supplementary Table 4**).

In Box 1, we conclude our observations for future genome designs. Follow-up works might translate and validate our results in ongoing prokaryotic and eukaryotic genome recoding _projects_18,19,21,26,29,42,74–76.

We expect that our combination of genome-transcriptome-translatome-proteome co-profiling, whole genome synthesis, genome editing, and directed-evolution-based troubleshooting will provide a universal and cost-effective strategy to construct synthetic genomes with customized properties, including genetic codes and functionalities not found in nature^34^.

#### Box 1 Guidelines for synthetic genomes

##### Genome design

- As genomic annotations errors and cryptic translated sequences yield unassigned codons, proteomics- and ribosome-profiling-based discovery of translated sequences^53,56^ is necessary before genome and genetic code refactoring.
- Before genome design, accurately identifying promoters and operon structures is necessary using simultaneous short- and long-read Cappable-seq for prokaryotic and ReCappable-seq for eukaryotic genomes^77^.
- Following promoter identification, experimental and computational transcriptional factor binding site analyses can reveal essential regulatory sequences needed to be preserved during genome design^78–80^.

##### Genome synthesis, assembly, and troubleshooting

- Projects should minimize DNA synthesis errors and avoid using cloning hosts containing active mobile genetic elements or hypermutator alleles.
- We hypothesize that genome-transcriptome-translatome-proteome co-profiling during genome construction, potentially in combination with genomic machine learning and metabolic models^81–83^ to discover and prioritize fitness-decreasing errors for troubleshooting, and the multiplexed correction of these design issues using genome editing and directed evolution, will yield high-fitness variants.
- Finally, adaptive laboratory evolution of synthetic genomes can substantially improve fitness, paving the way for industrial use.

**Future genome design projects should focus on developing and implementing algorithms that reduce changes in promoter activity, regulatory functions, and translation rate and avoid generating unwanted cryptic promoters and translated ORFs**. The development of predictive models capable of forecasting transcriptional and translational effects and integrating these predictions to refactor codon composition is expected to minimize the negative fitness impact of synonymous genome recoding^83–92^.

## Supporting information

Supplementary Material - Synthetic genomes unveil the effects of synonymous recoding

## Acknowledgments

We thank György Pósfai (Biological Research Centre, Hungary) for sharing MDS42 and Jason W. Chin’s team (Medical Research Council Laboratory of Molecular Biology, UK) for sharing Syn61 and Syn61Δ3 via Addgene. Funding for this research was provided by the US Department of Energy (DOE) under grant DE-FG02-02ER63445, by the National Science Foundation (NSF) Award number: 2123243 (both to G.M.C.), by a JGI DNA Synthesis Program grant (award number 508712) (to G.M.C.), by the National Health Institute (NIH) Award number 1K99EB035165-01 (to A.N), and by the EMBO LTF 160-2019 Long-Term fellowship (to A.N). A.C.P. acknowledges funding from the Swiss National Science Foundation (Award number P2ELP2_181884 to A.C.P.). The authors thank GenScript USA Inc. for their DNA synthesis support, Silvia Tornaletti and Luciano Brocchieri at TB-Seq, Inc., San Fransisco, CA for their support with Ribo-seq and RNA-seq experiments and data analysis; Ranomics, Inc., Canada, for experimental and DNA synthesis support; and Dan Snyder, Katrina Harris, and all members of SeqCenter, LLC, Pittsburgh, PA for their support with DNA and RNA sequencing experiments. We thank Bo Yan from New England Biolabs, Ipswich, MA for support with Cappable-seq experiments; and Tan Chen, Yun Wang, Fei Huang, Xianping Deng, Jiahui Zhang at BGI Research for their help with DNA synthesis for the project. DNA synthesis for troubleshooting experiments in this work (proposal: https://doi.org/10.46936/10.25585/60008479) was also conducted by the U.S. Department of Energy Joint Genome Institute (https://ror.org/04xm1d337), a DOE Office of Science User Facility, supported by the Office of Science of the U.S. Department of Energy operated under Contract No. DE-AC02-05CH11231.

## Author contributions

A.N. developed the project, performed or directed all experiments and analyses, and wrote the manuscript with input from all authors. A.N. and G.M.C. supervised research. B.B. carried out MS/MS analyses. R.F. assisted in troubleshooting experiments, carried out adaptive laboratory evolution and growth rate measurements. A.C-P. removed transposons; computational analyses including genome reannotations and analysis of impacted start codons, promoters, RBSes, and RNA levels; obtaining and troubleshooting: RNA-seq, adaptive laboratory evolution, growth of strains; DNA for BAC genome sequencing; construction, troubleshooting, and screening of recoded genome segments; corrected segments; and contributed to funding acquisition. M.B.T., S.Y., J.A., N.O., A.R., D.C. assisted in experiments at the early stage of the project. S.Y., M.L., M.W., Q.Z., K.C., F.H., I.B., Y.Y., M.H.S., A.T., A.J., Z.L., Y.S., M.H. performed DNA synthesis. V.A. and O.S. performed media preparation and DNA extraction for whole-genome sequencing. B.K., M.S., and V.S. provided reagents for the project. B.H. assisted with data visualization.

## Data availability

Raw data from whole genome sequencing, transcriptome, Cappable-seq, and Ribo-seq experiments have been deposited to Sequence Read Archive (SRA) under BioProject ID PRJNAxxxxxxxx. All materials used in this study are available from the corresponding authors upon request. The annotated genome of Ec_Syn57 is available in the Supplementary Material of this paper. Mass spectrometry and proteomic datasets are available on MassIVE, under MSVxxxxxxxx. To aid future recoding efforts and the investigation of the effects described in this work, we are depositing the current set of 57-codon Ec_Syn57 genome regions on Addgene, making it freely available together with the datasets generated in this paper. Source data is provided in the Supplementary Material of this paper.

## Code availability

The comparison of protein-coding gene annotations between the original and updated genome-versions were performed using the compare_annotations.py script available from https://github.com/rrwick/Compare-annotations, developed by Ryan Wick, Peter Doherty Institute for Infection and Immunity of the University of Melbourne, Melbourne, Australia. Cappable-seq data analysis scripts and protocols are available on GitHub from https://github.com/elitaone/cappable-seq, developed by Bo Yan from New England Biolabs, Ipswich, MA, USA, and described in references ^59,77^. We utilized ProteoMapper version 5.2.0 ^93^, available from https://tppms.systemsbiology.net/pm, to analyze mass spectrometry results.

## Methods

### Bacterial media and reagents

Adaptive Laboratory Evolution experiments utilized Lysogeny Broth Lennox (LBL) supplemented with Tris/Tris-HCL as a buffer, prepared by dissolving 10 g/l tryptone, 5g/l yeast extract, 1.5 g/l Tris/Tris-HCL, and 5 g/l sodium chloride in deionized H2O and sterilized by autoclaving. Bacterial cell cultures and genome editing experiments utilized 2×YT media consisting of 16 g/l casein digest peptone, 10 g/l yeast extract, and 5 g/l sodium chloride. 2×YT agar plates were prepared by supplementing LBL medium or 2×YT with agar at 1.6% w/v before autoclaving. Super Optimal Broth (SOB) was prepared by dissolving 20 g/l tryptone, 5 g/l yeast extract, 0.5 g/l sodium chloride, 2.4 g/l magnesium sulfate, and 0.186 g/l potassium chloride in deionized H2O, and sterilized by autoclaving.

### Bacterial genome sequencing and annotation

Genomic DNA from overnight saturated cultures of isogenic bacterial clones was prepared using the MasterPure™ Complete DNA and RNA Purification Kit (Lucigen) according to the manufacturer’s guidelines and sequenced at SeqCenter (Pittsburgh, PA, USA). BACs containing individual recoded chromosomal regions were isolated using the ZR BAC DNA Miniprep Kit (Zymo Research) based on the manufacturer’s protocol. Sequencing libraries were prepared using the Illumina DNA Prep kit and IDT 10 bp UDI indices with a target insert size of 280 bp and sequenced on an Illumina NextSeq 2000 or on an Illumina NovaSeq X Plus, producing 150 bp paired-end reads. Demultiplexing, quality control, and adapter trimming were performed with bcl-convert (v4.2.4). Reads were then trimmed to Q28 using BBDuk from BBTools and aligned to their corresponding reference by using Bowtie2 2.3.0^94^ in --sensitive-local mode. Single-nucleotide polymorphisms (SNPs) and indels were called using breseq (version 0.36.1)^95^. Only variants with a prevalence higher than 75% were voted as mutations. Following variant calling, mutations were also manually inspected within the aligned sequencing reads using Geneious Prime® 2023.2.1. We performed *de novo* sequencing and hybrid genome assembly on all parental strains, strains with completed recoded chromosomal sections, and SynOMICS fissioned and fusioned, partially recoded strains. Hybrid genome assemblies were performed as described earlier^2^. Briefly, we generated 300–600 Mbp of Oxford Nanopore (ONT) long-reads by PCR-free library generation (Oxford Nanopore, UK) on a MinION Flow Cell (R9.4.1) and 3–15×10^6^ 150 bp paired-end reads on an Illumina NextSeq 2000 or on an Illumina NovaSeq X Plus. Quality control and adapter trimming were performed with bcl2fastq 2.20.0.445 and porechop 0.2.3_seqan2.1.1 for Illumina and ONT sequencing, respectively. Next, we performed hybrid assembly with Illumina and ONT reads using Unicycler 0.4.8 with the default parameters. Finally, the resulted single, circular contig representing the entire genome was manually inspected for errors in Geneious Prime® 2023.2.1. and annotated based on sequence homology using the BLAST function implemented in Geneious Prime® 2023.2.1. based on *E. coli* K-12 MG1655 (NCBI ID: U00096.3) as reference. Protein-coding gene annotations between the original and updated genome-versions were performed using the compare_annotations.py script available from https://github.com/rrwick/Compare-annotations (developed by Ryan Wick, University of Melbourne, Melbourne, Australia). Gene essentiality in rich bacterial medium was determined based on Ref^96^. We note that the previously reported^14^ 51 bp insertion between *mrcB* and *hemL* of *E. coli* MDS42, not reported in the original AP012306.1 GenBank sequence, was confirmed by our analysis.

### Segment DNA synthesis and assembly

Chromosomal segments of Ec_Syn57 were synthesized and assembled into BAC vectors by BGI Research (Shenzhen, China) and GenScript USA Inc. (Piscataway, USA and Nanjing, China). Segments 50, 60, 62, 63, 65, 66, 67, 68, and 75 were synthesized and assembled by BGI Research, while Segments 2, 5, 6, 8, 10A, 10B, 13, 19, 22, 35, 37, 45, 53, 55, 57, 61, 64, 73, 86, and 89 were synthesized by GenScript USA Inc. Segment synthesis at BGI Research utilized 55-80 nucleotide-long oligonucleotides as building blocks that shared 20 nucleotide terminal homology, synthesized using the standard phosphoramidite DNA synthesis procedure^97^. These 55-80-nt ssDNA oligonucleotides were subsequently assembled into 600 bp dsDNA fragments using polymerase cycling assembly (PCA). Following agarose gel electrophoresis-based size verification, assembled dsDNA products were purified and cloned into a pBR-based plasmid containing an Amp-R marker gene using isothermal assembly and transformed into *E. coli* DH10B cells. Following plasmid extraction, each plasmid was validated using Sanger sequencing. Next, the sequence-verified 600 bp dsDNA fragments were assembled into 1000-2500 bp products, cloned into a pBR-based plasmid containing a kanamycin resistance gene a marker, using isothermal assembly as before, and sequence-validated using Sanger sequencing. Next, two or three 1000–2500 bp fragments were assembled into 4.5–8 kb fragments using the same strategy as before, sequence validated, and used for subsequent bacterial artificial chromosome assembly in yeast. Segment synthesis at GenScript USA Inc. followed the same strategy but with minor modifications. We synthesized and assembled 5 to 8 kbp intermediate dsDNA fragments, sharing 500 bp terminal homology with the neighboring fragment or, in the case of segment ends, with the pYES2L vector. These intermediate fragments were directly constructed from chemically synthesized ssDNA fragments and cloned into either pUC57 high-copy or pCC1 low-copy BAC-based vectors. The use of low-copy vectors allowed us to mitigate frequent instability issues due to the toxicity of endogenous *E. coli* genes in *E. coli* cloning hosts^98,99^. Following fragment assembly and sequence validation, yeast assemblies were performed in *Saccharomyces cerevisiae* W303, BY4733, or MaV203 using the intermediate dsDNA fragments, mixed at a 6:1 molar ratio of fragment to vector, according to a previously published procedure^100^, and validated using Illumina whole-BAC sequencing. The target BAC vector of yeast assemblies, pYES2L (Supplementary Data 1), was *de novo* synthesized based on the pYES1L-URA (Addgene plasmid #84301) plasmid and contains an additional oriV origin-of-replication for potential conditional amplification in EPI300 cells (Epicentre). Following yeast assembly, segments at BGI Research were isolated from 4 ml yeast culture by lysis and then electroporated into *E. coli* MDS42 Δ*recA* cells carrying the genomically integrated Segment 69 of Ec_Syn57 and, following overnight recovery at 32 °C, plated on LBL agar plates containing 80-100 μg/ml spectinomycin. Plates were incubated at 32 °C until colony formation. Following yeast assembly and sequence validation using Illumina whole-BAC sequencing, the pYES2L-Segment assemblies were electroporated into *E. coli* MDS42 Δ*recA* cells and, following overnight recovery at 32 °C, plated on 2×YT agar plates containing 80 μg/ml spectinomycin, and plates were incubated at 32 °C until colony formation. Following MASC PCR validation using the MASC PCR primer set described in Supplementary Data 1 and BAC extraction using the ZR BAC DNA Miniprep Kit (Zymo Research) based on the manufacturer’s protocol, pYES2L segment assemblies were validated using Illumina whole-BAC sequencing.

### Transcriptome and translatome analysis

We explored transcriptomic and translatomic changes by collecting cell from mid-exponential cultures using rapid filtration, and then performing RNA-seq and ribosome profiling from the same cell samples. As we sought to explore the native transcriptome and translatome of analyzed cells and the addition of antibiotics and cell-harvest conditions are known to influence ribosome profiling experiments^101,102^, we performed cell growth and sample collection without the presence of antibiotics and carried out all cell harvest steps immediately following cell growth. To minimize changes in cell state, we implemented the rapid filtration technique described in Ref^101^. Experiments were performed in three independent replicates, initiated from separate starter cultures. Cells from single colonies were inoculated into Lysogeny Broth Lennox (LBL) and grown at 37 °C, 250 rpm for 18 hours and then diluted into 500 ml 37 °C prewarmed LBL to an OD600 of 0.02 in a 2000 ml baffled Erlenmeyer flask with a vented cap. Cells were grown at 37 °C, 250 rpm until cultures reached OD600 = 0.40-0.45. Next, cells were immediately collected by pouring 400 ml culture into a Nalgene™ Rapid-Flow™ filter unit with 0.45 µm PES membrane (169-0045) and vacuum filtered. Cells were then washed off from the filter membrane using 1 ml prewarmed LBL, and immediately dripped into liquid N2. Once frozen, cell suspensions in liquid nitrogen were supplemented with 1 ml ice-cold lysis buffer (containing 10 mM MgCl2, 100 mM NH4Cl, 20 mM Tris pH 8.0, 0.1% NP-40, 0.4% Triton X-100, 100 U/ml of RNase-free DNase I (New England Biolabs, USA), 0.5 U/μl of SUPERase•In RNase Inhibitor (Invitrogen, catalog number AM2696) and 2 mM chloramphenicol). Frozen cell samples with lysis buffer were pulverized using a liquid nitrogen pre-chilled Cell Crusher Tissue Pulverizer (Cell Crusher, 607KSL) according to the manufacturer’s protocol. Samples were kept in liquid nitrogen throughout the entire procedure. Cell extracts were then centrifuged at 14,000 ×g for 20 minutes at 4 °C to remove cell debris. The supernatants were collected, and the RNA concentration in each sample was determined using the Qubit RNA high-sensitivity assay kit (Invitrogen USA). Ribosome profiling was performed according to the protocol described earlier^101^. In brief, the clarified cell lysates were digested with micrococcal nuclease from *Staphylococcus aureus* (MNase, Roche, USA) for 1 hour at 25 °C under continuous agitation. Digestions were stopped by supplementing samples with EGTA (Millipore-Sigma, USA) to a final concentration of 6 mM. Next, the monosome fraction was purified by size exclusion chromatography on Amersham MicroSpin S-400 HR columns (Cytiva, USA). Following monosome isolation, samples were denatured by adding SDS to a final concentration of 1%, and the RNA fraction was extracted and resolved on a 15% TBE-urea denaturing gel (Invitrogen, USA) according to the manufacturer’s protocol. The RNA band between 15 to 45 bp, corresponding to ribosome footprints, was excised, gel purified, and resuspended in 15 μl of 10 mM RNase-free Tris buffer, pH 7.0. The 3′ end of isolated RNA was subsequently dephosphorylated using T4 polynucleotide kinase (New England Biolabs) at 37 °C for 1 hour based on the manufacturer’s protocol. Next, the RNA was purified and resuspended in 10 μl of 10 mM RNase-free Tris buffer, pH 7.0. Ribo-seq RNA libraries were prepared for sequencing using the SMARTer^®^ smRNA-Seq Kit for Illumina (#635030) from Takara Bio USA, Inc., according to the manufacturer’s protocol. Finally, sequencing-ready libraries were sequenced on an Illumina NovaSeq 6000 to generate more than 30 million single-end 50 bp reads from each sample.

We obtained matched strand-specific (stranded) RNA-seq libraries for each Ribo-Seq library by extracting total RNA from the clarified cell lysates in parallel with Ribo-seq library preparations. Sequencing libraries were prepared according to our earlier protocol^2^, with minor modifications. Briefly, the rRNA-depleted and fragmented total RNA fraction was converted to sequencing libraries using the SMARTer^®^ smRNA-Seq Kit for Illumina (Catalog number #635030) from Takara Bio USA, Inc., according to the manufacturer’s protocol that generates strand-specific sequencing libraries. Library concentrations were measured using Qubit and analyzed for fragment length distribution using an Agilent 2100 Bioanalyzer. Finally, RNA-seq libraries were sequenced on an Illumina NovaSeq 6000 to generate more than 25 million single-end 50 bp reads from each sample. Demultiplexing, quality control, and adapter- and poly-A-trimming were performed with bcl-convert (version 3.9.3). cDNA reads were aligned to their corresponding reference by using Bowtie2 2.3.0^94^ in --very-sensitive-local mode. Sequencing reads mapping to rRNA genes were excluded from downstream analyses, and ribosome-footprint length distributions were determined based on ribosome footprint reads mapping to protein-coding CDSes. Next, we performed metagene analysis as described in reference^101^ to determine the offset position of the ribosomal P-site within Ribo-seq sequencing reads for each sample. Following metagene analysis, ribosome-footprint P-sites were mapped to their corresponding genomic reference sequence. Finally, translation levels of all protein-coding genes were determined as the number of ribosome footprints mapped by P-site location to each protein-coding CDS region. Transcription levels were evaluated by mapping RNA-seq reads using Bowtie2 2.3.0^94^ in --very-sensitive-local mode. Read counts and expression metrics were determined for each CDS, and differential expression analysis was performed using the EdgeR package (version 4.0.12, Ref^103^) using the standard settings. Relative ribosome A-(-aminoacyl-)-site occupancy of sense codons in *E. coli* MDS42 and Ec_Syn61Δ3 was determined based on the number of Ribo-seq reads mapping to a given codon position and the frequency of the given codon across all analyzed coding sequences. Analyses of codon occupancies were based on the set of protein-coding genes (CDS) in the given genome that (1) are not annotated as pseudogenes and (2) do not include internal in-frame stop codons. Codons overlapping between neighboring genes were also excluded. Finally, we also excluded protein-coding genes with low translation levels by excluding all genes displaying a normalized ribosome occupancy level below 50.0 (*i.e.*, RPKM ≤ 50.0). We evaluated codon occupancy at the A site for ribosome footprints in each sample, following the method described by Nedialkova and Leidel (2015)^104^. In each sample, we first determined the offset of the A-site within ribosome footprints using metagene analysis by choosing the P-site offset that aligned and formed a peak in coverage at the Translation Initiation Site (*i.e.*, the AUG start codon). A-site coordinates were derived from metagene analysis-determined P-site locations. A-site fractional coverage over codon frequency values in each sample and for all sense codons were determined as the log2 ratio of the codon frequency normalized codon coverage over codon frequencies averaged at the A-site. Finally, translation efficiency was evaluated as the ratio between Ribo-seq and RNA-seq coverage for the given protein-coding gene.

To ensure accurate genome and protein-coding gene annotations, we utilized *de novo* assembled, reannotated, and manually inspected reference genome sequences in all transcriptomic, translatomic, and proteomic analyses. Gene annotations were transferred based on sequence homology using the BLAST function implemented in Geneious Prime® 2023.2.1. from the genome of *E. coli* K-12 MG1655 (NCBI GenBank ID U00096.3) and manually inspected, as indicated in section “Bacterial genome sequencing and annotation” above.

Translation Initiation Rate (TIR) predictions for each protein-coding gene of *E. coli* MDS42 and Ec_Syn57 were performed using the Ribosome Binding Site Calculator Version 2.1.1 (available at www.denovodna.com)^84–86,90^. We predicted TIR values for each coding sequence by extracting the complete sequence of each CDS, including a 100 bp-long UTR, and TIR values were predicted at the annotated start codon of each CDS. The relative TIR for each CDS in its recoded form was determined by comparing the resulted TIR dataset from Ec_Syn57 to the TIR values obtained for the same CDS in the parental *E. coli* MDS42.

### Cappable-seq sample preparation and analysis

We identified transcriptional start sites across the genome of *E. coli* MDS42 and its 57-codon, partially recoded derivative containing Segments 1 to 8, Segments 9 to 18, 36 to 44, 46 to 49, and 51 to 59 using Cappable-seq, a transcriptomic method that enriches and sequences nascent, unprocessed triphosphorylated mRNAs originating directly from bacterial promoters. mRNA samples for Cappable-seq analysis were collected by inoculating both strains from single colonies into Lysogeny Broth Lennox (LBL) and grown at 37 °C, 250 rpm for 18 hours. Cultures were then diluted into 50 ml of 37 °C prewarmed LBL to an OD_600_ of 0.02 in a 300 ml baffled Erlenmeyer flask with a vented cap. Cells were grown aerobically at 37 °C, 250 rpm until cultures reached OD600 = 0.43-0.47, submerged in ice-water slurry for one minute with constant agitation, and cells were immediately collected by centrifugation at 5000 ×g at 4 °C for 8 minutes. Next, following the removal of the supernatant, cell pellets were immediately dropped into liquid nitrogen. Samples were stored at -80 °C until RNA extraction. Cell samples were prepared in duplicates. Total RNA from frozen samples was extracted by using the RNeasy Midi Kit (Qiagen, USA) according to the manufacturer’s instructions with in-column DNase treatment with the RNase-Free DNase Set (Qiagen, USA). Total RNA samples were then converted to Cappable-seq libraries according to the published protocol^59^ available at www.neb.com/en-us/protocols/2018/01/19/cappable-seq-for-prokaryotic-transcription-start-site-determination, with minor modifications. Modifications included the use of the Zymo Research RNA Clean & Concentrator-5 kit (R1015, Zymo Research, USA) for RNA purification and recovery instead of AMPure XP beads and the use of heat-denaturation to release streptavidin-bound 3’-Desthiobiotin-GTP (New England Biolabs, N0761) capped RNA. Following enrichment and decapping, RNA samples were converted to sequencing libraries using the NEBNext Small RNA Library Prep kit for Illumina (New England Biolabs, E7300) using the manufacturer’s protocol. Sequencing libraries were then sequences on a NextSeq 2000 sequencer using the Illumina P3 100 cycle flow cell and the P2 100 cycle flow cell (Illumina, USA) to generate approximately 0.4-1.5×10^9^ 50 bp paired-end reads. We applied ultradeep sequencing (*i.e.*, 1000-4000× average coverage across the entire genome, approximately tenfold higher than in prior Cappable-seq experiments^59^) to identify even weakly active transcriptional start sites (TSS) with high confidence. Non-mapping reads, formed due to adaptor-dimer formation in the NEBNext Small RNA Library Prep kit, were discarded from follow-up analyses. We identified TSS across each strain’s corresponding reference genome by mapping R1 reads using Bowtie2 2.3.0^94^ in local mode with -L 16, implemented in Geneious Prime® 2023.2.1. Alignments were exported as .bam files and analyzed using the previously described suite of programs for the identification and filtering of TSS data. Scripts and data analysis protocols are freely available on GitHub (https://github.com/elitaone/cappable-seq) and described in references^59,77^. Following the identification of genomic TSS and their direction, each TSS’s mRNA output was quantified as the TPM (transcript per million) of Cappable-seq reads originating from a given TSS. Finally, TSS coordinates were binned based on their location (intragenic or intergenic, and residing within the recoded chromosomal region or outside^59^) and each category was quantified, followed by averaging each category across the two independent replicates. We utilized the same data analysis workflow to measure antisense transcription using our stranded RNA-seq data.

We relied on previously generated Pacific Biosciences long-read SMRT-Cappable-seq data from *E. coli* K-12 MG1655 to analyze operon structures and identify their corresponding promoter on the genome of Ec_Syn57. Sequencing data (Sample GSM3290315, MG1655_rich_PacBio RS II, generated in rich bacterial medium and harvested in late exponential phase (*i.e.*, an OD600 = 0.55-0.6)) was downloaded from Gene Expression Omnibus dataset GSE117273^60^. Pacific Biosciences RS II circular consensus sequencing (CCS) reads were mapped to Ec_Syn57’s genome using Minimap4 (version 2.24) using -t 10 -x map-pb as settings, implemented in Geneious Prime® 2023.2.1. TSS coordinates were identified as the most 5’-mapped nucleotide of each sequencing read. Synonymous recoding impacted promoters were manually identified as sequences 0-80 bp upstream of each TSS that overlapped with intergenic mutations or codon changes on Ec_Syn57’s genome. Promoter troubleshooting DIvERGE oligonucleotides based on SMRT-Cappable-seq-identified promoter locations were designed and synthesized as described earlier^62^.

### Promoter transcriptomics

We synthesized 12 select promoters in their wild-type parental, *E. coli* MDS42-derived and 57-codon, Ec_Syn57-derived version, and individually cloned them into a pCC1-4k single-copy BAC vector, harboring the *E. coli* F plasmid origin-of-replication. The complete sequence of the 24 plasmids is available in Supplementary Data 1. We analyzed these promoter variants’ mRNA output in *E. coli* MDS42 cells using coupled DNA- and mRNA-sequencing in 5 independent biological replicates sequenced in two technical replicates (*i.e.*, 10 replicates in total for each promoter variant). To measure RNA expression in a multiplexed manner, we first barcoded each plasmid library member downstream of the assayed promoter using a 30-bp barcode containing randomized (*i.e.*, N) positions. Each barcode was sample-specific and contained degenerate positions to allow the measurement of DNA and mRNA levels of multiple variants for the same promoter-plasmid. Consequently, every promoter variant in our screen was assayed in multiple biological and technical replicates and also multiplexed across multiple members of the plasmid library in each experimental replicate. Following plasmid synthesis and purification, plasmids to be barcoded were diluted to 2 ng/µl and PCR amplified using the primers containing the full barcode sequence using KAPA HiFi HotStart ReadyMix (KK2602, Roche, USA). Barcoded PCR products were run on a 1.2% agarose gel, and the amplified plasmid DNA was extracted using the Monarch® DNA Gel Extraction Kit (New England Biolabs, USA) according to the manufacturer’s protocol. Barcoded plasmids were circularized by mixing 4 μl of the eluted, extracted PCR product with 2 μl of NEBuilder® HiFi DNA Assembly Master Mix (New England Biolabs, USA) according to the manufacturer’s protocol. Following assembly and DNA cleanup, the barcoded plasmids were individually transformed into electrocompetent TransforMax EPI300 (LGC Bioresearch Technologies, USA) cells and, following 1 hour of recovery, plated on LBL agar plates containing 12.5 μg/ml chloramphenicol. Next, we extracted plasmids from 16 independent colonies from each construct (16×24 in total), using the 96-Well Plate Plasmid DNA Mini-Preps Kit (Vacuum Based, B814152-0005) from Bio Basic, Inc. according to the manufacturer’s protocol. Plasmids were then transformed into electrocompetent *E. coli* MDS42 cells in 5 independent replicates to generate the assayed promoter-plasmid library. To generate sequencing libraries, overnight cultures of the promoter library, grown in LBL broth containing 12.5 μg/ml chloramphenicol at 37 °C aerobically, were diluted 1:200 into 100 ml LBL containing 12.5 μg/ml chloramphenicol, and incubated at 37 °C under continuous shaking. Upon reaching OD_600_ of ∼0.8, cultures were split into two technical replicates, quickly chilled on ice under constant agitation, and 50 ml from each culture was pelleted at 4 °C for subsequent DNA and RNA extraction.

Plasmid DNA was extracted from cultures using the EZ-10 Spin Column Plasmid DNA Miniprep Kit (Bio Basic, Inc.) according to the manufacturer’s protocol and the barcode region was amplified for Illumina amplicon sequencing using primers 5’-ATCGCAAACGTCACGGCTAATGGAAGCGTTCAACTAGCAG and 5’-GTGACATTTATGACGCGGGCCCGCGTCATAAATGTCACAC, followed by barcoding PCRs to multiplex samples. Total RNA from each sample was extracted using the PureLink RNA Mini Kit (Invitrogen, USA) and converted to first-strand cDNA using SensiFAST First-strand cDNA synthesis kit (BIO65054, Meridian Life Science) with 1 μg RNA as input. Amplicon RNA-seq libraries were constructed in the same way as plasmid DNA libraries outlined above, but 5 μl of the synthesized cDNA was used as the template for the amplification of the barcode region. Finally, purified amplicon DNA was sequenced with 20% PhiX control DNA spike-in on an Illumina MiSeq instrument using the MiSeq v2 chip to generate 250 bp paired-end reads. Relative expression levels were determined using the following steps. First, raw Illumina reads were processed to obtain per-barcode read coverage by merging paired-end reads using bbmerge from bbmap ver. 38.82. Next, reads were trimmed to remove all bases flanking the 30 bp barcode sequences using cutadapt ver. 3.4, and each barcode was counted across all samples. Next, barcode counts were first corrected for library sequencing depth by dividing the read count for the specific barcode by the number of reads obtained for the given DNA-seq or RNA-seq library. Next, the relative expression of each barcoded promoter was obtained by dividing the sequencing-depth-corrected RNA-seq barcode count by the sequencing-depth-corrected DNA-seq barcode count generated from the same culture and technical replicate. Finally, the mean relative expression for each specific promoter barcode was calculated across all replicates, and the promoter relative expression levels were calculated and plotted based on the resulting data for each promoter of interest. All utilized promoter-plasmid and barcode sequences are available in the Supplementary Data 1 of this work; Supplementary Data 5 contains the source data obtained from promoter transcriptomics experiments.

### SynOMICS-based genome assembly

We performed SynOMICS genome construction cycles in *E. coli* MDS42 Δ*recA* cells or in *E. coli* DH10B cells, containing the pRedCas2 plasmid. The pRedCas2 contains the cI857 repressor-pL promoter-driven Lambda-Red operon, consisting of *gam*, *exo*, and *bet* (from pORTMAGE-3^63^, Addgene plasmid # 72678; http://n2t.net/addgene:72678; RRID:Addgene_72678), a p15A origin-of-replication, a constitutively expressed chloramphenicol resistance marker and SpCas9 and tracrRNA (from pCas9^40,105^, Addgene #42876). The complete sequence of pRedCas2 is available in Supplementary Data 1. BACs containing individual recoded chromosomal regions were isolated using the ZR BAC DNA Miniprep Kit (Zymo Research) based on the manufacturer’s protocol and 500-1000 ng BAC DNA was electroporated into recipient cells using our standard electroporation protocol^2^. Following electroporation, cells were allowed to recover overnight and then plated onto 2×YT agar plates containing 80 μg/ml spectinomycin and 20 μg/ml chloramphenicol and incubated at 32 °C until colony formation (*i.e.*, 2-10 days). In rare cases when electroporation did not yield colonies (*i.e.*, in the case of Segment 11 and 32), we performed F-plasmid mediated conjugation utilizing the oriT sequence present on our pYES2L plasmid (for a detailed protocol, see Supplementary Methods). Next, we confirmed the presence of the recoded chromosomal region, first using MASC PCR (Supplementary Data 1) in KAPA2G Fast Multiplex MasterMix (Roche, USA) and then by validating its sequence using Illumina whole genome sequencing. To perform genomic deletions, cells were diluted 1:100 into 50 ml 2×YT broth containing 50 μg/ml spectinomycin and 20 μg/ml chloramphenicol and grown at 32 °C, 250 rpm until early exponential phase. We eliminated the parental copy of the extrachromosomal BAC-carried recoded segment by co-electroporating in two to three 40 μl replicates, 6-10 μg of the corresponding genome-targeting pCRISPR plasmid^40,105^-derived constitutive crRNA expression plasmid and 8-10 μg of the corresponding SynOMICS deletion cassette (for the complete list of crRNA plasmids and deletions cassettes, see Supplementary Data 1). Prior to electroporation, the expression of *gam*, *exo*, and *bet* was induced by a brief, 15-minute heat shock at 42 °C, 250 rpm. Following electroporation, cells were recovered overnight in 2×YT broth at 32 °C under constant agitation and then plated in 0.5-50 ml batches onto 2×YT agar plates containing 11 μg/ml gentamicin, 20 μg/ml chloramphenicol, and 50 μg/ml kanamycin. Plates were incubated at 32 °C until colony formation (*i.e.*, 2-12 days). Plated recovery cell culture volumes were determined and adjusted based on the recovery culture’s optical density prior to plating. We confirmed the deletion of the parental chromosomal copy using colony PCR with genome-specific, external primers, hybridizing 200-500 bp outside the flanking homology of the SynOMICS deletion cassette. All colony PCRs were performed using Platinum™ II Hot-Start Green PCR Master Mix (Thermo Fischer Scientific, USA). The sequence of all PCR primers is provided in Supplementary Data 1. Colonies displaying the correct PCR amplicon size (*i.e.*, ∼2700-3000 bp) were restreaked to 2×YT agar plates containing 8 μg/ml gentamicin, 20 μg/ml chloramphenicol, and 50 μg/ml spectinomycin and incubated at 32 °C until colony formation (*i.e.*, 2-12 days). Finally, the loss of the parental segment copy was confirmed using MASC PCR and Illumina whole genome sequencing. We integrated the extrachromosomal, BAC-carried segment by diluting cells from an overnight starter culture into 50 ml 2×YT broth, containing 1% D-glucose and 20 μg/ml chloramphenicol, and growing cells at 32 °C, 250 rpm until the early exponential phase. Following standard electrocompetent cell preparation^40,105^ and the induction of *gam*, *exo*, and *bet* off of pRedCas2 by a brief, 15-minute heat shock at 42 °C, 250 rpm, we electroporated 6 μg pINTsg plasmid. Cells were allowed to recover overnight at 32 °C in a rotor drum in 2×YT broth containing 1% D-glucose, washed once using 2×YT broth containing 1% D-glucose, and plated in 5 μl, 200 μl, and 500 μl volumes onto 2×YT agar plates containing 50 μg/ml carbenicillin, 20 μg/ml chloramphenicol, and 1% D-glucose, and incubated at 32 °C until colony formation (*i.e.*, 2-12 days). Cells from all SynOMICS deletion and integration experiments were plated on agar plates in 14.5 cm diameter Greiner Bio-One Petri dishes (VWR Catalog No. 82050-600). Following colony formation, segment integrations were confirmed using colony PCRs checking the terminal junctions between the segment and the neighboring genomic end. Colonies displaying the correct PCR amplicons were then restreaked to 2×YT agar plates containing 50 μg/ml carbenicillin, 20 μg/ml chloramphenicol, and 1% D-glucose, and the integrity of the recoded chromosomal region was validated using MASC PCR and whole genome sequencing. Finally, cells were made suitable for the next SynOMICS integration cycle by growing them in 2×YT broth containing 20 μg/ml chloramphenicol and 0.6% L-rhamnose (CAS number: 10030-85-0) at 32 °C, inducing the expression of the pRham (*rhaB*) promoter-driven *hok* bacterial toxin that resulted in the rapid loss of the pINTsg plasmid. Given the rapid loss of the pINTsg plasmid in the presence of L-rhamnose, we could directly proceed with the next SynOMICS cycle by electroporating the next segment. crRNA and sgRNA plasmids used in SynOMICS experiments were synthesized by GenScript USA as research-grade, primarily supercoiled plasmids, delivered freeze-dried, and resuspended to 1 μg/μl in nuclease-free H2O. On the course of SynOMICS cycles, we initiated DIvERGE-, MAGE-, and/or ALE-based troubleshooting after the deletion or integration steps, depending on cell fitness. The details of troubleshooting for each segment are summarized in Supplementary Methods. All primer sequences used during genome assembly are listed in Supplementary Data 1.

### Genome editing-based troubleshooting

Slow-growing variants from SynOMICS cycles were transformed with pORTMAGE plasmids^63^, selected based on the antibiotic resistance phenotype of the given strain. Strains originating from SynOMICS deletion steps (*i.e.*, Gentamicin-R and Kanamycin-R) were made electrocompetent using our standard protocol^2^ and electroporated with pORTMAGE-2 (Addgene plasmid # 72677; http://n2t.net/addgene:72677; RRID:Addgene_72677) or alternatively, pORTMAGE203D, a derivative of pORTMAGE-2 expressing CspRecT^106^ instead of *gam*, *exo*, and *bet.* Strains originating from the integration step of SynOMICS were transformed with pORTMAGE-Ec1^106^, unless noted otherwise in the Supplementary Methods. pORTMAGE plasmid containing cells were induced and made electrocompetent as described earlier^62,107^, and 40 μl freshly made electrocompetent cells were electroporated with 1 to 2 μl of 500 μM mixture of MAGE and DIvERGE ssDNA oligonucleotides in two replicates, targeting potential promoter-recoding- and DNA-synthesis-derived errors in the given strain. For a detailed list of all oligonucleotides, see Supplementary Data 1. All oligonucleotides chemically synthesized either by GenScript, USA or by Integrated DNA Technologies, USA and resuspended in IDTE buffer, pH 8.0 (Integrated DNA Technologies, USA) to 500 μM final concentration. Following oligo-delivery, cells were allowed to recover overnight in SOB or 2×YT broth, and electroporation cycles were repeated as described in Supplementary Methods for each segment. Following electroporation cycles, cells were diluted twice 1:100 in 50 ml SOB or 2×YT broth in a 300 ml Erlenmeyer flask with vented cap, grown aerobically between each dilution to saturation aerobically, and then plated onto 2×YT agar plates and incubated until colonies formed. Finally, individual fast-growing colonies from each experiment were isolated—based on colony size following colony formation on 2×YT agar plates—and subjected to whole-genome sequencing. Tn1000 and Insertion Sequence (IS) deletions were performed similarly, but the corresponding MAGE oligonucleotides, targeting the complete deletion of the given mobile genetic elements, were co-electroporated with a variant of the pCRISPR plasmid^40,105^, carrying a crRNA guide sequence to cleave the genomic or BAC-carried Tn1000 or IS element (Supplementary Data 1). For a detailed description of the utilized crRNA sequences, mobile genetic element removal experiments, see Supplementary Methods. The deletion of the genomic *recA* was performed as described before^2^.

### Adaptive laboratory evolution-based troubleshooting

We performed adaptive laboratory evolution experiments in rich bacterial medium on the course of SynOMICS genome construction cycles to evolve variants with increased fitness. At each daily transfer step, ∼5×10^9^ cells were transferred into 500 ml Lysogeny Broth Lennox (LBL) supplemented with Tris/Tris-HCL as buffer and incubated aerobically for 24 hours at 37 °C, 250 rpm in a 2000 ml baffled Erlenmeyer flask with vented cap. Adaptive laboratory evolution experiments were performed in two replicates, starting from the same seed culture or two isogenic clones of the starting strain. During ALE experiments, cells were plated onto 2×YT agar plates following every five transfers and incubated at 37 °C to assess growth rate based on the speed of colony formation. Finally, evolution experiments were terminated by spreading bacterial cells onto 2×YT agar plates, incubated at 37 °C, and an individual colony from each experiment was isolated and subjected to whole-genome sequencing. The duration of each ALE experiment is listed in the Supplementary Methods. The identified mutations in the final evolved variants are available in Supplementary Data 7.

### Doubling time and growth measurements

To determine growth parameters under standard laboratory conditions, saturated overnight cultures, initiated from isogenic, single colonies, were diluted 1:100 into 100 µl 2×YT in a 96-well flat-bottom transparent microtiter plate (Corning Inc., USA). To assess growth kinetics, diluted cultures in ten replicates were incubated aerobically at 37 °C, 800 rpm, and 1-mm orbital shake in a BioTek Synergy H1 Multimode plate reader (Agilent Technologies, Inc., USA). Optical density at 600 nm (OD600) measurements were taken every 9 minutes until stationary phase was reached. We calculated the doubling time using the open-source GrowthRates package, version 4.4, according to the method described in Ref ^108^. Source data from growth assays are available in Supplementary Data 2.

### Total proteome analysis and the detection of translated genes

We analyzed the proteome of *E. coli* MDS42 as wild-type and Ec_Syn61 (Addgene #174513) as a 61-codon variant of *E. coli* MDS42 by performing tandem mass spectrometry-based proteome analysis. 10 ml mid-exponential phase cultures (OD600 = 0.45) of each strain, growing at 37 °C, 250 rpm in 2×YT broth in 300 ml baffled flasks, were spun down at room temperature and washed four times in 1 ml phosphate-buffered saline (Invitrogen, USA). Cell samples were immediately frozen in liquid N2 and stored at -80 °C until total protein extraction and proteome analysis.

Next, frozen samples were thawed on ice and digested using the S-Trap™ micro columns (ProtiFi, NY, USA) digest procedure according to the manufacturer protocol. Samples were then solubilized in aqueous 0.1% formic acid for subsequent analysis by tandem mass spectrometry. LC-MS/MS analysis of digested samples was performed on a Q Exactive Orbitrap Mass Spectrometer equipped with a UNO nano-HPLC (both from ThermoScientific, USA). Peptides were separated on a 150 µm inner diameter microcapillary trapping column packed first with 1.9 cm of C18 PepSep column. Separation was achieved by applying a gradient from 5% to 27% acetonitrile in 0.1% formic acid over 45 minutes at 200 nl/minute. Electrospray ionization was performed by applying a voltage of 2 kV using a custom electrode junction at the end of the microcapillary column and sprayed from metal tips (PepSep, Denmark). The mass spectrometry survey scan was performed in the Orbitrap in the range of 400-550, 550-750 and 750-1250 m/z as gas-phase-fractionation-based analysis at a resolution of 6×10^4^, followed by the selection of the ten most intense ions for fragmentation using Collision Induced Dissociation (CID) in the second MS step (CID-MS2 fragmentation) in the ion trap using a precursor isolation width window of 10 m/z, AGC (automatic gain control) setting of 10,000 and a maximum ion accumulation of 150 ms. Singly charged ion species were excluded from CID fragmentation. The normalized collision energy was set to 30 V, and an activation time of 10 ms was utilized. Ions in a 10 m/z window around ions selected for MS/MS as wide window acquisition mode were excluded from further selection for fragmentation for 60 seconds.

Proteome reference files were generated by identifying all potential Open Reading Frames (ORFs) longer than 20 amino acids using the canonical genetic code and TTG, ATC, ATG, CTG, ATT, GTG, and ATA^58^ as potential start codons on each genome sequence. The raw data was analyzed using Proteome Discoverer 3.1 (Thermo Fisher Scientific, USA) and Chimerys 2.0 (MSAID, Germany). Assignment of MS/MS spectra was performed using the Sequest HT algorithm by searching the data against a protein sequence database, including all ORF entries from the sample’s corresponding reference genome and other known contaminants such as human keratins and common lab contaminants. Peptide amounts in each sample were quantified by LFQ (label-free quantitation). Sequest HT searches were performed using a 10-ppm precursor ion tolerance and requiring each peptide’s N-/C termini to adhere with trypsin protease specificity while allowing up to two missed cleavages. Methionine oxidation (+15.99492 Da) was set as a variable modification. All cysteines were set to permanent no modification due to no alkylation procedure. An overall false discovery rate of 1% on both protein and peptide levels was achieved by performing a target-decoy database search using Percolator^109^. Next, peptides originating from previously not annotated cryptic ORF sequences were identified by screening all identified peptides from a given sample using ProteoMapper version 5.2.0 (ref ^93^, available from https://tppms.systemsbiology.net/pm/) to filter out all peptides matching known proteins on a given reference genome. Finally, all identified peptides were mapped back to genomic ORFs using the BLAST function implemented in Geneious Prime® 2023.2.1. and inspected manually. All identified cryptic ORFs and ORF-derived peptides are listed in Supplementary Data 8.

