## Supplementary Material - Synthetic genomes unveil the effects of synonymous recoding for "Synthetic genomes unveil the effects of synonymous recoding"

[†]Anush Chiappino-Pepe and Bogdan Budnik are shared co-second authors.

**Guide to Supplementary Information**

- Supplementary Methods. The detailed description of the genome construction and troubleshooting steps of Ec_Syn57: Steps of SynOMICS-based genome construction and troubleshooting; Genome assembly.
- Supplementary Figure 1. Frequent mobile genetic element transposition into synthetic genomic segments during genome synthesis.
- Supplementary Table 1. Comparison of existing prokaryotic genome construction methods.
- Supplementary Table 2. SynOMICS achieves challenging recoding schemes.
- Supplementary Figure 2. Optimization of the SynOMICS integration plasmid toward increased stability and efficiency.
- Supplementary Figure 3. Efficient synthetic segment integration using SynOMICS.
- Supplementary Figure 4. Ribosomes stall at forbidden codons present in Ec_Syn61Δ3 due to genomic annotation errors prior to genome design.
- Supplementary Figure 5. Ribosome A-(-aminoacyl-)-site coverage of sense codons in *E. coli* MDS42.
- Supplementary Figure 6. Proteomics- and ribosome-profiling-based detection of cryptic translated ORFs in *E. coli* MDS42.
- Supplementary Figure 7. Proteomics-based detection of a cryptic translated ORF in *glpB* of *E. coli* MDS42 and Ec_Syn61.
- Supplementary Figure 8. Existing recoded organisms harbor and tolerate forbidden codons.
- Supplementary Figure 9. Predicted effect of synonymous recoding on translation initiation rates of protein-coding genes in Ec_Syn57.
- Supplementary Figure 10. Multiplexed genome editing-based troubleshooting of Segment 21.
- Supplementary Figure 11. Troubleshooting synonymous recoding induced issues of *msbA* in Segment 18 and *pgk* in Segment 56 of Ec_Syn57.
- Supplementary Table 3. Fitness of parental strains and their recoded derivatives.
- Supplementary Figure 12. Ribosome profiling of genomes with altered genetic codes indicates large changes in translation.

- Supplementary Figure 13. Lack of correlation between predicted and measured changes in translation within the recoded Ec_Syn57 genome.
- Supplementary Figure 14. Direction and expression level of primary mRNA transcripts in *E. coli* MDS42.
- Supplementary Figure 15. Synonymous recoding induces widespread antisense transcription.
- Supplementary Table 4. Cost estimate of genome synthesis and troubleshooting.
- Source data. Uncropped gel electrophoresis pictures.
- Supplementary references.

**Supplementary Tables** (.xlsx files)

- Supplementary Data 1. DNA constructs; Plasmids utilized in promoter-transcriptomics experiments; MAGE & DIvERGE oligonucleotides; MASC PCR primers; SynOMICS deletion cassettes; SynOMICS genome-targeting CRISPR/Cas9 guide RNAs; Sequence of additionally recoded genes; Strains and plasmids used in this study.
- Supplementary Data 2. Source data for fitness measurements.
- Supplementary Data 3. Source data for RNA-seq and Ribo-seq experiments.
- Supplementary Data 4. Source data for TIR predictions.
- Supplementary Data 5. Source data for promoter transcriptomics experiments.
- Supplementary Data 6. DNA synthesis errors in synthetic genomic segments of Ec_Syn57.
- Supplementary Data 7. Identified mutations in *E. coli* strains containing the genome of Ec_Syn57.
- Supplementary Data 8. MS/MS identified cryptic ORFs in *E. coli* MDS42 and Ec_Syn61.

**Supplementary sequences** (.gb files)

- Annotated genome of *Escherichia coli* Ec_Syn57.
- Annotated sequence of the RK2-FCas7 conjugation helper plasmid.

### Supplementary Methods

##### Steps of SynOMICS-based genome construction and troubleshooting

Using the following steps, we assembled and troubleshot the synthetic genome of Ec_Syn57 in 11 *E. coli* strains.

**Segment 0**: We integrated Segment 0 in MDS42 Δ*recA* containing Segments 82, 83, 84, and 85. Our attempts to obtain SynOMICS parental-copy-deleted variants failed, indicating the nonviability of the computationally-designed Segment 0, and therefore, we troubleshot Segment 0 in a 3-step process. Based on the available list of *E. coli* promoters[1–3] and Cappable-seq results, we hypothesized that the reason for nonviability is the recoding of the promoter region in front of the growth-essential *ribF-ispH* operon and a 31 bp deletion in front of the growth-essential *folA*, impacting the gene's expression. We deleted the parental copy of this segment in MDS42 following our SynOMICS deletion workflow using a gentamicin resistance cassette with only external genomic homologies. Deletion yielded slow-growing colonies that took more than six days to form ~0.5 mm-sized colonies. Whole-genome sequencing confirmed the loss of the parental copy in these cells. Next, we initiated simultaneous DIvERGE- and MAGE-based troubleshooting to repair the *ribF-ispH* operon's promoter and revert the 31 bp deletion in front of *folA*. pORTMAGE2-based editing (ref [4,5] (Addgene plasmid # 72677 ; http://n2t.net/addgene:72677 ; RRID:Addgene_72677)) with MAGE oligo 5'-G*A*TTTCCCGATAAAAAAAATTGTCGCCACTATACGTAAAGCGTAAACCGTCGTCGACTGGTGCGAGGATGATGTTGAGGAAAATTTTA*T*A and DIvERGE-oligos 5'-TGTATGCCGCGTATCAGCTTCATGTCTGGCTCAAAACAGTGAAAATCGTCCGAGTATACCTTGTACAGCGGTTAGGGTTAGCCGGCGATT and 5'-TAGGGTTAGCCGGCGATTGAGTACCGAAATCAGAAAGAAGTGAGATTTCATTGCCATGGCGCAAATCACGGGAAGAAACTGACCGCCTGC, synthesized using 2% phosphoramidite-spiking, followed by growth-based selection identified a variant with improved fitness. This mutant reverted the CTA (Leu) recoded codon to wild-type parental TTA (Leu) in the -35 region of *ribF-ispH* and corrected the promoter region of *folA*. Star (*) in oligonucleotides indicates a phosphorothioate bond. The resulting Segment 0 variant was then transformed into MDS42 Δ*recA* containing Segments 82, 83, 84, and 85, and the deletion of the parental copy succeeded with our standard SynOMICS workflow. However, further troubleshooting was necessary due to the slow growth of the deleted variants. Therefore, we initiated ALE-based troubleshooting after the SynOMICS deletion step. Ten days of ALE-based troubleshooting yielded a clone containing a synonymous mutation that induced a TTA →TTG (Leu) codon change within the -35 region of the *ribF* promoter, allowing us to integrate Segment 0. Following integration, we performed an additional 41 days of ALE to increase the generated strain's fitness.

**Segment 1**: We integrated Segment 1 in MDS42 Δ*recA* containing Segments 2 and 3 using our standard SynOMICS method.

**Segment 2**: We integrated Segment 2 in MDS42 Δ*recA* containing Segment 3 using our standard SynOMICS method.

**Segment 3**: We integrated Segment 3 in MDS42 Δ*recA* using our standard SynOMICS method.

**Segment 4**: We integrated Segment 4 in MDS42 Δ*recA* containing Segments 1, 2, 3, 7, and 8 using our standard SynOMICS method. Our initial attempts to obtain SynOMICS parental-copy-deleted variants failed, and therefore, we troubleshot Segment 4 in a 2-step process. Based on the available list of *E. coli* promoters[1–3] and Cappable-seq results, we hypothesized that the reason for nonviability is the recoding of the promoter region in front of the growth-essential *ispU* and *accA*, impacting these and the downstream genes' expression. We restored the expression of these genes by integrating a BBa_J23105 constitutive promoter in front of *ispU* and *accA* while preserving the ribosomal binding site (RBS) of *ispU* and integrating an additional RBS (5'-CCCCGGGAGGTAAAA) in front of *accA*. The integration of these promoters was performed using CRISPR/Cas9 assisted MAGE in MDS42 Δ*recA* using a constitutive Cas9 and tracrRNA and thermally inducible CspRecT expressing plasmid (termed pF20Cas, conferring chloramphenicol resistance) and using MAGE oligos 5'-A*C*TACTTACGATCAGATGGCGCAGACTATATCACTGAAGCCCGCTAGCATAGTACCTAGGACTGAGCTAGCCGTAAATACGCTAACAAATAGCGCGACTCTCTGTAGCCGGATTATCCTC and 5'-C*T*GCAATCGGCTGTTCAAAATCAAGGAAATTCAGTGACATTTTTACCTCCCGGGGGCTAGCATAGTACCTAGGACTGAGCTAGCCGTAAAGTATTAGTCAAACTCCAGTTCTACTTGTTCCGAACCAATG. Edited clones were selected following MAGE cycles using a multiplexed non-repetitive sgRNA plasmid (based on pINTsg (Supplementary Data 1)) containing the 5'-AACTGGAGTTTGACTAATAC and 5'-TCGCGCTATTTGTTAGCGTA guide sequences. Edited clones were identified using allele-specific PCR and validated using whole-genome sequencing. The promoter-corrected variant of Segment 4 was then transformed into MDS42 Δ*recA* containing Segments 1, 2, 3, 7, and 8, and the parental copy of Segment 4 was deleted using the standard SynOMICS procedure. Whole genome sequencing indicated that the parental copy of Segment 4 could not be fully eliminated and the terminal genes, *dkgB* and *yafC*, retained their parental copy. We attempted integrating the resulting variant using SynOMICS; however, our attempts to identify integrants failed. Therefore, we initiated ALE-based troubleshooting on the parental-copy-deleted variant and evolved it for 30 days. The evolved variant was successfully integrated using SynOMICS, but *dkgB* and *yafC* retained forbidden codons, necessitating further troubleshooting. These codons were corrected during the integration of Segment 5.

**Segment 5**: We integrated Segment 5 in MDS42 Δ*recA* containing Segments 1, 2, 3, 4, 6, 7, and 8 using our standard SynOMICS method, however integration was performed twice due to the duplication of the target chromosomal region in the parental strain. As the terminal genes of Segment 4 retained forbidden codons and due to multiple potential design errors in Segment 5, we resynthesized this segment using our updated assembly workflow. In the new variant, termed Segment 5_2, we appended the recoded version of *dkgB* and *yafC* from Segment 4 to the left end of the segment. Furthermore, as our Cappable-seq results identified the promoters of *rnhA* and *dnaQ* as targets for troubleshooting, we corrected the expression of these two genes by inserting an SLP2018-2-3178 constitutive promoter in front of *dnaQ* and an SLP2018-2-2267[6] constitutive promoter in front of *rnhA*. These nonrepetitive promoters minimized the chances of unwanted intramolecular recombination[6]. Segment 5_2 was integrated using SynOMICS and the resulting strain was subjected to a new SynOMICS cycle using the same segment and CRISPR/Cas9 construct to delete the parental copy. Our lack of success to collapse the genomic duplication with CRISPR/Cas9 cuts in this strain indicated that the duplicated region (a 234,789 bp duplication from *yadS* to *dnaX*) is essential for the strain's survival.

**Segment 6**: We integrated Segment 6 in MDS42 Δ*recA* containing Segments 1, 2, 3, 4, 7, and 8 using our standard SynOMICS method. As the resulted cells displayed slow growth, we evolved MDS42 Δ*recA* containing Segments 1, 2, 3, 4, 6, 7, and 8 using ALE for 25 days.

**Segment 7**: We integrated Segment 7 in MDS42 Δ*recA* containing Segments 1, 2, 3, and 8 using our standard SynOMICS method. Our initial attempts to obtain SynOMICS parental-copy-deleted variants failed and therefore we troubleshot Segment 7 using CRISPR/Cas9-based editing. Based on the available list of *E. coli* promoters[1–3] and Cappable-seq results, we hypothesized that the reason for nonviability is the recoding of the promoter region in front of the growth-essential *ribE*, impacting the downstream *ribE, nusB, thiL*, and *pgpA*'s expression. We restored the expression of these genes by integrating a BBa_J23105 constitutive promoter in front of *ribE* while preserving the ribosomal binding site using CRISPR/Cas9 assisted MAGE in MDS42 Δ*recA* using pF20Cas and MAGE oligo 5'-G*A*TTTTAGCATAATATTTCGTGCGCTGCTTCCCTTTCGAGCCGCTAGCATAGTACCTAGGACTGAGCTAGCCGTAAAGGGAGATCATGCACCCACAAGATGCAGGCAAACATCCGGGCCT. Edited clones were selected following MAGE cycles using an sgRNA plasmid containing the 5'-GCATGATCTCCCGGCTCGAA guide sequence. Edited clones were identified using allele-specific PCR and validated using whole-genome sequencing. The *ribE* promoter-corrected variant could then replace the parental copy of this segment using the standard SynOMICS workflow.

**Segment 8**: We integrated Segment 8 in MDS42 Δ*recA* containing Segments 1, 2, and 3 using our standard SynOMICS method.

**Segment 9**: We integrated and corrected the DNA synthesis errors of Segment 9 in MDS42 Δ*recA* using SynOMICS. Segment 9 harbors a 4402 bp deletion on its right segment end, deleting the nonessential *fdrA-ybcF*. Therefore, we utilized the SynOMICS deletion cassette to repair our synthetic variant. We designed a variant of the standard SynOMICS deletion cassette that repairs *fdrA* while replacing the parental genomic copy (09GentRC-2-08w10w, Supplementary Data 1) Following the repair of *fdrA*, the segment was integrated without issues. We observed the transposition of a Tn1000 transposon into *cnoX* of this segment therefore we eliminated the insertion and repaired the gene using CRISPR/Cas9-asssited MAGE[7] using 5'-G*T*CTACAGCTAACCCCAATTCTGGAATCACTCGCGGCGCAGTACAACGGGCAATTTATTCTGGCGAAGCTGGACTGCGACGCGGAGCAGA as MAGE oligo and a crRNA plasmid, based on pCRISPR[8], containing a crRNA 5'- TGTGGCTCGTGCGATTTGTTACGGACAACG.

**Segment 10A**: Segment 10A was integrated using the standard SynOMICS workflow in MDS42 Δ*recA* containing Segment 9, 11, 13, 14, 15, 16, 17 and 18.

**Segment 10B**: Segment 10B was integrated using the standard SynOMICS workflow in MDS42 Δ*recA* containing Segment 9, 10A, 11, 13, 14, 15, 16, 17 and 18.

**Segment 11**: We integrated Segment 11 in MDS42 Δ*recA* containing Segments 9 using our standard SynOMICS method. We observed the transposition of a Tn1000 transposon into *hcxA* of this segment therefore we eliminated the insertion and repaired the gene using CRISPR/Cas9-assisted MAGE using 5'-

C*A*TGATTTTTCACTGATGAACAACTTTCTCGCGCGGTGTGGATCTACGGCAAACGCGCC ATTGCTGCGGCGCAAACCAAACTTCCGCCA as MAGE oligo and a crRNA plasmid, termed pCRISPRM-Tn4, containing a crRNA 5’-TGTGGCTCGTGCGATTTGTTACGGACAACG.

**Segment 12**: Segment 12 was integrated using the standard SynOMICS workflow in MDS42 Δ*recA* containing Segment 9, 10A, 10B, 11, 13, 14, 15, 16, 17, and 18. Our attempts to delete the parental copy of Segment 12 failed and we traced back this segment’s nonviability to a frameshift mutation in the growth-essential *leuS* and a 230 bp deletion in *pgm*. We corrected the frameshift mutation in *leuS* using MAGE and utilized SynOMICS to repair *pgm* by appending the deleted region to the deletion cassette (12GentRC-2-11r13r, Supplementary Data 1). Whole genome sequencing following the deletion of the parental copy revealed the reversion of the growth essential *seqA* to its parental form. Therefore, following the integration of this segment, we performed CRISPR/Cas9-assisted dsDNA recombineering to repair the forbidden codons in this segment. The cotransformation of a 1573 bp *ybiF-seqA* dsDNA PCR amplicon cassette with a 5’- TGCAAGCCAGGCTTGACGCTATCCGCTGCC crRNA expressing pCRISPR plasmid followed by Sanger target sequencing and Illumina whole genome sequencing of the *ybiF-seqA* locus identified fully recoded variants.

**Segment 13**: We integrated Segment 13 in MDS42 Δ*recA* containing Segment 9, 11, 14, 15, 16, 17 and 18 using SynOMICS.

**Segment 14**: We integrated and corrected the DNA synthesis errors of Segment 14 in MDS42 Δ*recA* containing Segment 9 and 11 using SynOMICS. Segment 14 harbors a 1259 bp deletion on its right segment end, deleting the nonessential *uvrB*, and therefore we utilized the SynOMICS deletion cassette to repair our synthetic variant. We designed a variant of the standard SynOMICS deletion cassette that repairs *uvrB* while replacing the parental genomic copy (14GentRC-2-13w15r, Supplementary Data 1). Following the repair of *uvrB*, the segment was integrated without issues. We observed the transposition of a Tn1000 transposon into *nadA* of this segment therefore we eliminated the insertion and repaired the gene using CRISPR/Cas9-assisted MAGE using 5’- C*A*GCATACGATTGAGCGGCACCAGCGCACGCTCTCGCAGTCGTTCATCAACATGAACCT CGTGATTAGATCCTTCCTGTTCAAGTGCCTC as MAGE oligo and pCRISPRM-Tn4, containing a crRNA 5’-TGTGGCTCGTGCGATTTGTTACGGACAACG.

**Segment 15**: We integrated and corrected the DNA synthesis errors of Segment 15 in MDS42 Δ*recA* containing Segment 9, 11, and 14 using SynOMICS. The integration of this segment was performed using the standard SynOMICS workflow, but due to DNA synthesis errors that affected genes *ybiB-glnH*, including the transposition of a Tn1000 into *hcxB*, we deleted this non-growth-essential region using CRISPR/Cas9-assisted oligo-recombineering using 5'-GATGGTGAAAGCTAACAATAACGATGTGAAATCAGTGAAAGATTACGTTGGCGTAATACCCATAAGTTCAAAAATGGTTTCAGTCAGCAC and pCRISPRM-Tn4.

**Segment 16**: We integrated Segment 16 in MDS42 Δ*recA* containing Segment 9, 11, 14, and 15 using SynOMICS. We first eliminated a copy of Tn1000 from this segment from *ybjG* in MDS42 Δ*recA* using CRISPR/Cas9-assisted oligo-recombineering using 5'-C*A*GTATACAAAGGTCCCTTTTCAGGGACCCTTTCGTATGTTGTAGGAACTGAGAGCTCTAAATTTAAATATAAACAACGAATTATCTCCT and 5'-C*C*ATATAGAGCGTGTTCCACAGAACCACTGCATTAGTAACCAGGCGTAACGCAGTTCAGAGACGAGATATTATAGGCACCAATGGTGGCA and pCRISPRM-Tn4. These MAGE oligos deleted the internal ORFs of Tn1000, however left a scar, comprising the transposon ends, within Segment 16.

**Segment 17**: Segment 17 was integrated using the standard SynOMICS workflow in MDS42 Δ*recA* containing Segment 9, 11, 14, 15, and 16. Our attempts to delete the parental copy of Segment 17 failed and based on sequencing data, we traced back the nonviability of this segments to a 13 bp deletion in front of the growth-essential *infA*, a 10 bp deletion in the essential *cydD*, and a 11 bp deletion in *ftsK*. We repaired the deletions affecting *infA* and *cydD* using MAGE. After three MAGE cycles with oligos 5'-T*T*TAGCGCGCAAATCTTTACTTATTTACAGAACTTCGGCATTATCTTGCCGGTTCAAATTACGGTAGTGATACCCCAGAGGATTAGATGG and 5'-G*C*TTCCCTTTACGCTTCTGGTTCTGACCTTTGTACTGCGCGCATGGGTGGTCTGGCTACGCGAACGGGTGGGTTATCACGCCGGGCAGCA, edited clones were identified using allele specific PCR. Despite being described as a growth essential gene[9], in our experiments, the 11 bp deletion in *ftsK* did not prevent the deletion of Segment 17's parental copy. Following the transformation of the corrected Segment 17 into MDS42 Δ*recA* containing Segment 9, 11, 14, 15, and 16, we deleted the parental copy of Segment 17 and integrated this segment using the standard workflow.

**Segment 18**: Segment 18 was integrated using SynOMICS in MDS42 Δ*recA* containing Segment 9, 11, 14, 15, 16, and 17. Our initial attempts to obtain SynOMICS parental-copy-deleted variants failed and therefore we troubleshot Segment 18 using DIvERGE- and MAGE-based editing. Based on Cappable-seq and promoter-transcriptomics results, we hypothesized that the reason for nonviability is recoding changes in the promoter region of the growth-essential *msbA-lpxK-ycaQ* operon. The promoter for this operon lays upstream in *ycaI* and a recoded codon overlaps with the -10 region of this promoter. We performed SynOMICS-based troubleshooting on this strain by deleting the parental copy in MDS42 cells. Clones containing the recoded segment only displayed slow growth. Therefore, next, we performed DIvERGE-based randomization using pORTMAGE2 (Addgene plasmid # 72677) and 5'-CGGAATGTCTGCCACGTAGAGAGATCTTTGTCGTTATGCATTCAAAAACCAGCATTTGTTGAAATAGCCGCATATTTCACCCGTTATCC synthesized using 2% phosphoramidite-spiking[5]. Growth-based selection identified a variant that mutated a GGC glycine codon to a missense GAC wild-type TTA in the -35 region of the *msbA* promoter. The resulted Segment 18 variant was then transformed into MDS42 Δ*recA* containing Segments 9, 11, 14, 15, 16, and 17 and the deletion of the parental copy succeeded with our standard SynOMICS workflow. Finally, we integrated Segment 18 using pINTsg.

**Segment 19**: We integrated Segment 19 in MDS42 Δ*recA* containing Segment 20, 21, 22, 23, 24, 25, 26, 27, 28, and 29 using our standard SynOMICS method.

**Segment 20**: We integrated Segment 20 in MDS42 Δ*recA* containing Segment 21, 23 and 24 using our standard SynOMICS method. The resulted colonies displayed low fitness and therefore we performed ALE-based troubleshooting on two variants for 30 days.

**Segment 21**: We integrated Segment 21 in MDS42 Δ*recA* our standard SynOMICS method. As the replacement of the parental copy of this segment resulted is severe fitness drop (*i.e.*, a doubling time of 253 minutes in rich bacterial broth at 37 °C; **Supplementary Table 3**). Based on Cappable-seq we identified the intragenic promoters of the growth-essential *fabH-acpP* and the *tmk-holB* operons as the cause of fitness decrease (**Supplementary Figure 10**). We restored the expression of these operons by performing MAGE to insert a library of constitutive promoters (5'- TTKACGGCTAGCTCAGTCCTAGGTACWRTGCTAGC) in front of each target gene. Following three MAGE cycles using an equimolar mixture of 5'-T*A*TATACCGTCACTTGCAAACTGCGAGTTCGCTGGCAGCGTCCTGGCTAGCAYWGTACCTAGGACTGAGCTAGCCGTMAATCACCGCAGAGTTCCTGATTTGCCACCGTCCAGCAGCTCAAAA*C*C, 5'-C*A*CGACGCTTAGACACGTTTGTCCTCCAGGGAGGGAAAAAATGATTGCTAGCAYWGTACCTAGGACTGAGCTAGCCGTMAACTAGTGGGACAAAAAGATAAAACTCAGGCGGTCGAAC*G*A, and 5'-C*A*ATGACGATATACTTTGAGCGCATTTTTTTCCTTAAGCACTTTCGCTAGCAYWGTACCTA

GGACTGAGCTAGCCGTMAATTACTGGGCATTCTTCTCTTTTAGTACCTTGAGATAATCCTG C*A*C and growth-based selection, fast growing variants were identified containing three promoter insertions. The identified promoter variants in the fastest-growing strain were then used in a new genome editing experiment to validate the effect of these promoter insertions and rule out the potential effect of genomic compensatory mutations. We synthesized three MAGE oligonucleotides, 5'-T*A*TATACCGTCACTTGCAAACTGCGAGTTCGCTGGCAGCGTCCTGGCTAGCATAGTACCTAGGACTGAGCTAGCCGTCAATCACCGCAGAGTTCCTGATTTGCCACCGTCCAGCAGCTCAAAACC, 5'-A*C*GACGCTTAGACACGTTTGTCCTCCAGGGAGGGAAAAAATGATTGCTAGCACAGTACCTAGGACTGAGCTAGCCGTCAACTAGTGGGACAAAAAGATAAAACTCAGGCGGTCGAACGACCGCCT, and 5'-C*A*ATGACGATATACTTTGAGCGCATTTTTTTCCTTAAGCACTTTCGCTAGCATTGTACCTAGGACTGAGCTAGCCGTCAATTACTGGGCATTCTTCTCTTTTAGTACCTTGAGATAATCCTGCAC and performed a single CRISPR/Cas9-selection-based editing cycle using pF20Cas and a nonrepetitive sgRNA plasmid (based on pINTsg (Supplementary Data 1)), expressing the 5'-TTCCTTAAGCACTTTCTTAC and the 5'-ATCAGGAACTCTGCGGTGAC guides. Following editing, the fastest growing variants were identified, validated using whole genome sequencing, and used in follow-up integration steps.

**Segment 22**: Based on translation initiation rate (TIR) predictions and comparisons between the 57-codon genome and MDS42, we identified the *lolC-lolD-lolE* operon as a potential issue in Segment 22. Therefore, we resynthesized Segment 22 using our updated workflow. Segment 22_3 was integrated using the standard SynOMICS workflow in MDS42 Δ*recA* containing Segment 20, 21, 23, 24, 25, 26, 27, 28, and 29.

**Segment 23**: We integrated Segment 23 in MDS42 Δ*recA* containing Segment 21 using our standard SynOMICS method. However, the resulted colonies displayed slow growth, therefore we performed five genome editing cycles using a mixture of all DIvERGE and MAGE oligos targeting all potential issues in Segments 21 and 23 (Supplementary Data 1). Following five genome editing cycles and growth-based selection, colonies were plated onto Lysogeny broth Lennox (LBL) agar plates. Fast-growing colonies were subjected to whole genome sequencing and used to integrate the following segment.

**Segment 24**: We integrated Segment 24 in MDS42 Δ*recA* containing Segment 21 and 23 using our standard SynOMICS method.

**Segment 25**: We integrated Segment 25 in MDS42 ∆*recA* containing Segment 20, 21, 23, and 24 using our standard SynOMICS method. However, we identified a missense mutation due to DNA synthesis error in the growth-essential *topA* that we hypothesized to be deleterious, and therefore we corrected this mutation using ssDNA recombineering using 5'-C*C*GCGAACCATATTTACCGCGTCCTGTGACAGGTTAGTGGAGTCGGTACGCATGTAAGTGATATAGCCTGCTTCATAAAGACGCTGCGCC, before integrating Segment 25 with our standard workflow.

**Segment 26**: We integrated Segment 26 in MDS42 ∆*recA* containing Segment 20, 21, 23, 24, and 25 using our standard SynOMICS method.

**Segment 27**: We integrated Segment 27 in MDS42 ∆*recA* containing Segment 20, 21, 23, 24, 25 and 26 using our standard SynOMICS method.

**Segment 28**: We integrated Segment 28 in MDS42 ∆*recA* containing Segment 20, 21, 23, 24, 25, 26, and 27 using a modified SynOMICS method. As Segment 28 contains the *dif* (deletion-induced filamentation) site necessary for chromosome segregation during cell division in *recA*(+) cells[10], we modified the SynOMICS deletion cassette to integrate an ectopic *dif* site immediately after the gentamicin deletion cassette (28GentRC-2-27r29w, Supplementary Data 1). Following the deletion of the parental copy, the transformation of pINTsg eliminated the ectopic *dif* site and resulted in the scarless integration of this segment.

**Segment 29**: We integrated Segment 29 in MDS42 ∆*recA* containing Segment 20, 21, 23, 24, 25, 26, 27, and 28 using the standard SynOMICS method, however a deletion cassette with only 100 bp external genomic homologies was used. Due to the low growth rate of the resulted strain, we performed 40 days of ALE to increase its fitness.

**Segment 30**: We integrated Segment 30 in MDS42 ∆*recA* containing Segment 31 and 34 using the standard SynOMICS method. Segment 30 contains the terminus of the genome, which we hypothesized might hinder the deletion of the chromosomal copy, but we successfully obtained multiple slow-growing variants that lost the parental copy of Segment 30, demonstrating the stringent selection of SynOMICS deletion step. We observed the transposition of a Tn10000 transposon into *rstB*, and we eliminated this insertion using pF20Cas-based CRISPR/Cas9-assisted oligo-recombineering with 5'-G*T*TTTGTGGGCGATCGAGTCGGGCATAAGTCAGCAGCTCTTCAATTAGAGCTTCAAGTTGTGAGATATCACGATTTAGCGCCTGGGATTC as MAGE oligo and pCRISPRM-Tn4 as crRNA plasmid.

**Segment 31**: We integrated and corrected the DNA synthesis error of Segment 31 in MDS42 ∆*recA* containing Segment 34 using the standard SynOMICS method. Segment 31 contained a 3127 bp deletion in its *ydiJ* gene eliminating most of this gene and the *sodC-ydhL* region contained multiple nonrecoded, forbidden codons due to DNA synthesis errors. We corrected these errors by appending the recoded version of *sodC-ydhL* and *menI-ydiJ* to the two ends of the deletion cassette as terminal segment homologies, in total 5840 bp of appended, and used this modified cassette (31GentRC-2-30w32w, Supplementary Data 1) to replace the parental copy.

**Segment 32**: We corrected the design and DNA synthesis errors and integrated Segment 32 in MDS42 ∆*recA* containing Segment 30, 31, 33, 34, and 35 using our standard SynOMICS method. Our initial attempts to obtain SynOMICS parental-copy-deleted variants failed and therefore we troubleshot Segment 32 using multiple rounds of editing. We traced back the nonviability of this segment to a 1 bp deletion causing frameshift in the growth-essential *rpmI*, and, based on Cappable-seq results, to recoding changes in intragenic promoters driving the expression of the growth-essential *infC-rpmI-rplT*. Furthermore, we hypothesized that, based on our previous results[11] and our computational translation initiation rate prediction (**Supplementary Figure 9**), that the expression level of the growth-essential *pheS-pheT-ihfA* operon is reduced. We also identified a missense mutation in the growth-essential *thrS* (V338I), near the active site of this enzyme. Therefore, we first repaired the 1 bp deletion of *rpmI* using ssDNA recombineering with oligo 5’-T*T*TTAAGCACAAGCACGCTAACCTGCGTCACATTCTGACCAAAAAAGCGACCAAACGTAAACGTCATCTCCGGCCGAAAGCCATGGTTTC. Next, we increased the expression of the *pheS-pheT-ihfA* operon by implanting a novel, strong RBS (5’- GAGGAGGTAAGATA) in front of *pheS* using ssDNA recombineering with 5’-T*A*GGCTCTAAGTCCAACGAACCAGTGTCACCACTGACAGAGGAGGTAAGATAATGTCACATCTCGCAGAACTGGTTGCCTCAGCGAAGGC and inserted a strong promoter (BBa_J23100) downstream the *pheM (*encoding the *pheST-ihfA* operon leader peptide) to increase the expression of the *pheS-pheT-ihfA* operon. The BBa_J23100 promoter insertion used oligo 5’-T*G*CTGCTATTTTCCGCTTCTTTTTTTACTTTTCGACCTGAATCCATTGACGGCTAGCTCAGTCCTAGGTACAGTGCTAGCGGAGGCTAGCGCGTGAGAAGAGAAACGGAAAACAGCGCCTGAAAG. Following the verification of these edits using whole-genome sequencing, we corrected the *thrS* V338I mutation using oligo-recombineering with 5’-

A*G*AACCGTGAATACTGCATTAAGCCGATGAACTGCCCGGGTCACGTACAAATTTTCAACC AGGGGCTGAAGTCTTATCGCGATCTGCC*G*C. Finally, we simultaneously repaired the growth-essential *infC-rpmI-rplT* operon's expression and deleted the parental copy of Segment 32 using a modified SynOMICS deletion cassette that contained a synthetic expression unit for these genes, implanted in antisense orientation to minimize unwanted recombination at the border between Segments 32 and 33, driven by a strong SLP2018-2-18 constitutive promoter and the wild-type RBS (5'-AGTCTTAAACAATTGGAGGAATAAGGT) of *infC*. The details of the deletion cassette, 32TetRC-2-31r33r, is available in Supplementary Data 1. This SynOMICS deletion cassette conferred tetracycline resistance to allow its use in cells that are gentamicin resistant. Following the integration of this segment, we performed 40 days of ALE-based troubleshooting to increase the fitness of the resulted strain.

**Segment 33**: We integrated Segment 33 in MDS42 Δ*recA* containing Segment 30, 31, and 34 using the standard SynOMICS method with a minor modification. Instead of our commonly used pINTsg integration vector, we utilized a modified deletion cassette containing modified sgRNA targets (5'- TTAATCTGGCTGTGGTCTAGACATTCCAGGCGG and 5'-GCGCAGGCTTGGATCGAGATGAAATCTCCGGGG (PAM underlined), named 33GentRC-2-32w34r, Supplementary Table 1), and a corresponding pINTsg variant that expresses the 5'-TTAATCTGGCTGTGGTCTAGACATTCCAGG and 5'-GCGCAGGCTTGGATCGAGATGAAATCTCCG as sgRNA guides.

**Segment 34**: We integrated and corrected the DNA synthesis error of Segment 34 in MDS42 Δ*recA* using the standard SynOMICS method. We hypothesized that Segment 34 is not viable due to a 12 bp deletion in the growth-essential *gapA*. We corrected *gapA* by appending the *yeaC-gapA* region of this segment to the SynOMICS deletion cassette and integrating the recoded variant while eliminating the parental copy of this segment. Following deletion, the segment was integrated using the standard workflow. We also detected the presence of a Tn1000 transposon in *nimT,* and we removed it using our pF20Cas-based CRISPR/Cas9-assisted MAGE workflow by transforming the 5'-C*T*CATCTCCCACGGCTATTCAGAAGCACAGGCGGGTTCACTGCATGGTCTACTGCAACTA GCCACAGCAGCACCCGGTCTTCTGATCCCA MAGE oligonucleotide and the pCRISPRM-Tn4 crRNA plasmid.

**Segment 35**: We integrated Segment 35 in MDS42 Δ*recA* containing Segment 30, 31, 33, and 34 using the standard SynOMICS method. However, the resulted colonies displayed slow growth and therefore we troubleshot this strain using our 2-step method. We performed five

DIvERGE- and MAGE-based editing cycles using pORTMAGE203B (Carbenicillin-R, pBBR1 ORI, with heat-inducible expression of CspRecT based on the cI857-pL expression system, derived from pORTMAGE-2[4]) with all oligonucleotides targeting all potential errors in these segments. Following diversification, cells were allowed to recover and were passaged twice with 1:100 dilution in 2×YT broth to select fast-growing variants. By spreading bacterial cells onto 2×YT agar plates, we identified the fastest-growing variants and subjected them to whole-genome sequencing. We utilized the fastest growing colony to integrate the following segment.

**Segment 36**: We integrated and corrected the design errors of Segment 36 in MDS42 ∆*recA* using the standard SynOMICS method. Cappable-seq identified the promoter of the growth-essential *pgsA* as a potential design issue. In the promoter driving the expression of *pgsA*, a recoded leucine codon overlaps with the -35 region of the promoter. We validated the reduction of *pgsA* mRNA expression from this promoter using promoter transcriptomics and by comparing the transcription of the parental and recoded version. Promoter transcriptomics indicated an 80% reduction in mRNA production from the recoded version compared the parental variant, confirming the role of *pgsA*'s promoter in the fitness issue of this segment. We attempted the SynOMICS-based deletion of Segment 36, however, in line with our hypothesis, variants that lost the parental copy of Segment 36 reverted the *sdiA-pgsA* locus to its 64-codon, parental variant. Following the SynOMICS-based integration of the segment, we troubleshot this design error using a two-step process. We first appended the entire *sdiA-pgsA* locus to the SynOMICS deletion cassette during the deletion step of Segment 45 in the same strain. In this cassette, integrated at the boundary of Segments 45 and 46, we redesigned *pgsA*'s expression to be driven by a BBa_J23116 constitutive promoter. Following the replacement and integration of Segment 45 using the 45GentRC-2-44r46r deletion cassette (Supplementary Data 1), we eliminated the incorrectly integrated *sdiA-pgsA* locus by deleting it using CRISPR/Cas9-assisted dsDNA recombineering. We performed the deletion of this locus by cotransforming into induced, pRedCas2 containing cells a dsDNA PCR amplicon containing 400 bp terminal homologies immediately outside the *sdiA-pgsA* locus, a tetracycline resistance gene, and a recoded copy of *pgsA* driven by the SLP2018-2-2113 strong constitutive promoter (Supplementary Data 1). We employed Cas9-based counterselection against the parental copy of the *sdiA-pgsA* locus using a crRNA expression array with 5'- AAAACCACTGCCCAGATGGTGGCGTTGGCA and 5'-TTTACTATGCAGGATAAGGATTTTTTCAGC as crRNAs. Edited clones were identified based on their antibiotic resistance profile (kanamycin-R, chloramphenicol-R, and tetracycline-R) and validated using colony PCR with primers 5'-CGTGGTCAGTGTTGTATTTGGTC and 5'-GATGAAACGGTCTTGGCTTTGA, and whole genome sequencing.

**Segment 37**: We integrated Segment 37 in MDS42 ∆*recA* containing Segment 36 using the standard SynOMICS method.

**Segment 38**: We integrated and troubleshot the design errors of Segment 44 in *E. coli* DH10B containing Segment 39, 40, 41, 42, and 43 using the standard SynOMICS workflow.

**Segment 39**: We integrated Segment 39 in *E. coli* DH10B using the standard version of SynOMICS but without employing CRISPR/Cas9-based counterselection against the parental copy. We detected the transposition of a Tn1000 transposon into the *mdtQ* pseudogene. Therefore, we removed this mobile genetic element using pRedCas2 and CspRecT-assisted MAGE using 5'-C*C*CGCGCCACGTCATTCACCGCTTCAACCACCGCTTTGTTGTAAGAGGCGATAGATAGGTTTGATTCGGCTTTTGCGATATCGAGATTGG as oligo and a constitutive crRNA (5'-TCTATTAATTATAAAACCGAGCTTTCCATA) expressing plasmid, based on pCRISPR (Addgene plasmid # 42875 ; http://n2t.net/addgene:42875 ; RRID:Addgene_42875)[8].

**Segment 40**: We integrated Segment 40 in *E. coli* DH10B containing Segment 39 using the standard version of SynOMICS but without employing CRISPR/Cas9-based counterselection against the parental copy. We detected the transposition of a Tn1000 transposon into *psuT* and removed this mobile genetic element using pRedCas2 and CspRecT-assisted MAGE using 5'-G*T*CGCTGACATCAGGTTGGAAAGCGTTGCTGCTGCCAGCGCCCGAAGACCAAGCTGGGCGATTTCCGGCGCGCGTTTTGGCGAAATAGCC and 5'-T*T*TACTGGCTCAACCGTTATTGCTGAATTTGAATCGCTGGAAGCAATCAACGCCCAGTTGTTGTTGCACTTTCGGATGTGTCAGGCCCAC (the latter oligo targeting the endogenous copy of Tn1000) as oligos and a constitutive crRNA (5'-TCTATTAATTATAAAACCGAGCTTTCCATA) expressing plasmid, based on pCRISPR (Addgene plasmid # 42875 ; http://n2t.net/addgene:42875 ; RRID:Addgene_42875)[8]. Edited clones were identified using colony PCR with primers 5'- TGGCGTTTCTTATCGACATCT and 5'-AATTATTGCGCTGATCAACGG, followed by validation with whole genome sequencing.

**Segment 41**: We integrated Segment 41 in *E. coli* DH10B containing Segment 39 and 40 using the standard version of SynOMICS but without employing CRISPR/Cas9-based counterselection against the parental copy.

**Segment 42**: We integrated Segment 42 in *E. coli* DH10B containing Segment 39, 40, and 41 using the standard SynOMICS workflow. Following the integration of this segment, we initiated five cycles of DIvERGE and MAGE-based troubleshooting using pORTMAGE-Ec1 (Addgene plasmid # 138474 ; http://n2t.net/addgene:138474 ; RRID:Addgene_138474)[12] and an equimolar mixture of all oligos targeting all potential issues and DNA synthesis errors in this strain. Following five genome editing cycles and growth-based selection, colonies were plated onto LBL agar plates. Fast-growing colonies were subjected to whole genome sequencing and used to integrate the following segment.

**Segment 43**: We integrated Segment 43 in *E. coli* DH10B containing Segment 39, 40, 41, and 42 using the standard SynOMICS workflow.

**Segment 44**: We integrated and troubleshot the design errors of Segment 44 in *E. coli* DH10B containing Segment 39, 40, 41, 42, and 43 using the standard SynOMICS workflow. We successfully deleted the parental copy of Segment 44. However, the parental-copy-deleted clones displayed slow growth and our attempts to integrate this segment remained unsuccessful. We troubleshot the strain containing Segments 39, 40, 41, 42, 43 and the parental-copy-deleted Segment 44 using seven cycles of pORTMAGE203B-based DIvERGE and MAGE-based editing using an equimolar mixture of all oligos targeting all potential issues and DNA synthesis errors in this strain. Following seven genome editing cycles and growth-based selection, colonies were plated onto LBL agar plates. Fast-growing colonies were subjected to whole genome sequencing, and we attempted the integration of Segment 44. Using the optimized strain, we successfully integrated Segment 44.

**Segment 45**: Segment 45 was integrated using SynOMICS in MDS42 Δ*recA* containing Segment 9, 10A, 10B, 11, 12, 13, 14, 15, 16, 17, 18, 36, 37, 38, 39, 40, 41, 42, 43, 44, 46, 47, 48, 49, 50, 51, 52, 53, 54, 55, 56, 57, 58, and 59. We appended to the SynOMICS deletion cassette the *sdiA-pgsA* locus to troubleshoot Segment 36 during the deletion step of Segment 45. In this cassette, integrated at the boundary of Segment 45 and 46, we redesigned *pgsA*'s expression to be driven by a new BBa_J23116 promoter. Following the deletion of Segment 45 using the 45GentRC-2-44r46r deletion cassette (Supplementary Data 1), we integrated this segment using pINTsg.

**Segment 46**: We integrated Segment 46 in *E. coli* DH10B using the standard version of SynOMICS but without employing CRISPR/Cas9-based counterselection against the parental copy.

**Segment 47**: We integrated Segment 47 in *E. coli* DH10B containing Segment 46 using the standard version of SynOMICS but without employing CRISPR/Cas9-based counterselection against the parental copy.

**Segment 48**: We integrated Segment 48 in *E. coli* DH10B containing Segment 46 and 47 using the standard version of SynOMICS but without employing CRISPR/Cas9-based counterselection against the parental copy.

**Segment 49**: We integrated and corrected the design issues of Segment 49 in *E. coli* DH10B containing Segment 46, 47, and 48 using the standard SynOMICS workflow. We detected the transposition of a Tn1000 transposon in *purL* and therefore we removed this mobile genetic element using pRedCas2 and CspRecT-assisted MAGE using 5'-G*T*TCATTGCCGGGCCGCCGAGAACGACCAGCTTCGCACCGACGTTGATCTCGCCTTTTTGTACGTGATCGGCGCGAATGTTGCCGATCCC as oligo and a constitutive crRNA (5'-TCTATTAATTATAAAACCGAGCTTTCCATA) expressing plasmid, based on pCRISPR (Addgene plasmid # 42875 ; http://n2t.net/addgene:42875 ; RRID:Addgene_42875)[8]. The deletion of the parental copy of Segment 49 resulted in slow growth, and our translation initiation rate analysis identified the accidental recoding of the start codon of *tadA* as the source of the fitness decrease. To correct this issue and identify additional beneficial mutations, we initiated DIvERGE- and MAGE-based troubleshooting, targeting all DNA synthesis errors and potential design issues in this strain using pORTMAGE-Ec1 (Addgene plasmid # 138474 ; http://n2t.net/addgene:138474 ; RRID:Addgene_138474)[12] and an equimolar mixture of all oligos at 500 µM concentration. Following 5 genome editing cycles and growth-based selection, colonies were plated onto LBL agar plates. Fast-growing colonies were subjected to whole genome sequencing and used to integrate the following segment. As expected, genome editing resulted in the correction of *tadA*'s start codon and allowed us to integrate this segment.

**Segment 50**: We integrated Segment 50 using SynOMICS in MDS42 ∆recA containing Segment 9, 10A, 10B, 11, 12, 13, 14, 15, 16, 17, 18, 36, 37, 38, 39, 40, 41, 42, 43, 44, 46, 47, 48, 49, 51, 52, 53, 54, 55, 56, 57, 58, and 59. Following the integration of this segment, we performed 28 days of ALE to increase the fitness of this strain.

**Segment 51**: We integrated Segment 51 in *E. coli* DH10B using the standard version of SynOMICS but without employing CRISPR/Cas9-based counterselection against the parental copy. As the presence and active expression of RecA abolishes the counterselection efficiency of CRISPR/Cas9[13] dsDNA cuts during the deletion step of SynOMICS, performing SynOMICS in this strain required the elimination of *recA* during genome construction from Segment 51. We deleted *recA* from the integrated Segment 51 using CRISPR/Cas9-mediated ssDNA recombineering utilizing oligo 5'-C*A*AAAGGGCCGCAGATGCGACCCTTGTGTATCAAACAAGACGATTTTACTCCTGTCATGC

CGGGTAATACCGGATAGTCAATATGTTCTG and a constitutive crRNA (5’-TCGCCGTAGAAGTTGATACCTTCGCCGTAG) expressing plasmid, based on pCRISPR (Addgene plasmid # 42875 ; http://n2t.net/addgene:42875 ; RRID:Addgene_42875)[8]. Strains harboring the complete deletion of *recA* were identified with colony PCR using primers 5’-ACTGAAAGCGGCTCGTGCT and 5’-GAGTTTACGTCGCAGTTCTTG.

**Segment 52**: We integrated Segment 52 in *E. coli* DH10B containing Segment 51 using the standard version of SynOMICS but without employing CRISPR/Cas9-based counterselection against the parental copy. We detected the transposition of a Tn1000 transposon in *cysJ* and therefore we removed this mobile genetic element using pRedCas2 and CspRecT-assisted MAGE using 5'-G*T*TTGAATTTATAGTCGCCCGCGTTCACCAGCTTAACGTTCAGTTtTGCTGCTAGAAGATCATCACGAAGTGCTTCAGCAACCCGGCGCG as oligo and a constitutive crRNA (5’-TCTATTAATTATAAAACCGAGCTTTCCATA) expressing plasmid, based on pCRISPR (Addgene plasmid # 42875 ; http://n2t.net/addgene:42875 ; RRID:Addgene_42875)[8]. Edited clones were identified using colony PCR with primers 5’-CGGACTGGCAGAAAAATTCAT and 5’-GCAGAAACAGCTTTGCTTACT, and validated using whole genome sequencing.

**Segment 53**: We integrated Segment 55 in *E. coli* DH10B containing Segment 51, 52, 54, 55, 56, 57, 58, and 59 using the standard SynOMICS workflow.

**Segment 54**: We integrated Segment 54 in *E. coli* DH10B containing Segment 51, 52, and 56 using the standard SynOMICS workflow. We detected the presence of a Tn1000 transposon in *argA* following the integration of this segment and therefore we removed this mobile genetic element using pRedCas2- (*i.e.*, CRISPR/Cas9- and Lambda-Red-) assisted MAGE using 5'-A*T*TCACTGGTTCCAGGAACGTGGATTTACCCCAGTGGATATTGATCTACTGCCCGAGTCAAAAAAGCAGCTCTACAACTACCAGCGTAAA as oligo and a constitutive crRNA expression plasmid, termed pCRISPRM-Tn4 (Supplementary Data 1), containing a crRNA 5’-TGTGGCTCGTGCGATTTGTTACGGACAACG, generated from pCRISPR (Addgene plasmid # 42875)[8]. Finally, clones that lost the Tn1000 were identified using colony PCR with primers 5’-GGTACGCAGATTGTGATGGAA and 5’-GGTGATGATTGCCCGTAATGA, and validated using whole genome sequencing.

**Segment 55**: We integrated Segment 55 in *E. coli* DH10B containing Segment 51, 52, 54, 56, 57, 58, and 59 using the standard SynOMICS workflow.

**Segment 56**: We integrated Segment 56 in *E. coli* DH10B containing Segment 51 and 52 using the standard version of SynOMICS but without employing CRISPR/Cas9-based counterselection against the parental copy. Following the integration of Segment 56, we initiated DIvERGE- and MAGE-based troubleshooting, targeting all DNA synthesis errors and potential design issues in this strain using pORTMAGE-Ec1 (Addgene plasmid # 138474 ; http://n2t.net/addgene:138474 ; RRID:Addgene_138474)[12] and an equimolar mixture of all oligos at 500 µM concentration. Following five genome editing cycles and growth-based selection, colonies were plated onto LBL agar plates. Fast growing colonies were subjected to whole genome sequencing and used to integrate the following segment. We also detected the transposition of a Tn1000 transposon in *gcvP* and therefore, we removed this mobile genetic element using pRedCas2 and CspRecT-assisted MAGE using 5'-A*T*TCGGTCGCCGGTGCGCCAACCTGCGGTGGTGTCGCAAGTTGAATATCTTTCGGCACAATCTGGCCGGTCAGCGCGTTTAGCGATTGTG and 5'-T*T*TACTGGCTCAACCGTTATTGCTGAATTTGAATCGCTGGAAGCAATCAACGCCCAGTTGTTGTTGCACTTTCGGATGTGTCAGGCCCAC (the latter oligo targeting the endogenous copy of Tn1000 in *E. coli* DH10B) as oligos and a constitutive crRNA (5'-TCTATTAATTATAAAACCGAGCTTTCCATA) expressing plasmid, based on pCRISPR (Addgene plasmid # 42875)[8]. Edited clones were identified using colony PCR with external primers 5'-GATTTTCCAGCATGTTACGCA and 5'-CTGGTGAACTCAGAACCGTAT, and validated using whole genome sequencing.

**Segment 57**: We integrated and troubleshot the design issues in Segment 57 in *E. coli* DH10B containing Segment 51, 52, 54, 56, 58, and 59 using the standard SynOMICS workflow. The SynOMICS-based deletion of the parental copy was successful, however we failed to identify integrants using pINTsg. Therefore, we initiated a troubleshooting cycle after the deletion step. We performed five cycles of pORTMAGE203B-based DIvERGE and MAGE-based editing using an equimolar mixture of all oligos targeting all potential issues and DNA synthesis errors in this strain. Following five genome editing cycles and growth-based selection, colonies were plated onto LBL agar plates, and the fastest-growing variants were identified. Using the fastest-growing colony, we successfully integrated Segment 57. The resulting integrants were growing slowly, and therefore, we repeated our troubleshooting cycle. As earlier, we performed five cycles of DIvERGE and MAGE-based editing using an equimolar mixture of all oligos to target all potential issues and DNA synthesis errors in this strain. Due to the antibiotic resistance profile of the strain, we utilized pORTMAGE-Ec1 at this step. Following five genome editing cycles and selection, we utilized the fastest-growing variants to initiate the next SynOMICS segment integration cycle.

**Segment 58**: We corrected and integrated Segment 58 in *E. coli* DH10B containing Segment 51, 52, 54, and 56 using the standard SynOMICS workflow. We detected the presence of a Tn1000 transposon in the growth-essential *parE* gene of this segment (Supplementary Figure 1) and therefore we first eliminated this mobile genetic element using our standard CRISPR/Cas9-assisted oligo-recombineering workflow in MDS42 Δ*recA* cells. We utilized 5'-C*T*GAAGCTGTGGACTGGGCGCTACTGTGGCTGCCGGAAGGCGGTGAACTGCTGACCGAATCATACGTCAACCTTATCCCAACGATGCAGG as oligonucleotide and a constitutive crRNA (5'-TGTGGCTCGTGCGATTTGTTACGGACAACG) expressing plasmid, based on pCRISPR (Addgene plasmid # 42875)[8]. Edited clones were identified using colony PCR with allele-specific external primers targeting the extrachromosomal segment, and validated using whole genome sequencing. Next, we integrated the edited copy of Segment 58 using the standard SynOMICS workflow. The replacement of this segment resulted in a significant fitness drop; therefore, we performed seven cycles of pORTMAGE503B-based DIvERGE- and MAGE-based editing using an equimolar mixture of all oligos to target all potential issues and DNA synthesis errors in this strain. Following seven genome editing cycles and growth-based selection, colonies were plated onto LBL agar plates, and the fastest-growing variants were identified. Fast-growing colonies were subjected to whole genome sequencing and used in subsequent segment integration steps.

**Segment 59**: We integrated and troubleshot the design issue of Segment 59 in *E. coli* DH10B containing Segment 51, 52, 54, 56, and 58 using a modified version of the SynOMICS workflow. Based on Cappable-seq results, Segment 59 contains a design error that caused the separation of the promoter region driving the expression of the growth-essential *rpsU-dnaG-rpoD* operon, and consequently, the deletion of this segment failed using the standard workflow. We corrected this promoter issue by redesigning our SynOMICS deletion cassette to only delete the parental genome up to *rpoD*, thus leaving the parental *rpsU-dnaG-rpoD* operon intact on the genome while deleting the rest of the parental copy (59GentRC-2-58w60w, Supplementary Data 1). As expected, the use of the 59GentRC-2-58w60w deletion cassette resulted in the deletion of the parental copy of Segment 59. Next, we integrated the synthetic Segment 59 by modifying our pINTsg plasmid to—instead of the terminal end of the gentamicin resistance cassette—target the cutting of the wild-type *dnaG* using sgRNA 5'-GCCAGATCGAGCGCCATCAG. The modified plasmid version, termed pINTsg59 (Supplementary Data 1), liberated the genomic antibiotic resistance cassette, including a significant fraction of *dnaG,* allowing the removal of the parental copy of the *rpsU-rpoD* operon. The integration of Segment 59 using pINTsg59 resulted in clones that replaced the entire parental copy with the recoded variant.

**Segment 60**: We integrated and troubleshot Segment 60 in MDS42 Δ*recA* containing Segment 62, 63, 65, 66, 67, 68, and 69 using the standard SynOMICS method. The deletion and integration steps of Segment 60 were successful, but the resulted colonies displayed slow growth. Therefore, we troubleshot this strain using 27 days of ALE. Following the isolation of fast-growing colonies, we initiated the integration of the next segment using the optimized strain.

**Segment 61**: We integrated and troubleshot the design issue of Segment 61 in MDS42 Δ*recA* containing Segment 60, 62, 63, 65, 66, 67, 68, and 69 using the standard SynOMICS workflow. Our tests indicated the nonviability of Segment 61. Cappable-seq results and TIR predictions indicated multiple design errors, therefore we redesigned and synthesized a modified version of Segment 61. This new version, termed Segment 61_2, contained a repaired version of the following loci: Due to genome annotation errors, the erroneously recoded *pnp-sraG* region was repaired and recoded according to our latest genome annotation. This defective recoding impacted the expression of *pnp*, demolishing its RBS and promoter. We also deleted the synthetic strong terminator inserted in our original genome design between *infB* and *rbfA,* based on Cappable-seq results that indicated that *rbfA* and *truB* are driven by the upstream promoter of *rimP* and *metY*. We replaced the synthetic terminator with the wild-type repeat region from the same location in MDS42. Our TIR predictions indicated the decreased expression of *nusA* and *rimP,* and we troubleshot these genes by replacing the RBS of *nusA* with a synthetic strong RBS (5'- CCGAATAAGGAGTCCC) and inserting a strong constitutive promoter (BBa_J23100) upstream of *rimP*. Our Cappable-seq results also indicated recoding changes in the promoter region of *glmM* and TIR predictions indicated the perturbation of this gene's RBS. Therefore, we replaced the promoter and RBS of *glmM* with a synthetic RBS–promoter pair (5'-TTTACGGCTAGCTCAGTCCTAGGTACTATGCTAGCGTAAAACGACGGGGGGTTTG), inserted immediately downstream *folP*. Our Cappable-seq results and TIR comparisons between the synthetic and parental variant also indicated multiple errors in the *yrbG-npr* operon: Multiple intragenic promoters were impacted by recoding changes and our TIR predictions indicated decreased expression for *yrbG, kdsD, lptC, lptA, rpoN*. We troubleshot the expression of this region with a focus on growth-essential genes, by inserting a strong constitutive promoter (BBa_J23100) with a synthetic RBS (5'-GTATCGATTAAGGAGCGTCAAAC) upstream *kdsD* and synthetic strong RBS (5'-CAGGGGTACAACTGGTATC) before *lptA*. We expected that the increased expression, driven by the BBa_J23100, in combination with the endogenous intragenic promoters of this region and the increased ribosome density[14–16], would ensure the viability and proper expression of this region. We synthesized and assembled the redesigned version of this segment (Segment 61_2) using our updated segment synthesis workflow. As expected, the updated design was viable and allowed us to replace the parental copy of Segment 61 using the standard SynOMICS workflow.

**Segment 62**: We integrated and corrected a DNA synthesis error in Segment 62 in MDS42 Δ*recA* containing Segment 69 using the standard SynOMICS method. We detected the transposition of an IS1 insertion sequence 48 bp upstream of the growth-essential *rplM,* and therefore, we removed this insertion sequence using pRedCas2-based CRISPR/Cas9-assisted ssDNA-recombineering. We utilized 5'-G*G*GGTAGGTTTGCCGGACTTTGTCGTGTGAACCTCAACAATTGAAGACGTTTGGGTGTTCACCAACGTGTAACTATTTATTGGGTAAG*C*T as recombineering oligo and a constitutive crRNA expression plasmid, expressing the 5'-AAATCGGTGGAGCTGCATGACAAAGTCATC guide, based on the pCRISPR plasmid. Edited variants that lost the IS1 were identified using recoded-segment-specific primers, 5'-CAGCGGGCCTTTTGGAAG and 5'-CTGATGGACCTTTTCTATCAATCA, validated using whole genome sequencing, and integrated in MDS42 Δ*recA* containing Segment 69 using the standard SynOMICS workflow.

**Segment 63**: We integrated and troubleshot the design issue of Segment 63 in MDS42 Δ*recA* containing Segment 62, 65, 66, 67, 68, and 69 using a modified version of SynOMICS. Based on Cappable-seq results, we expected that Segment 63 cannot complement its parental copy because the promoters driving the expression of the growth-essential *rpsM-rplQ* operon lay upstream in Segment 64, and therefore, they were separated from their corresponding expression unit. Similarly to Segment 59, we modified our SynOMICS deletion cassette to only delete the parental copy of Segment 63 until *yhdN*, a non-growth-essential gene that is driven by a promoter in Segment 63. As expected, the deletion of Segment 63's parental copy until *yhdN* using 63GentRC-2-62r63w (Supplementary Data 1) could be easily performed and was viable. Finally, Segment 63 was integrated using pINTsg and the standard SynOMICS integration workflow. During integration, due to the crossover between the recoded and parental *rpsM-rplQ* operon, the *rpsM-rplQ* operon remained wild-type. We corrected this issue and resolved the separation of the *rpsM-rplQ* operon and its promoter by appending the recoded *rpsM-rplQ* operon to Segment 64.

**Segment 64**: We integrated and troubleshot the design issue of Segment 64 in MDS42 Δ*recA* containing Segment 60, 61, 62, 63, 65, 66, 67, 68, and 69 using the SynOMICS workflow. Our tests indicated the lack of Segment 64’s viability, and our earlier tests revealed the separation of a growth-essential operon (i.e., the *rpsM-rplQ* operon) in Segment 63 from its promoters in Segment 64. Furthermore, Cappable-seq results and TIR predictions indicated multiple design errors and therefore, we redesigned and synthesized a modified version of Segment 64. This new version, termed Segment 64_2, contained a repaired version of the following loci: Based on Cappable-seq results that indicated recoding changes in the intragenic promoters of *rpmJ-rplQ*, we inserted a strong constitutive promoter upstream *rpmJ* (SLP2018-1-464). Our TIR predictions indicated the decreased expression of *rpsS, rplW, rplD* in the *rpsJ-rpsQ* operon, and therefore, we increased the expression of these genes by inserting a synthetic RBS in front of them. We also increased the expression of the entire *rpsJ-rpsQ* operon, because we hypothesized that recoding changes impact essential intragenic promoters in this region based on Cappable-seq results, by inserting a strong (SLP2018-1-645[6]) constitutive promoter upstream of the *rpsJ-rpsQ* operon. Following the synthesis of Segment 64_3, we hypothesized that additional changes will be needed to restore the viability of this segment. Our Cappable-seq results indicated the recoding of the intragenic promoters driving the growth-essential *rpsL, rpsG, fusA,* and *tufA*, and therefore, we performed pF20Cas- (*i.e.*, CRISPR/Cas9 and CspRecT-) assisted oligo-recombineering to insert a strong constitutive promoter at this location. Using oligo 5’-G*A*CTATACTGATTTCGTCCGTCTTACGGTTAAGCACCCCTCGCAAATGGCCTGGTGATTAGATTTCTCATTTGAAGATTGACAGCTGAAAGTAGTGCAAATATAATCACCATCCGTAAAGCGTAAGTCGTTTATCATTGTGTGAGGACGTTTTATTACGTGTTTACGAAGCAAAAGCT*A*A and a 5’-AACGATCCCGCCATCACCAGGCCATCTGCG crRNA expressing pCRISPR variant, we inserted an SLP2018-1-290 strong, constitutive promoter upstream *rpsL*. Using the same strategy, we also inserted a synthetic constitutive promoter (SLP2018-1-561[6]) upstream of *rpmJ* to replace the SLP2018-1-464 promoter (using the 5’-G*T*CTGCACTTAAGAAGGCGAACCTGAAAGGCTACGGCCGATAATTTAACGTCTATGCATATGCTTTTGACAAGTTTATCGAATATGCATATAATGACATACCTGGTGCCGCATAACCAAGAATGGTCGCCCGAGAAGTTACGGAGAGTAAAAATGAAAGTTCGTGC*T*T oligo and the 5’-TCGGGCGACCCGCTTTAAGCATCCTGGCAC crRNA). Following these edits, we attempted the deletion of Segment 64 and *rpmJ-rplQ,* however we could not obtain viable variants containing the deletion of the parental chromosomal region. As we hypothesized that the reason for nonviability is the lack of uniform expression for ribosomal genes in the *rpsL-tufA* and *rpsJ-*

*rplQ* operons due to the presence of antisense promoters and recoding changes in promoters driving these genes, we performed a two-step deletion procedure for this segment before attempting integration. Using a modified SynOMICS deletion cassette, we first deleted the parental copy of this segment up to *gspB*, starting from Segment 65. Following deletion, we utilized the strategy of Segment 59 and integrated this segment using our standard SynOMICS workflow, however we observed the duplication of the ribosomal operon region, containing the complete recoded variant.

**Segment 65**: We integrated Segment 65 in MDS42 Δ*recA* containing Segment 62 and 69 using the standard SynOMICS method.

**Segment 66**: We integrated Segment 65 in MDS42 Δ*recA* containing Segment 62, 65, and 69 using the standard SynOMICS method.

**Segment 67**: We integrated Segment 65 in MDS42 Δ*recA* containing Segment 62, 65, 66, and 69 using the standard SynOMICS method.

**Segment 68**: We integrated Segment 65 in MDS42 Δ*recA* containing Segment 62, 65, 66, 67, and 69 using the standard SynOMICS method.

**Segment 69**: We integrated Segment 69 in MDS42 Δ*recA* using the standard SynOMICS method.

**Segment 70**: We integrated Segment 70 in *E. coli* DH10B containing Segment 77 and 78 using the standard version of SynOMICS but without employing CRISPR/Cas9-based counterselection against the parental copy. We troubleshot the fitness of the resulting strain by performing five cycles of DIvERGE and MAGE-based troubleshooting using pORTMAGE-Ec1 (Addgene plasmid # 138474 ; http://n2t.net/addgene:138474 ; RRID:Addgene_138474)[12] and an equimolar mixture of all oligos targeting all potential issues and DNA synthesis errors in this strain. Following five genome editing cycles and growth-based selection, cells were plated onto LBL agar plates. Fast-growing colonies were subjected to whole genome sequencing and used in the following steps. However, we detected the reversion of E. coli DH10B's *recA1* allele to wild-type *recA,* which in turn abolished the selection stringency of our CRISPR/Cas9-based SynOMICS system and resulted in the lack of deletability for Segment 76. We therefore deleted *recA* from this strain using CRISPR/Cas9-mediated ssDNA recombineering utilizing oligo 5'-C*A*AAAGGGCCGCAGATGCGACCCTTGTGTATCAAACAAGACGATTTTACTCCTGTCATGCCGGGTAATACCGGATAGTCAATATGTTCTG and a constitutive crRNA (5'-

TCGCCGTAGAAGTTGATACCTTCGCCGTAG) expressing plasmid, based on pCRISPR (Addgene plasmid #42875)[8]. Strains harboring the deletion of *recA* were identified with colony PCR using primers 5'-ACTGAAAGCGGCTCGTGCT and 5'-GAGTTTACGTCGCAGTTCTTG, and validated using whole genome sequencing.

**Segment 71**: We integrated Segment 71 in *E. coli* DH10B containing Segment 70, 76, 77, 78, and 80 using the standard SynOMICS workflow.

**Segment 72**: We integrated and troubleshot the design errors of Segment 72 in *E. coli* DH10B containing Segment 70, 71, 73, 74, 76, 77, 78, 79, 80, and 81 using the standard SynOMICS workflow. Our Cappable-seq experiment indicated that the promoter of the growth-essential *yidC* contains multiple recoding changes, and we hypothesized that these changes impact the expression of *yidC* and consequently, the viability of this segment. In line with our hypothesis, colonies displayed extremely slow growth following the deletion of the parental copy. We troubleshot this fitness issue by performing 30 days of ALE to evolve faster-growing variants. Following the isolation of fast-growing colonies, we succeeded with the integration of Segment 72. Adaptive laboratory evolution resulted in the duplication of the *yidX-cbrB* locus containing *yidC*, further confirming the role of the decreased expression of *yidC* in the fitness decrease of this strain.

**Segment 73**: We integrated Segment 73 in *E. coli* DH10B containing Segment 70, 71, 74, 76, 77, 78, 80, and 81 using the standard SynOMICS workflow. As Segment 73 contains the growth-essential origin of chromosome replication (oriC), we hypothesized that the SynOMICS deletion cassette must contain a copy of oriC while deleting the parental copy of this segment. We, therefore, synthesized a version of our standard deletion cassette that contains an ectopic oriC inserted directly after the gentamicin resistance gene (73GentRC-2-72w74r, Supplementary Data 1), flanked by the target sites of the sgRNAs expressed by pINTsg. In this design, the integration step liberates the entire gentamicin-R-oriC construct, allowing the scarless integration of Segment 73. The deletion of the parental copy was successful using 73GentRC-2-72w74r, and our standard pINTsg-based integration allowed us to replace Segment 73. The resulting colonies displayed low fitness. Therefore, we performed five cycles of pORTMAGE203B-based DIvERGE- and MAGE-based editing using an equimolar mixture of all oligos to target all potential issues and DNA synthesis errors in this strain. Following five genome editing cycles and growth-based selection, colonies were plated onto LBL agar plates, and the fastest-growing variants were identified based on colony size. Fast-growing colonies were subjected to whole genome sequencing. As the resulting clones were still growing slowly, we performed an additional five cycles of pORTMAGE503B-based DIvERGE- and MAGE-based editing using an equimolar mixture of all oligos targeting all potential issues and DNA synthesis errors in this strain. Following the additional five genome editing cycles and growth-based selection, cells were plated onto LBL agar plates, and the fastest-growing variants were identified. Fast-growing colonies were subjected to whole genome sequencing and used for subsequent integration steps.

**Segment 74**: We integrated and troubleshot the design issues of Segment 74 in *E. coli* DH10B containing Segment 70, 71, 76, 77, 78, and 80 using the standard SynOMICS workflow. We detected the transposition of a Tn1000 into *wzyE*, but we identified a clone that did not harbor the transposon and we used the transposon-free variant for strain construction. The deletion of Segment 74 resulted in drastic fitness decrease and whole genome sequencing indicated that the deleted clone contained multiple codon reversions in the growth-essential *rhoL-rho* operon. Our TIR comparison results between Ec_Syn57 and MDS42 indicated the increase of translation initiation rate for *rhoL* and TIR decrease for *rho*. We also detected the transposition of an IS1 element 41 bp from the start of *rhoL*. As IS elements are known to contain internal promoters and drive the transcription of genes outside their insertion point, and based on Cappable-seq results, we hypothesized that the reason for the fitness decrease is the decreased expression of *rhoL*. We troubleshot this error and reverted the nonrecoded forbidden codons while simultaneously deleting the IS element by performing CRISPR/Cas9-assisted dsDNA recombineering using pRedCas2 and amplified the recoded locus using primers 5'-C*C*TCATTTATTTGGTAAGATTGGGTGA and 5'-G*G*GCGTACAGTTATGAAACC*CTTTTTTTTC. We utilized terminally and internally phosphorothioated oligonucleotides (phosphorothioate bonds are indicated as a star within oligonucleotide sequences) as PCR primers to decrease the chew-back and nuclease digestion of the resulting ~3500 bp amplicon and enhance the full-length integration of the entire PCR amplicon using Lambda-Red dsDNA recombineering[17,18]. The co-transformation of this PCR amplicon with a multiplexed sgRNA expression plasmid, based on the nonrepetitive design of pINTsg expressing 5'-CTGCGTACTCTCCTGTGACC and 5'-TTGAAACGGCGGATTTGGCT guide sequences and a tetracycline resistance gene, and the selection of cells on chloramphenicol and tetracycline, generated multiple edited clones. We PCR amplified the entire *rhoL-rho* region from multiple, independent clones and subjected them to Sanger sequencing. Sanger amplicon sequencing identified multiple clones that reverted the *rhoL-rho* region to its 57-codon recoded design. We continued strain construction using these edited variants.

**Segment 75**: We integrated and troubleshot the design errors of Segment 75 in *E. coli* DH10B containing Segment 70, 71, 72, 73, 74, 76, 77, 78, 79, 80, and 81 using the standard SynOMICS workflow. Following segment integration, we performed 39 days of ALE to increase the resulting strain's fitness.

**Segment 76**: We integrated Segment 76 in *E. coli* DH10B containing Segment 70, 77, and 78 using the standard SynOMICS workflow. We detected the transposition of a Tn1000 transposon in the intergenic region between *glnA* and *bipA*, and therefore we removed this mobile genetic element using pF20Cas-assisted ssDNA-recombineering using 5'-A*T*ATTGGTGCAACATTCACATCGTGGTGCAGCCCTTTTGCACGATGGTGCGCATGATAACGCCTTTTAGGGGCAATTTAAAAGTTGGCAC as oligo and a constitutive crRNA (5'-TGTGGCTCGTGCGATTTGTTACGGACAACG) expressing plasmid, based on pCRISPR (Addgene plasmid #42875)[8]. Edited clones were identified using colony PCR with external primers, 5'-AATGCCTTTCCAGCCGCCAA and 5'-ATTGTTGGAGCAGCTTGTCTA, and validated using whole genome sequencing.

**Segment 77**: We integrated Segment 77 in *E. coli* DH10B containing Segment 78 using the standard version of SynOMICS but without employing CRISPR/Cas9-based counterselection against the parental copy. We detected the transposition of a Tn1000 transposon in *glpK* and therefore we removed this mobile genetic element using pRedCas2 and CspRecT-assisted MAGE using 5'-G*A*GCGCGAGTTCCGTCCAGGCATCGAAACCACTGAGCGTAATTACCGTTACGCAGGCTGGAAAAAAGCGGTTAAACGCGCGATGGCGTGG and 5'-T*T*TACTGGCTCAACCGTTATTGCTGAATTTGAATCGCTGGAAGCAATCAACGCCCAGTTGTTGTTGCACTTTCGGATGTGTCAGGCCCAC (the latter oligo targeting the endogenous copy of Tn1000) as oligos, and a constitutive crRNA (5'-TCTATTAATTATAAAACCGAGCTTTCCATA) expressing plasmid, based on pCRISPR (Addgene plasmid # 42875 ; http://n2t.net/addgene:42875 ; RRID:Addgene_42875)[8]. Edited clones were identified using colony PCR with external primers 5'-GGTTGAGCATAATACGCATGG and 5'- CGCAGTAGCAAACAATTTCCT, and validated using whole genome sequencing.

**Segment 78**: We integrated Segment 78 in *E. coli* DH10B using the standard version of SynOMICS but without employing CRISPR/Cas9-based counterselection against the parental copy.

**Segment 79**: We integrated and troubleshot the design issues of Segment 79 in *E. coli* DH10B containing Segment 70, 71, 73, 74, 76, 77, 78, 80, and 81 using the standard SynOMICS workflow. Our translation initiation rate comparison between Ec_Syn57 and MDS42 indicated the decreased expression of *rplL, nusG*, and *secE*. In line with this prediction, our deletion experiments failed to generate variants that lost the parental copy of Segment 79. Cappable-seq experiments also indicated the separation of the promoter driving the expression of *secE* and *nusG* in Segment 78 from the *secE-nusG* operon in Segment 79. We troubleshot this segment by inserting a synthetic RBS (5'-GGAGCTTAATGA) upstream *rplL* that increased gene expression for the recoded variant using pORTMAGE503B and 5'-T*G*CAACTGCTTCAATGATTTGATCTTTAGTGATAGACATTCATTAAGCTCCTTAAATTGTTCCTGAATATCAGAACAAGTTTATACGTAG as MAGE oligo in MDS42 cells, lacking *recA*. We temporarily corrected the expression of the *secE-nusG* operon by appending it to the SynOMICS deletion cassette under the control of its native promoter (79GentRC-78r80r, Supplementary Data 1). As expected, we succeeded with the deletion of the parental copy of Segment 79, however whole genome sequencing indicated the reversion of recoded codons in the *secE-nusG* operon, confirming our prediction of decreased expression level due to changes in translation initiation rate. Following the integration of the segment, we corrected the presence of these forbidden codons in the *secE-nusG* operon using CRISPR/Cas9-assisted dsDNA recombineering. We amplified the recoded full-length *secE-nusG* genes and transformed the resulted amplicon into pRedCas2-induced competent cells of the strain together with a multiplexed CRISPR/Cas9 sgRNA expression construct, expressing 5'-ATACCGAAGCTCAAGGAAGC and 5'-TCGACTTCTTTATCGCTGAT as targeting guides. We PCR amplified the entire *secE-nusG* region from multiple clones and subjected it to Oxford Nanopore sequencing. Amplicon sequencing identified multiple clones that reverted the *secE-nusG* region to its recoded form except for the first AGT serine codon of *secE*, indicating the deleterious effect of this codon's synonymous recoding. As our TIR predictions revealed the decreased expression of *secE* and we hypothesized that the decreased expression of this growth-essential gene prevents the recoding of its first AGT serine codon, we repaired this segment and recoded the AGT forbidden codon using MAGE and the insertion of a synthetic ribosomal binding site that counterbalances the 5' recoding effects of this gene. Oligo recombineering with 5'-C*T*TCCAGGCCGCGCCCTGATCCTTGAGCTTCGGTATTCGCAGACATACTAGTCCCTCCTAAACCGTTTCTACAAACATTTTCACCCCGCGATCGCGAGGCAAACCAAA*T*C that simultaneously inserts an upregulating RBS upstream *secE* and recodes the forbidden AGT serine codon as TCT, instead of TCA in our original genome design, followed by allele-specific PCR-based screen identified multiple clones with RBS insertion. Sanger sequencing of *secE* in these clones indicated the insertion of the 5’-CGGTTTAGGAGGGACTAGT RBS and the successful recoding of the AGT forbidden codon in *secE*.

**Segment 80**: We integrated Segment 80 in *E. coli* DH10B containing Segment 70, 76, 77, and 78 using the standard SynOMICS workflow. We detected the transposition of a Tn1000 transposon in *malM*; therefore, we removed this mobile genetic element using pF20Cas-assisted MAGE using 5'-G*G*AGGAGTTCGTTTTCACTTTCAGTTTCAGAAGGCCATCGGTGGTATGACGAGCAACCGGATCGGGGATATCCGGGATCGAGTTACCGAC and a constitutive crRNA (5’-TGTGGCTCGTGCGATTTGTTACGGACAACG) expressing plasmid, based on pCRISPR (Addgene plasmid # 42875 ; http://n2t.net/addgene:42875 ; RRID:Addgene_42875)[8]. Edited clones were identified using colony PCR with external primers 5’- TTGGCGAACTGACCCTGAC and 5’- TACCCCTTGCCTTTTACTGAAGA, and validated using whole genome sequencing.

**Segment 81**: We integrated and troubleshot the DNA synthesis errors of Segment 81 in *E. coli* DH10B containing Segment 70, 71, 74, 76, 77, 78, and 80 using the standard SynOMICS workflow. We utilized the deletion cassette of SynOMICS to repair the loss of the growth-essential *ssb*. We appended the *ssb* gene driven by its native promoter at the border between Segment 81 and Segment 82 (81GentRC-2-80w82r, Supplementary Data 1). As expected, the deletion of the parental copy of Segment 81 with the modified deletion cassette was successful, and we integrated this segment using our standard SynOMICS steps.

**Segment 82**: We integrated Segment 82 in MDS42 ∆*recA* containing Segment 83, 84, and 85 using the standard SynOMICS method.

**Segment 83**: We integrated Segment 83 in MDS42 ∆*recA* containing Segment 84 and 85 using the standard SynOMICS method. We detected the transposition of a Tn1000 in the BAC backbone; however, this transposon insertion did not affect the integration of this segment, as the BAC backbone degrades during the segment integration step of our standard SynOMICS workflow.

**Segment 84**: We integrated Segment 84 in MDS42 ∆*recA* containing Segment 85 using the standard SynOMICS method.

**Segment 85**: We integrated Segment 85 in MDS42 Δ*recA* using the standard SynOMICS method.

**Segment 86**: We integrated Segment 86 in MDS42 Δ*recA* containing Segment 0, 82, 83, 84, and 85 using the standard SynOMICS method following the resynthesis of this segment using a modified design. Our genome reannotation based on the latest genome of *E. coli* K-12 MG1655 revealed the presence of a previously undetected and, therefore, not-recoded gene in our original design. This gene, *bglJ*, encodes the DNA-binding transcriptional regulator BglJ. As *bglJ* contained 29 forbidden codons in our original Ec_Syn57 design[11], we recoded *bglJ* to eliminate these forbidden codons and match our 57-codon genetic code. Furthermore, our Cappable-seq experiments indicated promoter-impacting synonymous codon swaps in multiple intergenic regions of Segment 86, therefore we reverted these intergenic regions to their parental form in *E. coli* MDS42. We also appended a 147 bp part of Segment 85 containing the promoter region of *idnK* that became separated from its corresponding gene in our original genome design. Our translation initiation rate comparison between Ec_Syn57 and MDS42 indicated the decreased expression of *dnaT* and *dnaC* and our Cappable-seq results indicated the presence of a recoded leucine codon in the promoter region of the *dnaT*-*dnaC* operon. We restored the expression of these two growth-essential genes by inserting a synthetic RBS sequence in front of them to upregulate their expression. Furthermore, we reverted the inserted synthetic terminator to the parental repeat region in the intergenic region between *deoC* and *deoA* because our Cappable-seq results indicated that *deoC*-*deoA*-*deoB*-*deoD* forms a single operon. We also reverted the synthetic terminator between *nadR* and *ettA* to MDS42's intergenic repeats. We synthesized the modified version using our updated segment assembly protocol and validated the updated segment design using whole plasmid sequencing. The revised version easily replaced the parental copy and allowed us to recode this segment.

##### Genome assembly

Following the SynOMICS-based genomic integration and troubleshooting of the 88 individual synthetic chromosomal segments, we initiated the assembly of the fully recoded Ec_Syn57 genome in MDS42 Δ*recA*. We merged chromosomal regions by reversing the final integration step of SynOMICS using a modified version of our pINTsg plasmid termed pFISSIONsg, resulting in the liberation of the entire recoded chromosomal region from the parental genome (**Figure 4a**). We performed individual fission experiments in MDS42 Δ*recA* containing Segment

9, 10A, 10B, 11, 12, 13, 14, 15, 16, 17 and 18; MDS42 ∆*recA* containing Segment 30, 31, 32, 33, 34, and 35; *E. coli* DH10B containing Segment 38, 39, 40, 41, 42, 43, and 44; *E. coli* DH10B containing Segment 46, 47, 48, and 49; *E. coli* DH10B containing Segment 51, 52, 53, 54, 55, 56, 57, 58, and 59; and *E. coli* DH10B containing Segment 70, 71, 72, 73, 74, 75, 76, 77, 78, 79, 80, and 81. We reversed the integration step of SynOMICS by first delivering a fission BAC containing terminal homologies to the recoded region (**Figure 4a**), next by delivering 3 µg / 40 µl induced fresh competent cell, a modified version of the nonrepetitive sgRNA plasmid, pINTsg, termed pFISSIONsg, resulting in the liberation of the entire recoded chromosomal region from the parental genome. The resulting double-stranded genomic cut was repaired by co-delivering 8-10 µg of a double-stranded antibiotic resistance cassette sharing 400 bp terminal homologies with the genome outside the Cas9 cut sites and by inducing Lambda-Red Exo, Beta, and Gam expression from pRedCas2. The sequence of the utilized fission BACs, pFISSIONsg plasmids, and the corresponding antibiotic resistance cassettes are listed in Supplementary Data 1. Following overnight cell recovery in 2×YT broth after electroporation, cells were plated on 2×YT agar plates containing 11 µg/ml gentamicin, 50 µg/ml kanamycin, and 20 µg/ml chloramphenicol to select for successful genome-fission events. Colonies were then screened using colony PCR with primers hybridizing immediately outside the genomic antibiotic cassette, and positive hits were validated using inverse PCR spanning the fission BAC and terminal regions of the first and last segment of the fissioned region. Finally, all fissioned variants were subjected to Illumina and Oxford Nanopore sequencing and we performed hybrid *de novo* genome assembly. Variants with the correct genome sequence were used in follow-up SynOMICS-based genome fusion experiments. Following fission, we removed the pRedCas2 plasmid from each fissioned strain by passaging cells twice in 2×YT broth at 37 °C[19], 250 rpm with 1:50,000 dilution between steps and plating cells to 2×YT agar plates before identifying colonies that lost pRedCas2 by replica plating colonies to 2×YT agar plates containing 20 µg/ml chloramphenicol. Fissioned cell lines were then electroporated with 800 ng FCas7 (pLZ119 (F-CAS7), derived from pOX38[20]) plasmid (in the case of *E. coli* DH10B containing Segment 46, 47, 48, and 49) or RK2-FCas7, and the resulted clones were validated using Illumina whole genome sequencing. Instead of conjugation, we relied on electroporation to deliver the fissioned BAC from *E. coli* DH10B containing Segment 38, 39, 40, 41, 42, 43, and 44. We purified the fission BAC using the ZR BAC DNA Miniprep Kit (Zymo Research) from 20 ml mid-exponential culture ($OD_{600}$ = 0.5-0.6). We extracted plasmids in two replicates according to the manufacturer's protocol and eluted it in 30 µl elution buffer. 10 µl of the eluted BAC DNA was then mixed with 2×40 µl freshly prepared electrocompetent cells of the recipient strain and electroporated using our standard electrotransformation protocol[5,21]. Transformants were plated and selected on 2×YT agar plates containing 22 µg/ml chloramphenicol and 100 µg/ml spectinomycin, followed by colony PCR, and Illumina and Oxford Nanopore sequencing and hybrid *de novo* genome assembly-based validation.

We delivered the fissioned recoded region encoded on the fission BAC by F-plasmid-assisted conjugation. FCas7 or RK2-FCas7 containing donor cells and pRedCas2-containing recipient cells were grown from a single colony at 32 °C aerobically in Super Optimal Broth (SOB; prepared by dissolving 20 g/l tryptone, 5 g/l yeast extract, 0.5 g/l sodium chloride, 2.4 g/l magnesium sulfate, and 0.186 g/l potassium chloride in deionized $H_2O$) or 2×YT broth containing 12 µg/ml tetracycline (in the case of the donor cells), or 20 µg/ml chloramphenicol in the case of the recipient cells. General cell growth protocols followed our earlier methods[21]. Next, 1-3 ml of the donor cell culture and 1-50 ml of the recipient cell culture were pelleted at 4,000 ×g and washed four times in 2×YT broth, prewarmed to 32 °C. Donor and recipient cells were mixed at a 10-3:1 donor-to-recipient cell ratio, pelleted at 4,000 ×g and resuspended in 200 µl 2×YT broth, and spotted on 32 °C prewarmed 90-mm-diameter 2×YT agar plates. Cells were conjugated without any movement or shaking for 4-8 hours at 32 °C and washed off from plate surfaces using prewarmed 2×YT broth. Cells were washed once with 2×YT broth and then plated on 2×YT agar plates containing 22 µg/ml chloramphenicol and 100 µg/ml spectinomycin and incubated at 32 °C until colony formation (4–14 days). Colonies were then restreaked to 2×YT agar plates containing 20 µg/ml chloramphenicol and 80 µg/ml spectinomycin, screened using MASC PCR for the presence of the newly conjugated recoded region and the parental genome, and validated using hybrid Illumina and Oxford Nanopore sequencing and *de novo* genome assembly.

The assembly of the Ec_Syn57 genome was performed in the following steps:

**Construction of MDS42 ∆*recA* containing Segment 36, 37, 46, 47, 48, and 49**: We merged the fissioned Segments 46, 47, 48, and 49 into the genome of MDS42 ∆*recA* containing Segment 36 and 37 and deleted the parental copy of Segments 46-49 using our standard SynOMICS-based genome assembly workflow.

**Construction of MDS42 ∆*recA* containing Segment 36, 37, 38, 39, 40, 41, 42, 43, 44, 46, 47, 48, and 49**: We merged and troubleshot the fissioned Segments 38, 39, 40, 41, 42, 43, and 44 with MDS42 ∆*recA* containing Segment 36, 37, 46, 47, 48, and 49 using our standard SynOMICS-based genome assembly workflow. Our attempts to deliver the fission BAC containing Segment 38, 39, 40, 41, 42, 43, and 44 using our FCas7-based conjugation method failed, and therefore, we isolated and electroporated the fission BAC into MDS42 ∆*recA* containing Segment 36, 37, 46, 47, 48, and 49. We hypothesized that the low conjugation efficiency is due to the incompatibility of the F plasmid's origin-of-replication with the matching origin-of-replication of our pYES2L-based fission BAC. Therefore, we generated a version of FCas7 on which we replaced the F plasmid origin-of-replication with the broad-host-range low copy origin-of-replication of the RK2 plasmid[22] from pSEVA221[23–25] using CRISPR/Cas9-assisted dsDNA recombineering (see Supplementary Materials for the annotated sequence of RK2-FCas7). Conjugation proceeded smoothly, and we obtained conjugants using our updated RK2-FCas7 plasmid. Following the SynOMICS-based fusion of the two recoded chromosomal parts, cells displayed extremely slow growth. We troubleshot them using our standard ALE and MAGE- and DIvERGE-based workflow. We first performed 16 days of ALE and selected the fastest-growing colony for further troubleshooting. Following the transformation of pORTMAGE203B, we performed five MAGE- and DIvERGE-based genome editing cycles using all oligonucleotides targeting all DNA synthesis errors and potential promoter issues in the strain. Following five genome editing cycles and growth-based selection, colonies were plated onto 2×YT agar plates, and the fastest-growing variants were identified. Fast-growing colonies were subjected to whole genome sequencing and used in subsequent steps.

**Construction of MDS42 ∆*recA* containing Segment 36, 37, 38, 39, 40, 41, 42, 43, 44, 46, 47, 48, 49, 51, 52, 53, 54, 55, 56, 57, 58, and 59**: We merged the fissioned Segments 51, 52, 53, 54, 55, 56, 57, 58, and 59 with MDS42 ∆*recA* containing Segment 36, 37, 38, 39, 40, 41, 42, 43, 44, 46, 47, 48, and 49 using our standard SynOMICS-based genome assembly workflow. The resulting final strain grew extremely slowly and retained an extrachromosomal copy of the fission BAC. Therefore, we troubleshot this strain by performing 32 days of ALE. Following ALE, cells were plated onto 2×YT agar plates, and the fastest growing variants were identified, subjected to whole genome sequencing, and retransformed with pINTsg to eliminate the extrachromosomal fission BAC. Finally, cells were validated using whole genome sequencing. As the fully integrated variant drastically reduced fitness, we performed an additional 23 days of ALE to restore the growth rate and selected the fastest-growing variant.

**Construction of MDS42 ∆*recA* containing Segment 9, 10A, 10B, 11, 12, 13, 14, 15, 16, 17, 18, 36, 37, 38, 39, 40, 41, 42, 43, 44, 45, 46, 47, 48, 49, 50, 51, 52, 53, 54, 55, 56, 57, 58, and 59**: We merged and troubleshot the fitness of the fissioned Segments 9, 10A, 10B, 11, 12, 13, 14, 15, 16, 17, and 18 with MDS42 ∆*recA* containing Segment 36, 37, 38, 39, 40, 41, 42, 43, 44,

46, 47, 48, 49, 51, 52, 53, 54, 55, 56, 57, 58, and 59 using our standard SynOMICS-based genome assembly workflow. Following the SynOMICS-based deletion of the parental copy of Segments 9, 10A, 10B, 11, 12, 13, 14, 15, 16, 17, and 18, we performed 45 days of ALE to recover the fitness of the parental-copy-deleted strain. The deletion step was first performed using the deletion cassette from the fission step and later replaced with a tetracycline marker with flanking internal homologies before pINTsg-based integration (Supplementary Data 1). As the resulting integrants took ten days to form colonies on 2×YT agar plates, we performed 61 days of ALE, followed by an additional 43 days of ALE using two selected clones from the previous ALE step, before selecting the fastest-growing variant after the second ALE cycle for the integration of Segment 50. As Segment-50-integrated clones were growing slowly, we performed an additional 28 days of ALE before we integrated Segment 45 using our standard SynOMICS workflow.

**Supplementary Figure 1.**

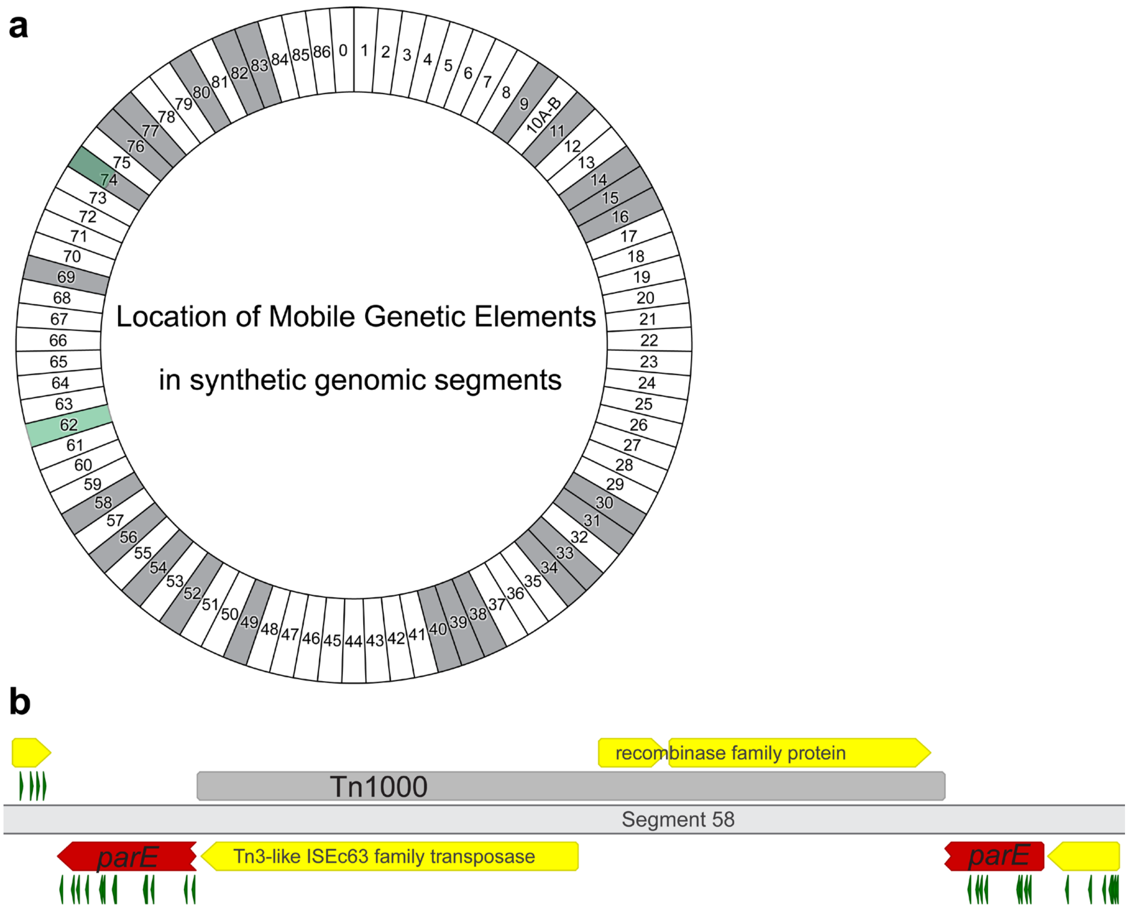


**Supplementary Figure 1. Frequent mobile genetic element transposition into synthetic genomic segments during genome synthesis.** We detected the transposition of *E. coli* DH10B's endogenous Tn1000 (γδ)[26,27] (marked in gray) and the transposition of endogenous Insertion Sequences (ISes, marked in green) into multiple synthetic chromosomal segments following DNA synthesis. These mobile genetic elements frequently disrupted growth-essential genes, leading to the nonviability of the synthetic segment. (**a**) Figure shows the location of the detected mobile genetic elements on the chromosomal map of Ec_Syn57. (**b**) Figure shows the location of the Tn1000 transposon in the growth-essential *parE* of Segment 58, based on whole genome sequencing data.

**Supplementary Table 1.**

| | SynOMICS | REXER and GENESIS | SIRCAS | CONEXER and CGS |
|---|---|---|---|---|
| **Sequence-based counterselection against parental copy** | Yes, CRISPR/Cas9-based. | No | No | No |
| **Maximum target size demonstrated in a single step** | 451 kbp | 136 kbp | 25 kbp | 136 kbp |
| **Maximum cumulative target size demonstrated** | 4 Mbp | 4 Mbp | 200 kbp | 500 kbp |
| **Identification of design issues** | Multiplexed multi-omics-based identification. | Based on the detection of not-edited chromosomal locations following experiments. | Not demonstrated. | Based on the detection of not-edited chromosomal locations following experiments. |
| **Troubleshooting design issues and DNA synthesis errors** | Multiplexed multi-omics-based identification and multiplexed MAGE-, DIvERGE-based, and non-hypermutator ALE-based troubleshooting. | Segment-resynthesis or recombineering-based correction (non-multiplexed), or hypermutator ALE. | Not demonstrated. | Design issue correction not demonstrated. Sequence correction is based on singleplex retron-editing. |

**Supplementary Table 1. Comparison of existing prokaryotic genome construction methods**. For a detailed description of REXER, GENESIS, SIRCAS, CONEXER, and CGS, see Refs. [11,28–30]. Our analysis does not include CAGE[31,32] and the previously developed flippase- (Flp-POP cloning) and Cas9-based chromosome fission/fusion-based approaches[33–35], as these methods did not demonstrate the assembly or debugging of synthetic chromosomal segments.

**Supplementary Table 2.**

| | **REXER-based recoding of *mraZ-ftsZ*** (Ref[36]) | **SynOMICS-based recoding of Segment 2** (this work) |
|---|---|---|
| Length of the target region | 20 kbp | 45 kbp |
| Number of genes' recoding attempted | 15 | 37 |
| Number of TTG and TTA codons targeted for recoding | 157 | 308<br>(including 157 in *mraZ-ftsZ*) |
| Number of all codons targeted for recoding | 157 | 585 |
| Outcome | No recoded variant identified. | Slow-growing recoded variants identified. |
| **Recoding Efficiency (%)** | **0** | **92** |

**Supplementary Table 2. SynOMICS achieves challenging recoding schemes.** The details of the previous, REXER-based recoding of *mraZ-ftsZ* in *E. coli* MDS42 is based on Reference[36]. Recoding efficiency was calculated as the fraction of assayed clones containing only the recoded allele following the elimination of its wild-type parental copy.

**Supplementary Figure 2.**

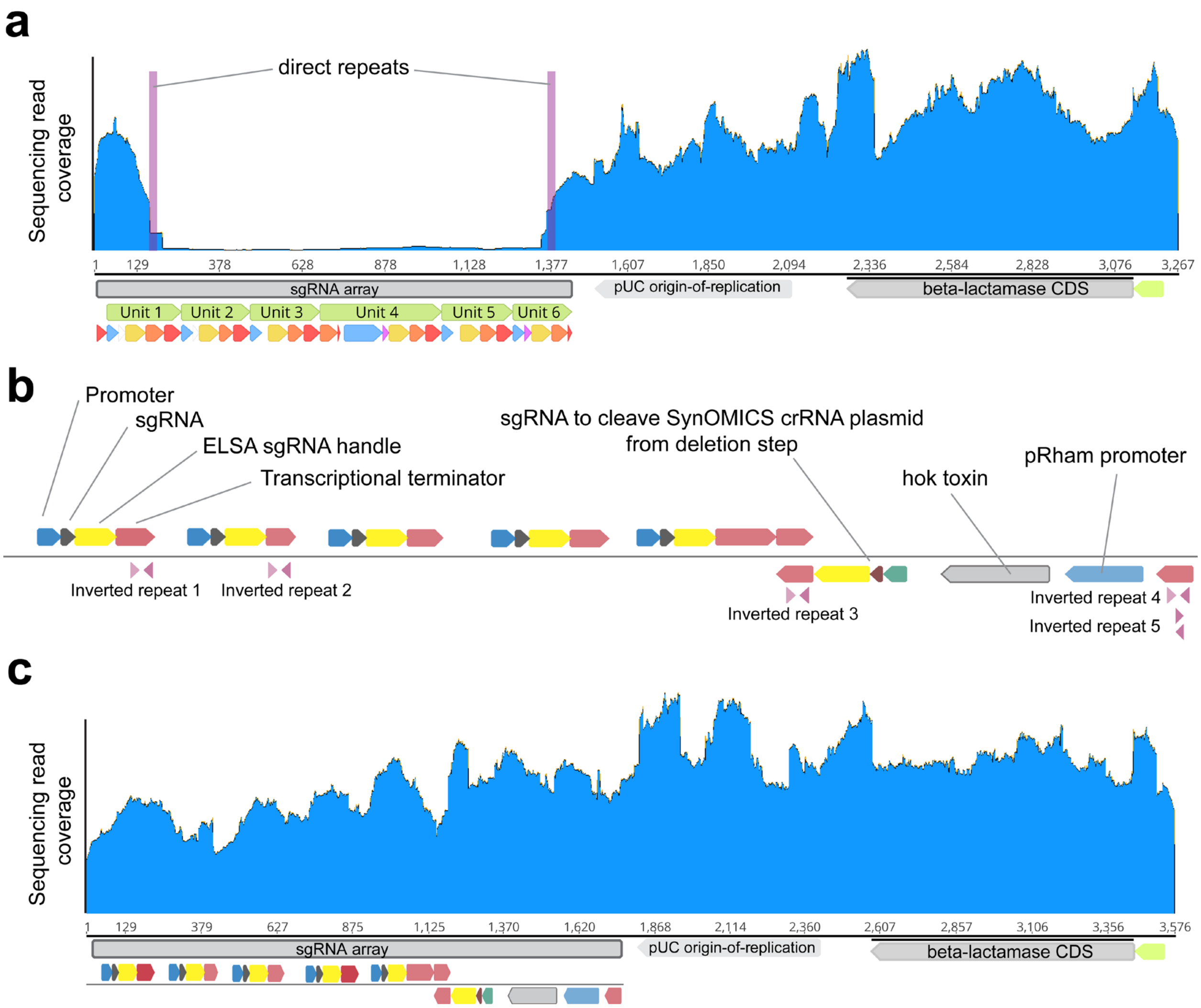


**Supplementary Figure 2. Optimization of the SynOMICS integration plasmid toward increased stability and efficiency.** (**a**) Illumina sequencing read coverage along an early version of pINTsg that contains a 14 basepairs-long intramolecular repeat leading to frequent recombination between sgRNA Unit 1 and Unit 6 during SynOMICS' integration step, and consequently, the loss of sgRNA expression for Units 2, 3, 4, 5, and 6, and the disruption and efficiency-decrease of the SynOMICS workflow. (**b**) The structure of the nonrepetitive, optimized pINTsg 6-plex sgRNA expression unit that devoids intramolecular direct repeats longer than 11 nucleotides. Short, inverted repeats (*i.e.*, Inverted repeat 1-5) in pINTsg localize in terminator structures, do not share sequence homology between repeats, and do not lead to unwanted recombination events. (**c**) Illumina sequencing read coverage along the optimized pINTsg confirms plasmid stability. The sequence of pINTsg is available in Supplementary Data 1.

**Supplementary Figure 3.**

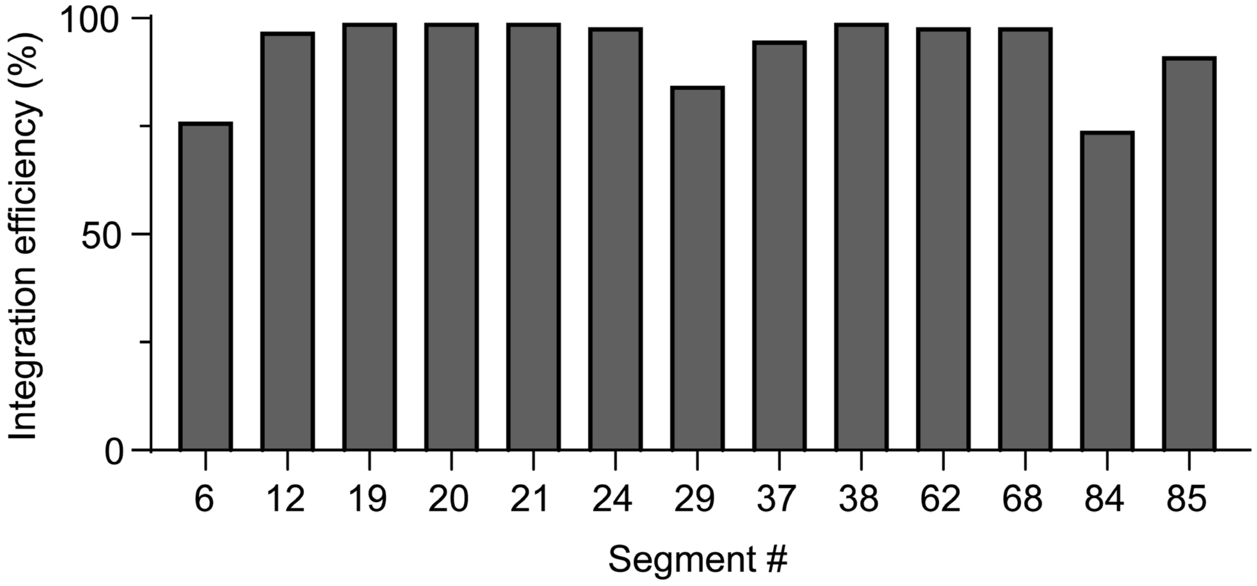


**Supplementary Figure 3. Efficient synthetic segment integration using SynOMICS.** Figure shows segment integration efficiency using the optimized pINTsg plasmid. Segment integration efficiencies were assessed based on colony-Polymerase Chain Reactions (colony-PCRs) of 34–96, randomly picked colonies following the integration of the corresponding segment. Segment integration experiments and PCR assays were performed once (*n*=1). Source data and experimental details are provided at the end of the Supplementary Material.

**Supplementary Figure 4.**

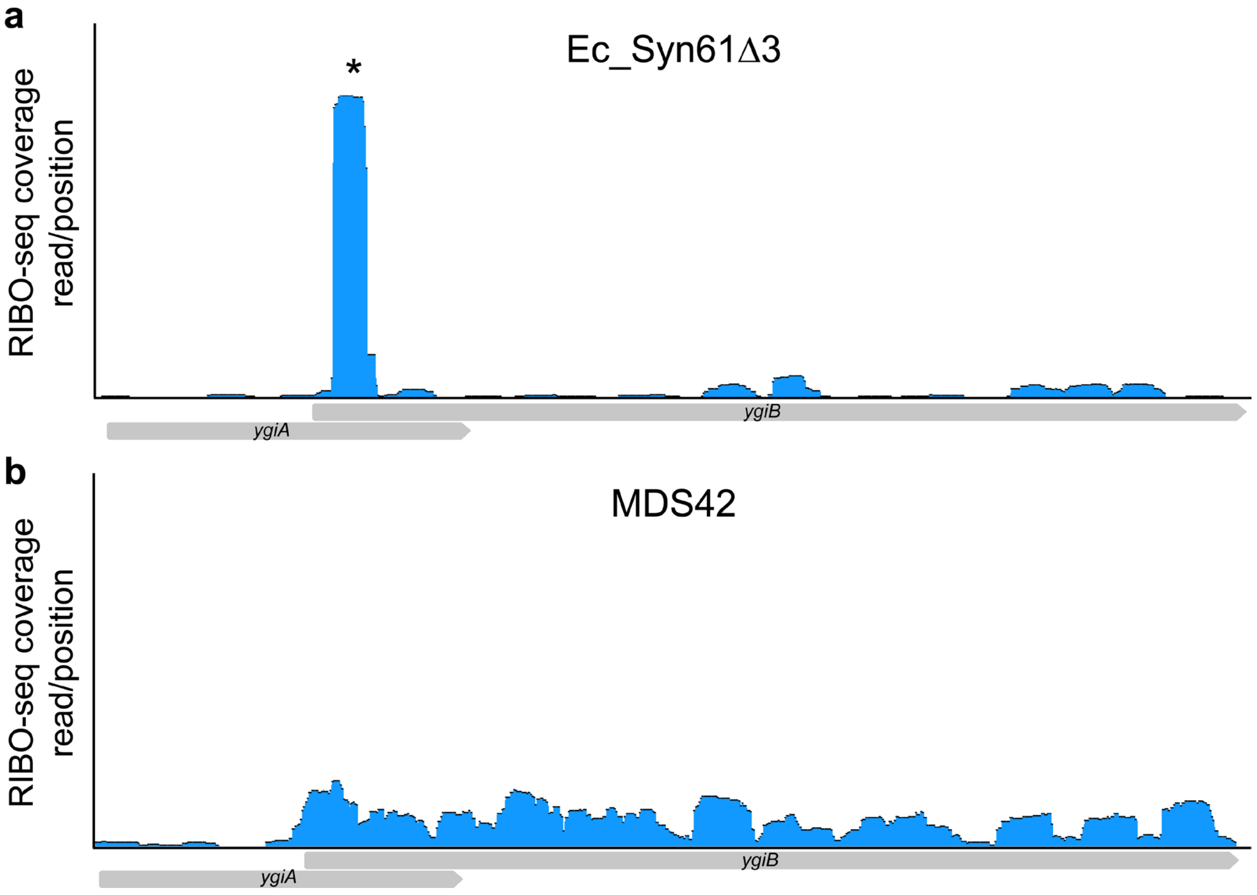


**Supplementary Figure 4. Ribosomes stall at forbidden codons present in Ec_Syn61Δ3 due to genomic annotation errors prior to genome design.** The figure shows ribosome footprint coverage based on Ribo-seq along *ygiA* and *ygiB* in Ec_Syn61Δ3 lacking $tRNA^{Ser(UGA)}$ and $tRNA^{Ser(CGA)}$ needed to translate TCA and TCG codons, respectively (**a**), and in the parental *E. coli* MDS42 bearing the unaltered, canonical genetic code (**b**). The updated annotation of *ygiB* indicated that the start codon of *ygiB* is 96 bp upstream compared to its original location used during the design of Ec_Syn61Δ3's genome, leading to the presence of an unassigned TCG codon (marked with *). Ribo-seq data was collected in three independent replicates; figure shows ribosome footprint coverage based on a representative sample.

**Supplementary Figure 5.**

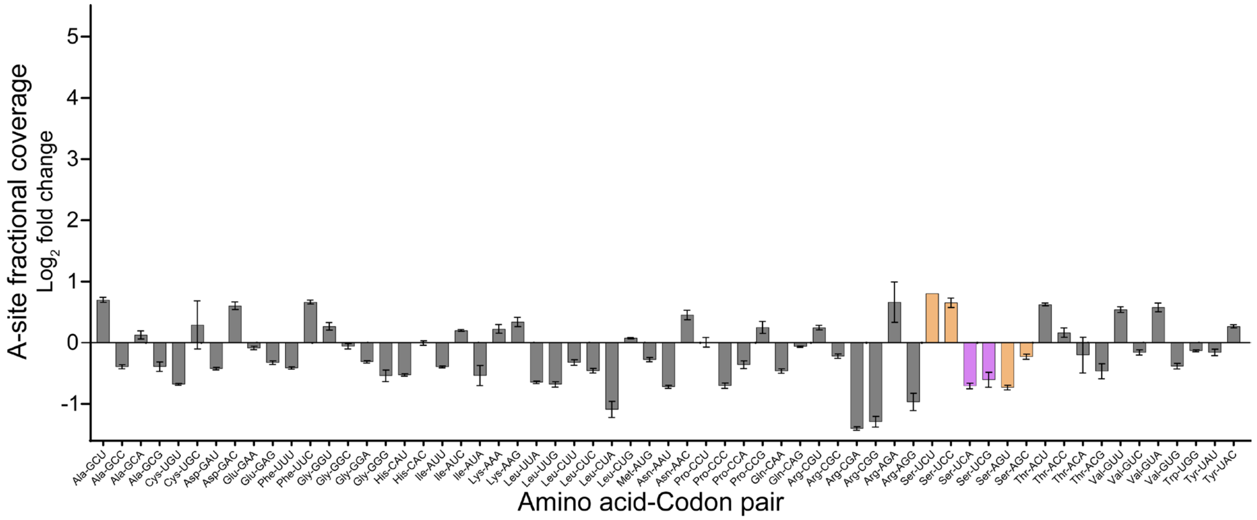


**Supplementary Figure 5. Ribosome A-(-aminoacyl-)-site coverage of sense codons in *E. coli* MDS42.** The figure shows the fold-change of relative A-site coverage of sense codons in the unaltered genetic code of *E. coli* MDS42, compared to the frequency of each codon in analyzed genes. Ribo-seq data was collected in three independent replicates; error bars show standard deviation based on *n*=3.

**Supplementary Figure 6.**

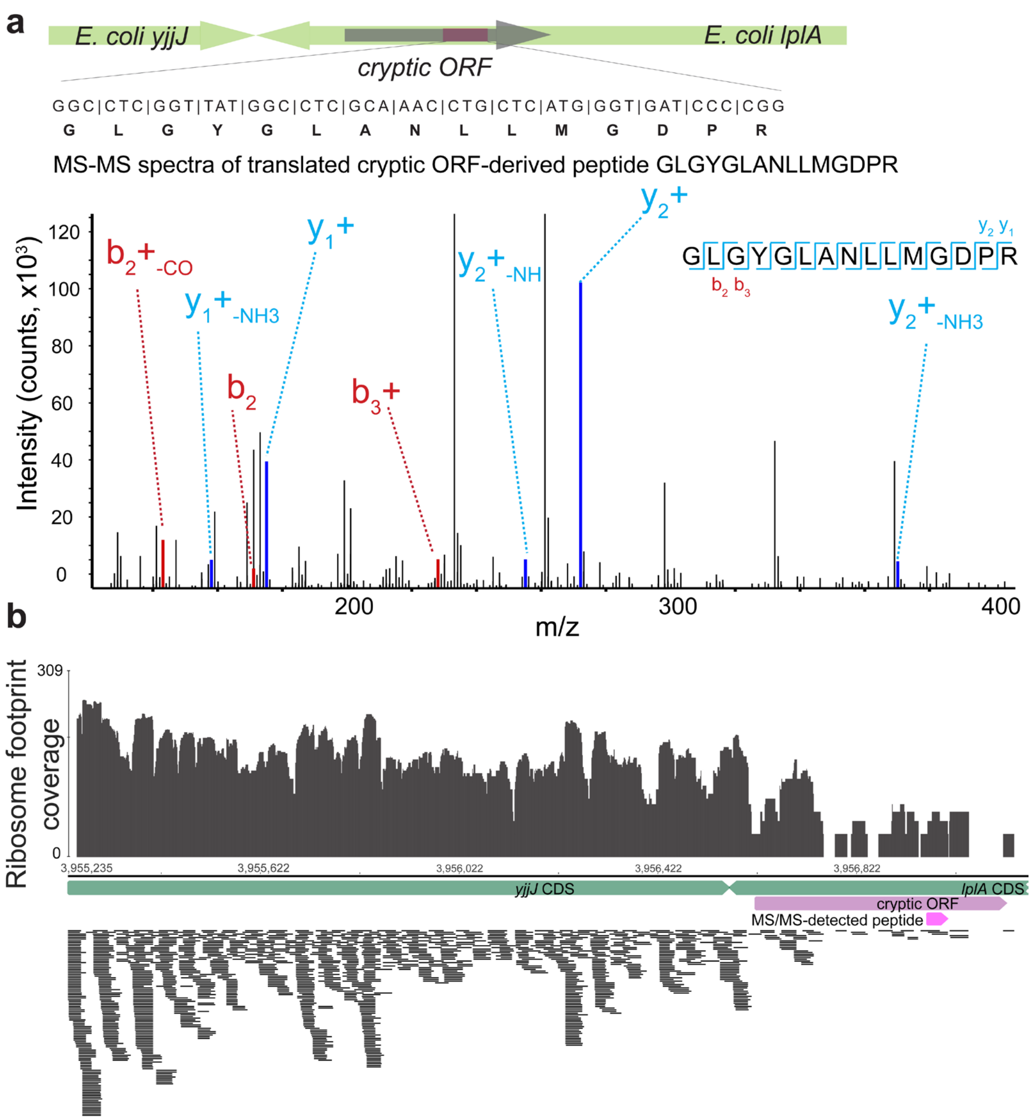


**Supplementary Figure 6. Proteomics- and ribosome-profiling-based detection of cryptic translated ORFs in *E. coli* MDS42.** (**a**) The detection of the translated cryptic intragenic ORF of *lplA* in *E. coli* MDS42 based on MS/MS data. The figure shows the amino acid sequence and MS/MS spectrum of the detected cryptic-ORF-derived peptide and its coding sequence. MS/MS data were collected once. (**b**) Figure shows Ribo-seq ribosome footprint coverage, in forward orientation (based on genomic coordinates), at the endogenous *yjjJ* and *lplA* of MDS42 and the MS/MS-detected cryptic ORF. Magenta marks the MS/MS-detected cryptic translated ORF and the MS/MS-detected cryptic ORF-derived peptide. Ribo-seq experiments were performed in three independent replicates; figure shows a representative sample.

**Supplementary Figure 7.**

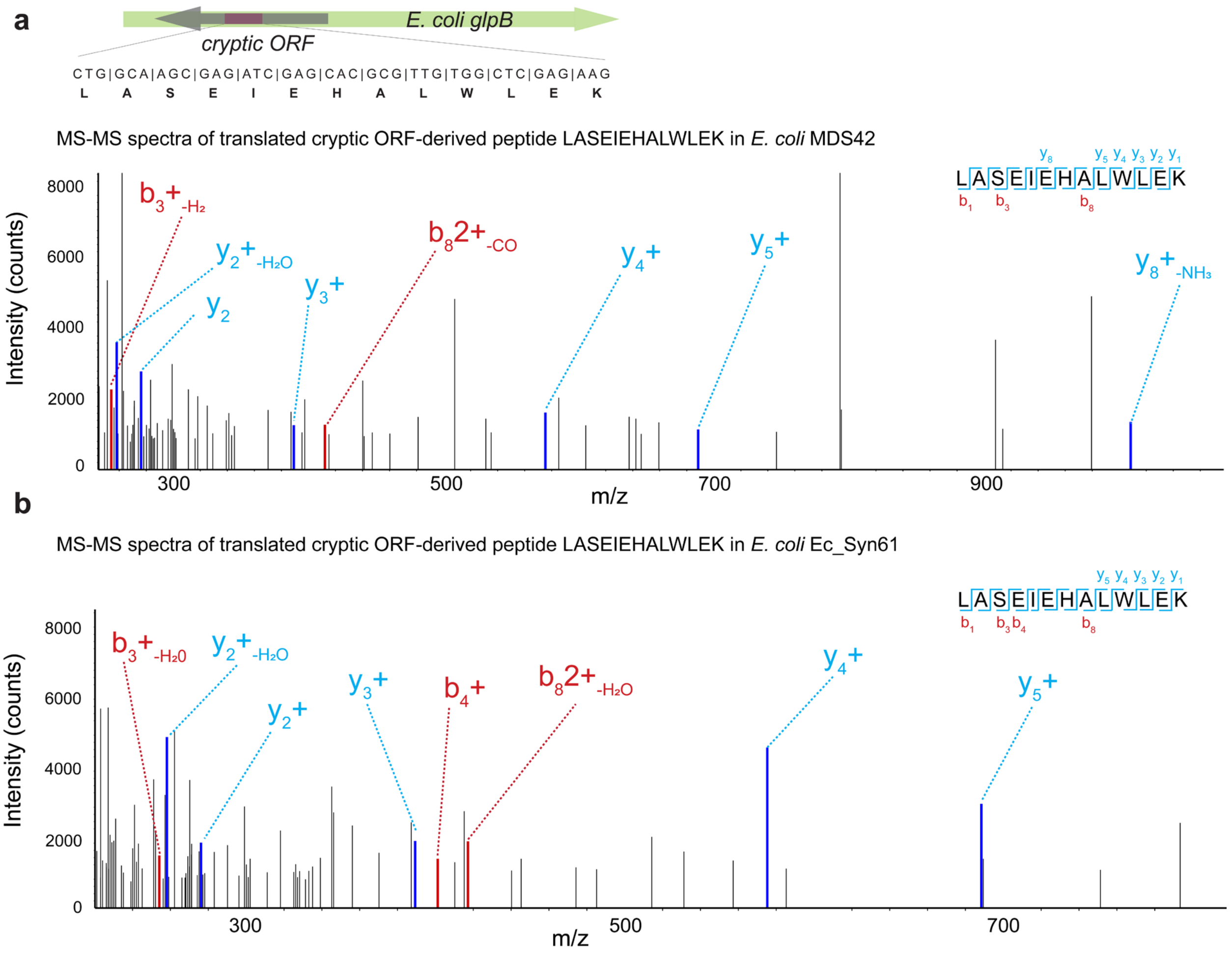


**Supplementary Figure 7. Proteomics-based detection of a cryptic translated ORF in *glpB* of *E. coli* MDS42 and Ec_Syn61.** The amino acid sequence of the translated cryptic antisense ORF in *glpB* was confirmed by tandem mass spectrometry in both *E. coli* MDS42 and Ec_Syn61. The figure shows the coding sequence, its location within *glpB*, and the MS/MS spectrum of the detected cryptic ORF-derived peptide. MS/MS data was collected once. The sequence of all identified cryptic ORFs is listed in Supplementary Data 8.

**Supplementary Figure 8.**

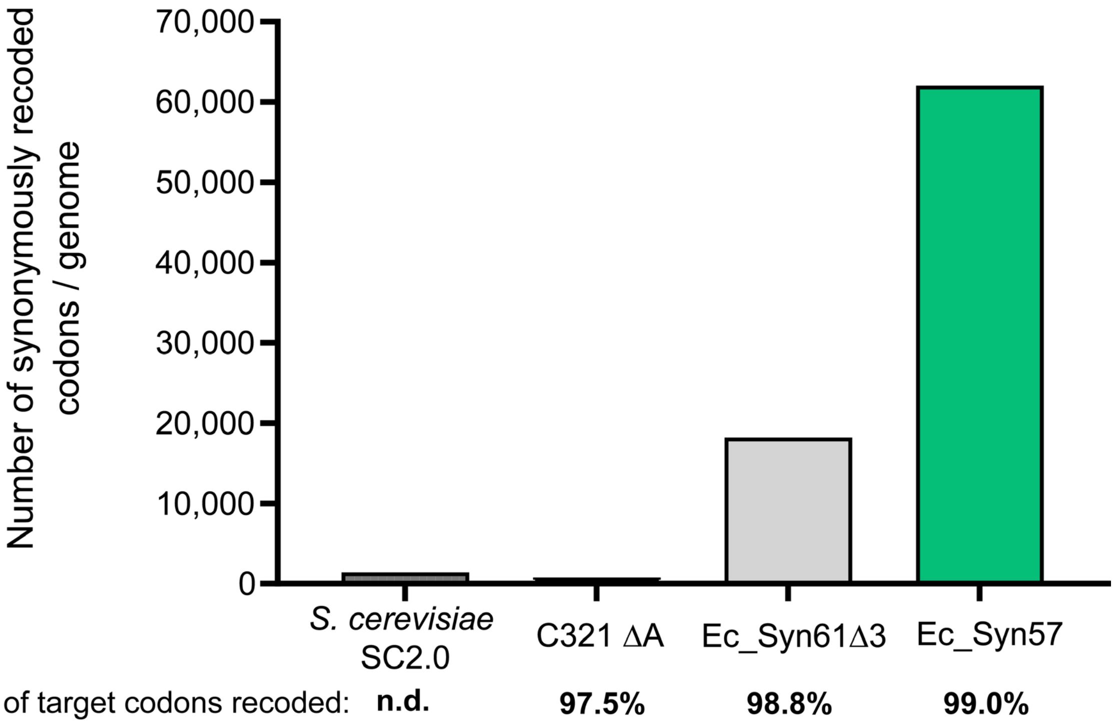


**Supplementary Figure 8. Existing recoded organisms harbor and tolerate forbidden codons.** The fraction of forbidden codons on a given genome was determined by first reannotating each genome using the most recent version of the *E. coli* K-12 MG1655 genome (GenBank accession number U00096.3), and in the case of C321.ΔA, the genome of both *E. coli* K-12 MG1655 and the λ coliphage (GenBank accession number NC_001416.1) as reference, and then updating CDS annotations across the genome to terminate at the first identified stop codon of each CDS. The fraction of forbidden codons in the case of Ec_Syn57 was calculated based on the anticipated number of forbidden codons following the outlined troubleshooting steps in this study. In the case of the *S. cerevisiae* Sc2.0 genome, *n.d.* marks *not determined*. We note that the reduction in the number of synonymously recoded codons in Ec_Syn57 compared to our 2016 design[11] (*i.e.*, *n*=62,251 codons in Reference [11]) is due to changes in gene coordinates and genome updates described in this work. The updated genome annotation of Ec_Syn57, with the erroneously recoded positions annotated as “Design_Error_Recoded_codon”, is available in the Supplementary Material of this paper.

**Supplementary Figure 9.**

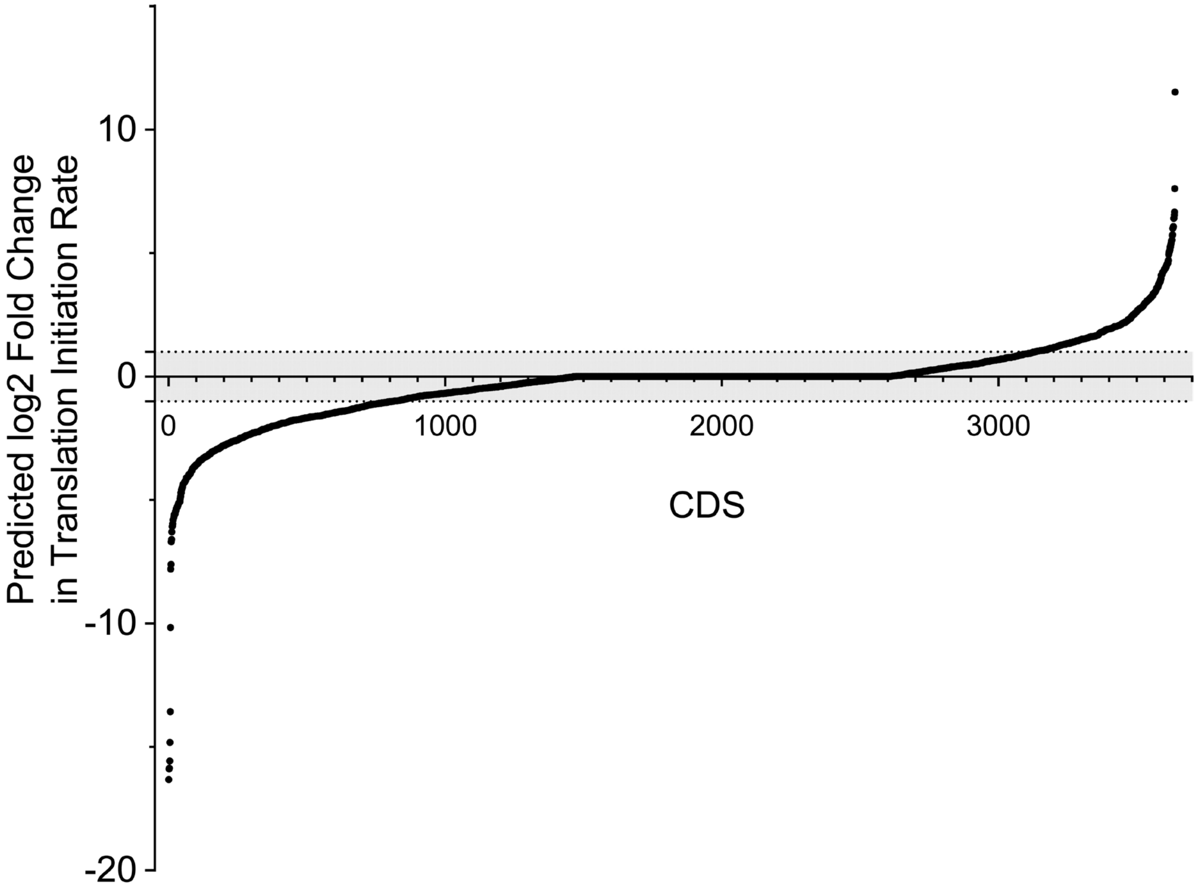


**Supplementary Figure 9. Predicted effect of synonymous recoding on translation initiation rates of protein-coding genes in Ec_Syn57.** The figure shows the predicted $\log_2$ fold-change in the translation initiation rate (TIR) of all protein-coding genes of Ec_Syn57, compared to the same gene of the parental *E. coli* MDS42. The region in grey marks a predicted fold-change in translation initiation rate lower than 2-fold, compared to the translation initiation rate of the corresponding gene in *E. coli* MDS42. The translation initiation rate of each gene was predicted using the Ribosome Binding Site Calculator Version 2.1.1 (available at *www.denovodna.com*)[14–16,37]. Source data is available within the Supplementary Material of this paper.

**Supplementary Figure 10.**

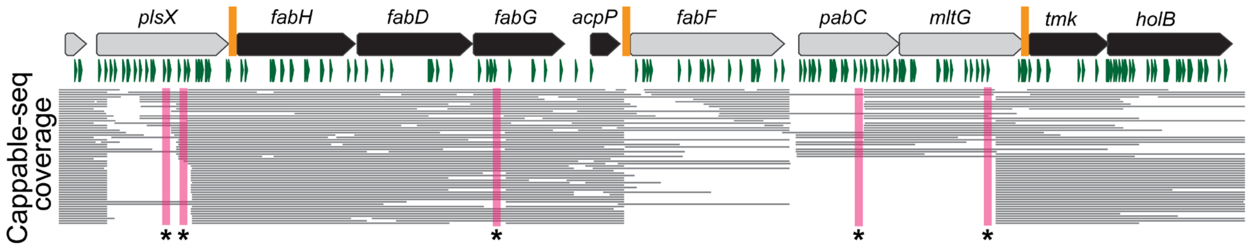


**Supplementary Figure 10. Multiplexed genome editing-based troubleshooting of Segment 21.** The figure shows the operon structure and Cappable-seq sequencing read coverage of the *plsX*-*holB* locus of Ec_Syn57's Segment 21 based on SMRT-Cappable-seq data. Genes marked in black are essential for cell growth. Green triangles mark codons synonymous recoded in Ec_Syn57; SMRT-Cappable-seq-identified promoter regions marked in magenta and with a star (*) overlap with recoding changes on the 57-codon genome and, therefore, were selected for multiplexed troubleshooting. Locations marked in orange indicate MAGE oligonucleotide target sites for the insertion of a small, focused library of constitutive promoters at each locus. The sequence of all utilized MAGE oligonucleotides and the details of the troubleshooting experiment are listed in the Supplementary Methods.

#### Supplementary Figure 11.

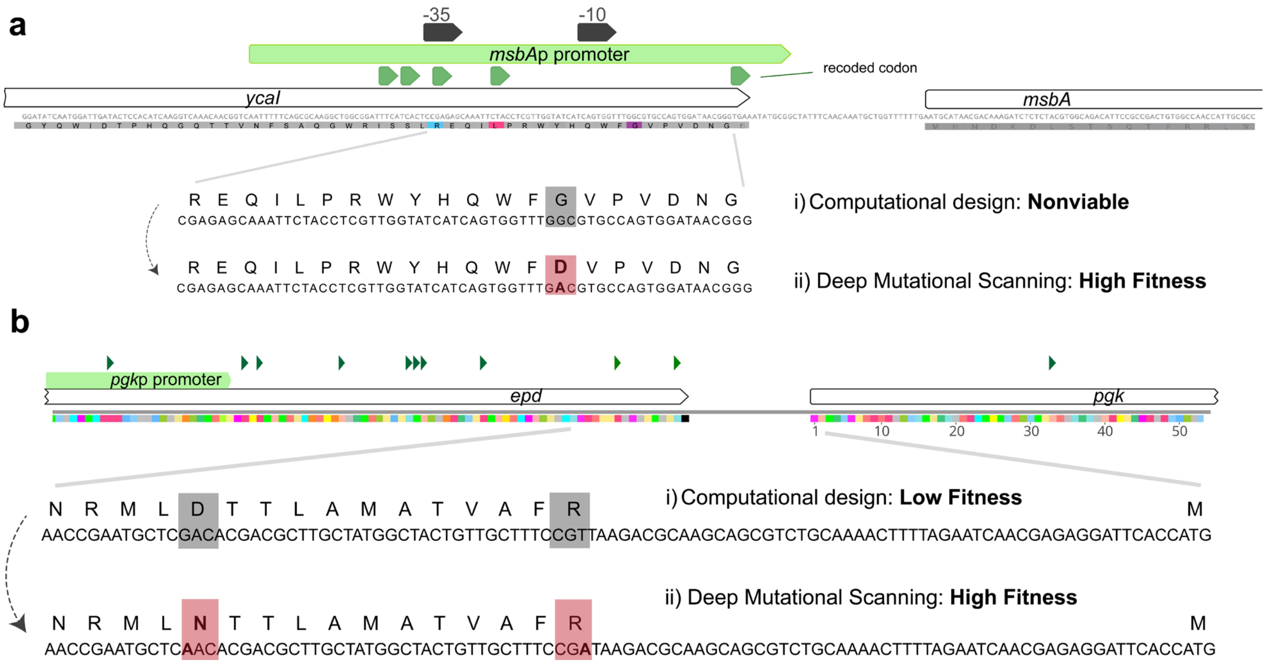


**Supplementary Figure 11. Troubleshooting synonymous recoding induced issues of *msbA* in Segment 18 and *pgk* in Segment 56 of Ec_Syn57.** Figure shows the parental allele and the targeted-mutagenesis-identified optimal variant within the promoter region of *msbA* (***a***), and the 5' UTR of *pgk* (***b***), based on Illumina whole-genome sequencing Additional details about the troubleshooting workflow are available in the Supplementary Methods of this paper.

**Supplementary Table 3.**

| Strain | Doubling time (min) | ±SD | Maximum $OD_{600}$ | ±SD |
|---|---|---|---|---|
| *E. coli* K-12 MG1655 | 27.3 | 0.973 | 1.3566 | 0.0068 |
| *E. coli* DH10B | 48.29 | 0.913 | 1.0574 | 0.0117 |
| *E. coli* MDS42 Δ*recA* | 35.37 | 0.385 | 1.3482 | 0.0155 |
| *E. coli* MDS42 Δ*recA* Segment_36-37 | 54.05 | 1.548 | 1.3581 | 0.0122 |
| *E. coli* DH10B Segment_46-49 | 44.88 | 1.944 | 0.9603 | 0.0447 |
| *E. coli* MDS42 Δ*recA* Segment_36-37_46-49 | 90.72 | 9.704 | 1.1508 | 0.0257 |
| *E. coli* DH10B Segment_38-44 | 69.24 | 22.115 | 0.463 | 0.0784 |
| *E. coli* MDS42 Δ*recA* Segment_36-44_46-49 | 147.7 | 34.706 | 0.1967 | 0.0291 |
| *E. coli* MDS42 Δ*recA* Segment_36-44_46-49 **Troubleshot** | 100.66 | 1.620 | 1.0197 | 0.0377 |
| *E. coli* DH10B Δ*recA* Segment_51-59 | 147.36 | 24.005 | 0.24 | 0.0155 |
| *E. coli* MDS42 Δ*recA* Segment_36-44_46-49_51-59 | 95.66 | 3.520 | 0.639 | 0.0213 |
| *E. coli* MDS42 Δ*recA* Segment_36-44_46-49_51-59 **Troubleshot** | 94.37 | 5.265 | 0.6656 | 0.0385 |
| *E. coli* MDS42 Δ*recA* Segment_9-18 | 91.09 | 5.891 | 0.751 | 0.0295 |
| *E. coli* MDS42 Δ*recA* Segment_9-18_36-44_46-49_51-59 | 121.78 | 6.129 | 0.4166 | 0.0280 |
| *E. coli* MDS42 Δ*recA* Segment_9-18_36-44_46-49_51-59 **Troubleshot** | 109.26 | 1.649 | 0.5193 | 0.0108 |
| *E. coli* MDS42 Δ*recA* Segment_9-18_36-59 | 123.49 | 20.151 | 0.393 | 0.0513 |
| *E. coli* MDS42 Δ*recA* Segment_1-8 | 94.02 | 8.146 | 1.0785 | 0.0626 |
| *E. coli* MDS42 Δ*recA* Segment_19-29 | 59.58 | 4.822 | 0.9795 | 0.0057 |
| *E. coli* MDS42 Δ*recA* Segment_30-35 | 63.43 | 0.793 | 1.2559 | 0.0125 |
| *E. coli* MDS42 Δ*recA* Segment_60-69 | 142.66 | 6.400 | 0.26753 | 0.0041 |
| *E. coli* DH10B Δ*recA* Segment_70-81 | 57.44 | 2.101 | 0.8212 | 0.0046 |
| *E. coli* MDS42 Δ*recA* Segment_82-0 | 51.18 | 5.548 | 1.1987 | 0.0048 |
| *E. co*li MDS42 Δ*recA* Segment_21 | 252.83 | 30.294 | 0.3874 | 0.0251 |
| *E. coli* MDS42 Δ*recA* Segment_21 **Troubleshot** | 83.12 | 0.916 | 1.3049 | 0.0176 |

**Supplementary Table 3. Fitness of parental strains and their recoded derivatives.** Doubling times were determined based on the optical density at 600 nm. Maximum $OD_{600}$ represents the maximum attainable optical density aerobically, measured at 600 nm, based on ten independent replicates. ±SD represents the standard deviation based on *n*=10 independent replicates. All measurements were performed in 2×YT broth at 37 °C. Source data is provided in this paper.

**Supplementary Figure 12.**

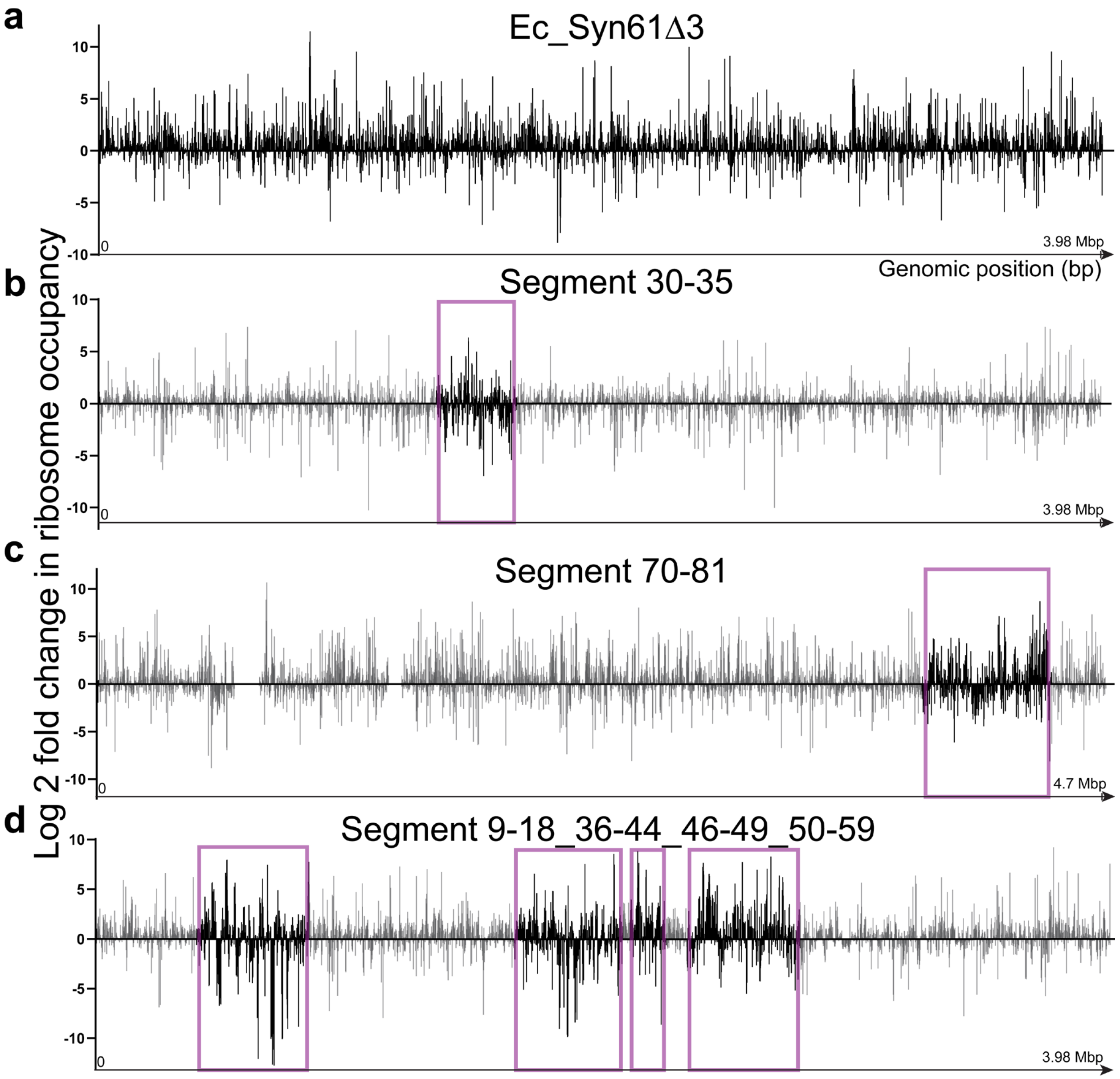


**Supplementary Figure 12. Ribosome profiling of genomes with altered genetic codes indicates large changes in translation.** The figure shows the $\log_2$ fold change in ribosome occupancy based on Ribo-seq experiments in recoded and their corresponding parental genomes. Ribo-seq data was collected in three independent replicates; bars graphs show the mean based on *n*=3. Source data is provided in Supplementary Data 3 of this paper.

**Supplementary Figure 13.**

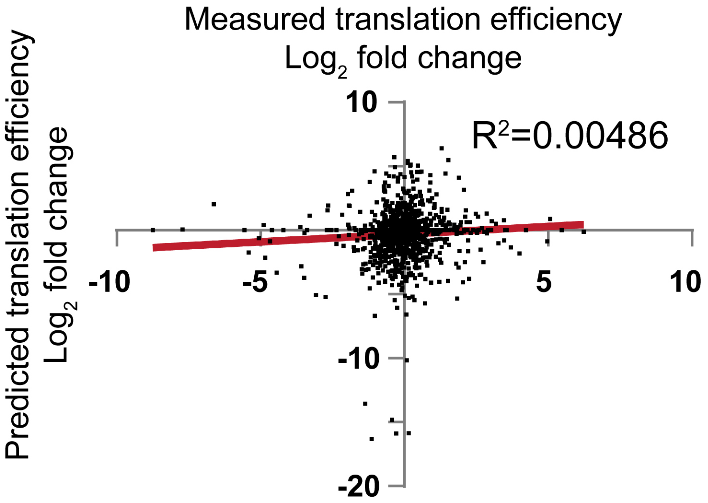


**Supplementary Figure 13. Lack of correlation between predicted and measured changes in translation within the recoded Ec_Syn57 genome.** Figure shows the correlation between the predicted relative translation initiation rate and the experimentally measured relative (*i.e.*, compared to the parental *E. coli* MDS42 derived wild-type variant) translation efficiency of genes within Ec_Syn57's Segments 9–18, 36–44, 46–49, and 51–59. Translation efficiency measurements are based on *n*=3 independent measurements. Source data is available in the Supplementary Material of this paper.

**Supplementary Figure 14.**

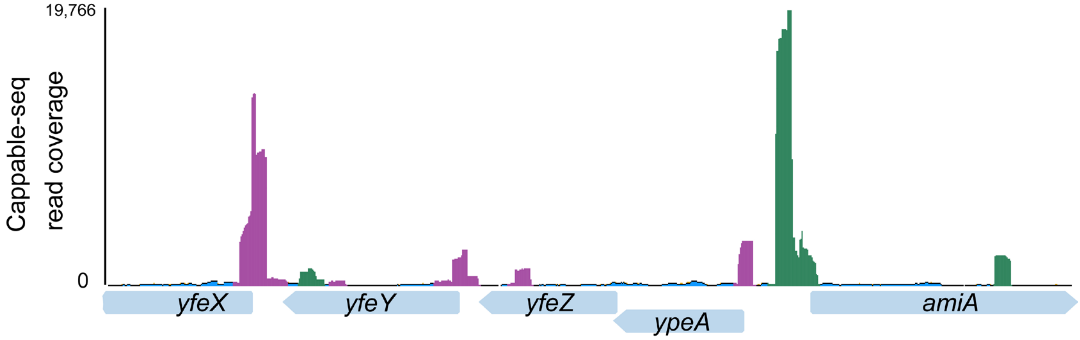


**Supplementary Figure 14. Direction and expression level of primary mRNA transcripts in *E. coli* MDS42.** The figure shows the short-read Illumina Cappable-seq read coverage within the *yfeX-amiA* locus in exponentially growing *E. coli* MDS42 cells. For the Cappable-seq coverage of the same locus in the recoded Ec_Syn57, see **Figure 5b**. Sequencing reads in magenta indicate reads in reverse orientation compared to genomic coordinates, while sequencing reads in green indicate reads in forward orientation. Cappable-seq experiments were performed in two independent replicates (*n*=2); figure shows a representative sample.

**Supplementary Figure 15.**

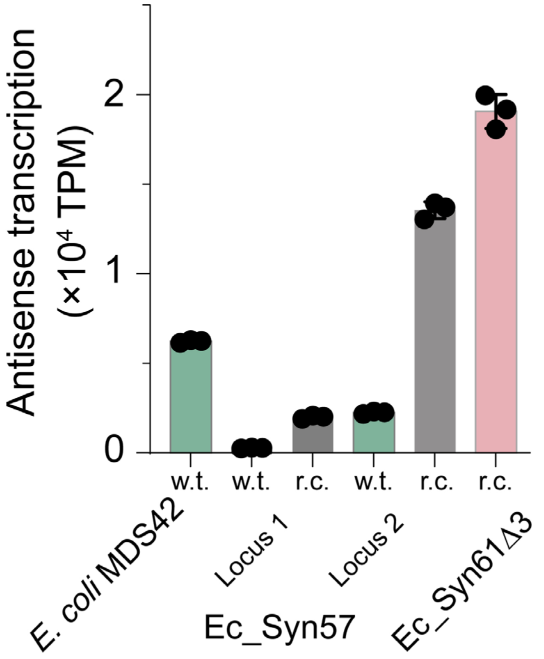


**Supplementary Figure 15. Synonymous recoding induces widespread antisense transcription.** Figure shows the level of antisense transcription within genomes utilizing the canonical 64-codon genetic code (*i.e.*, *E. coli* MDS42), a synthetic 61-codon genetic code (*i.e.*, Ec_Syn61Δ3), and two recoded, separately constructed 57-codon chromosomal regions of Ec_Syn57, based on stranded RNA-seq. w.t. indicates the parental variant of the 57-codon recoded (r.c.) synthetic region (*i.e.*, the same genomic region in *E. coli* MDS42). Locus 1 corresponds to Segments 30-35 in its *E. coli* MDS42 derived w.t. and Ec_Syn57 derived r.c. variant, while Locus 2 corresponds to Segments 9–18, 36–44, 46–49, and 51–59 in its *E. coli* MDS42-derived w.t. and Ec_Syn57-derived r.c. variant. Stranded RNA-seq experiments were performed in *n*=3 independent replicates. Bar graphs show the mean and standard deviation based on *n*=3.

**Supplementary Table 4.**

Cost estimate of genome synthesis and troubleshooting. Estimates up to our initial publication in 2016 are based on Reference [11] Supplementary Table 4.

| Step | Duration | Personnel | Materials: Materials | Materials: Cost |
|---|---|---|---|---|
| Period: 2012 - 2016 | | | | |
| Genome Design | 1 year | 1.5 FTEs | | |
| Initial DNA synthesis | 1 year | 0.5 FTE | Genomic DNA synthesis | $ 322,718 |
| | | | PCR primer synthesis for segment assembly | $ 17,000 |
| | | | MASC PCR primer synthesis | $ 7,000 |
| Initial DNA assembly and testing | 2 years | 3 FTEs | Molecular and Microbiology Reagents, Consumables | $ 72,000 |
| | | | Sequencing | $ 62,400 |
| Total cost until 2016 (FTE + Materials, based on Ref[11]) | | | $ 841,118<br>(inflation-adjusted value of $ 1,080,865 in 2024) | |
| Period: 2016 - 2024 | | | | |
| Segment DNA synthesis and assembly | 3 years | (Segment DNA synthesis and assembly were performed at CDMOs.)<br><br>0.2 FTE at Harvard | | $ 452,733 |
| Genome construction | 5 years | 1 FTE | SynOMICS deletion cassette and crRNA plasmid synthesis | $ 48,640 |
| | | | PCR primer synthesis for genome construction | $ 3,500 |
| | | | MASC PCR primer synthesis | $ 10,072 |
| | | | Reagents, consumables | $ 13,500 |
| Genome troubleshooting | 5 years | 1 FTE<br><br>(not counting FTE work at CROs, CDMOs already included in itemized cost breakdown) | DNA cassette and MAGE/DIvERGE oligonucleotide synthesis for troubleshooting | $ 36,568 |
| | | | Molecular biology reagents | $ 52,000 |
| | | | Microbial media, microbiology consumables, plasticware | $ 48,000 |
| | | | Ribosome profiling | $ 96,000 |
| | | | Proteomics | $ 15,000 |
| | | | DNA sequencing | $ 130,000 |
| | | | RNA sequencing | $ 29,000 |
| Subtotal Personnel cost between 2016 - 2024 | $ 848,000 | | Subtotal Material cost between 2016 - 2024 | $ 935,013 |
| Total cost to date | | | | $ 2,863,878 |
| Genome troubleshooting and final assembly (predicted) | 2 years (predicted) | 1 FTE | Materials (predicted) | $ 200,000 |
| **Total Final cost** (predicted) | | | | $ **3,223,878** ** |

**Harvard Medical School's overhead is not included in these calculations.

#### Source data

Uncropped gel image of 96 randomly picked colonies from the SynOMICS-based deletion of the parental copy of Segment 2, screened using colony-PCR primers 5'-TTATTGTTTCGACGTCGTTGCCT and 5'-ACTACATGTTCCACACCAAACCG.

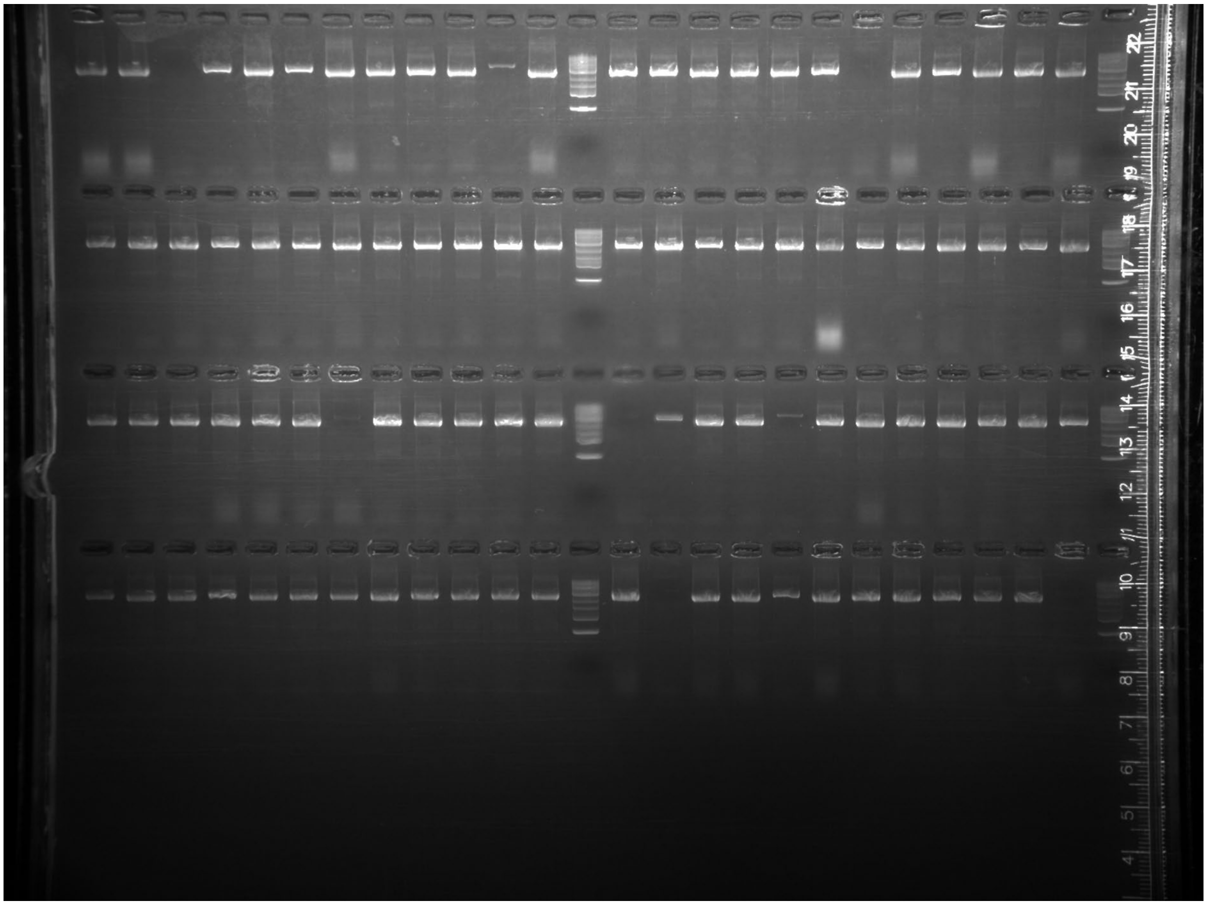

Uncropped gel image of 96 randomly picked colonies from the SynOMICS-based integration of Segment 6, screened using colony-PCR primers 5’-TGGCTCTCCCTTCCCATTATTGT and 5’-CGCAAATCGCCGACATCATTTTT for the presence of the downstream segment-genome junction[11]; and 5’-GTATTATCCAGTCGCATCCGGTG and 5’-TATCCACTTCCACCTGAGTCACC for the presence of the upstream segment-genome junction.

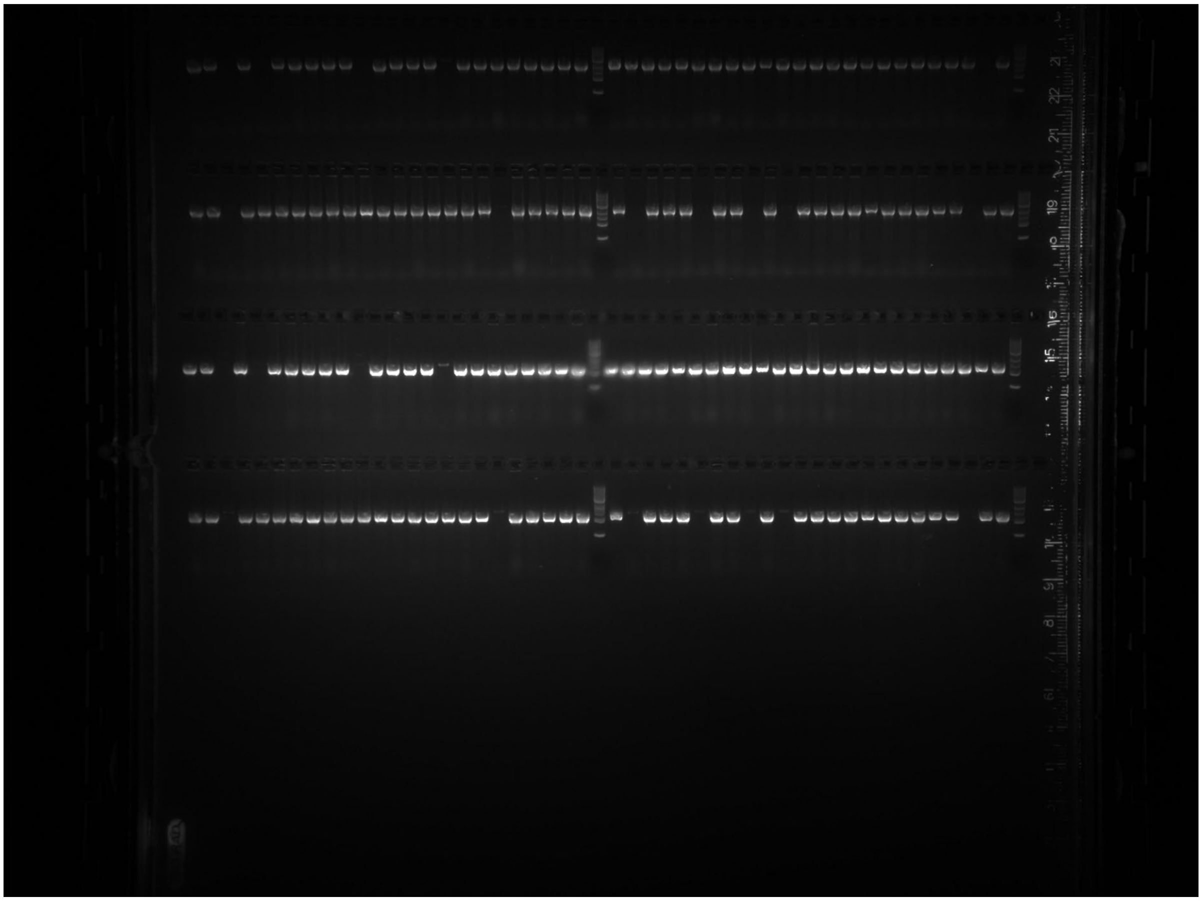

Uncropped gel image of 96 randomly picked colonies from the SynOMICS-based integration of Segment 12, screened using colony-PCR primers 5’-GGTAGAAGTCCCGGTAGGTTTCA and 5’-CATGTTCAGGTCCGGGTTTCATT for the presence of the downstream segment-genome junction[11]; and 5’-TGCTCACCCTCTTTGTCGAGTAT and 5’-TGAAACGCCGTGGTTTAACATCT for the presence of the upstream segment-genome junction.

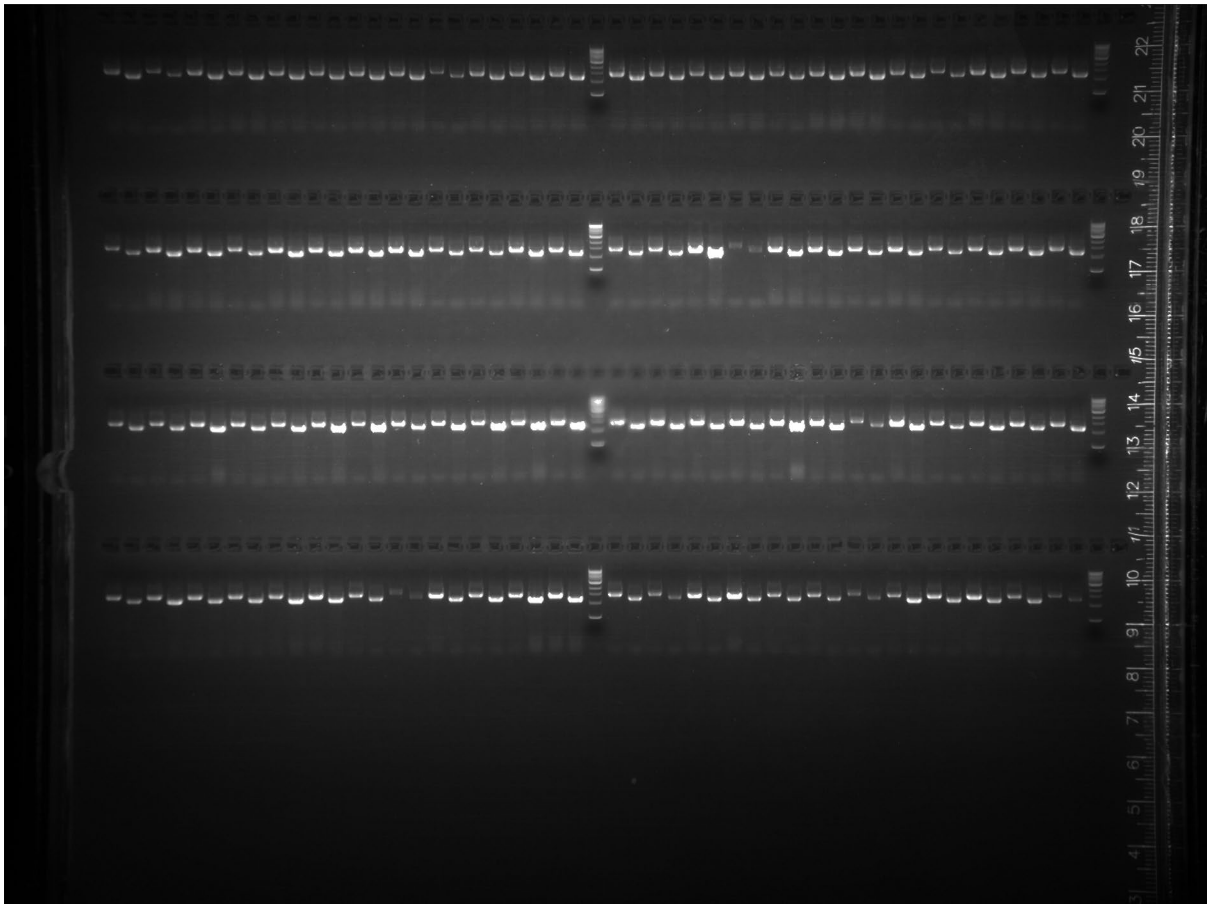

Uncropped gel image of 96 randomly picked colonies from the SynOMICS-based integration of Segment 21, screened using colony-PCR primers 5'-CGCGGGAATAATAAAGAGCACGA and 5'-CGTTGAGGATGCTGAAGGTTGAA for the presence of the downstream segment-genome junction[11]; and 5'-ACCCATTGACAACACGTTCTTGA and 5'-GGAAATTAACGTTGTGTCACGCG for the presence of the upstream segment-genome junction.

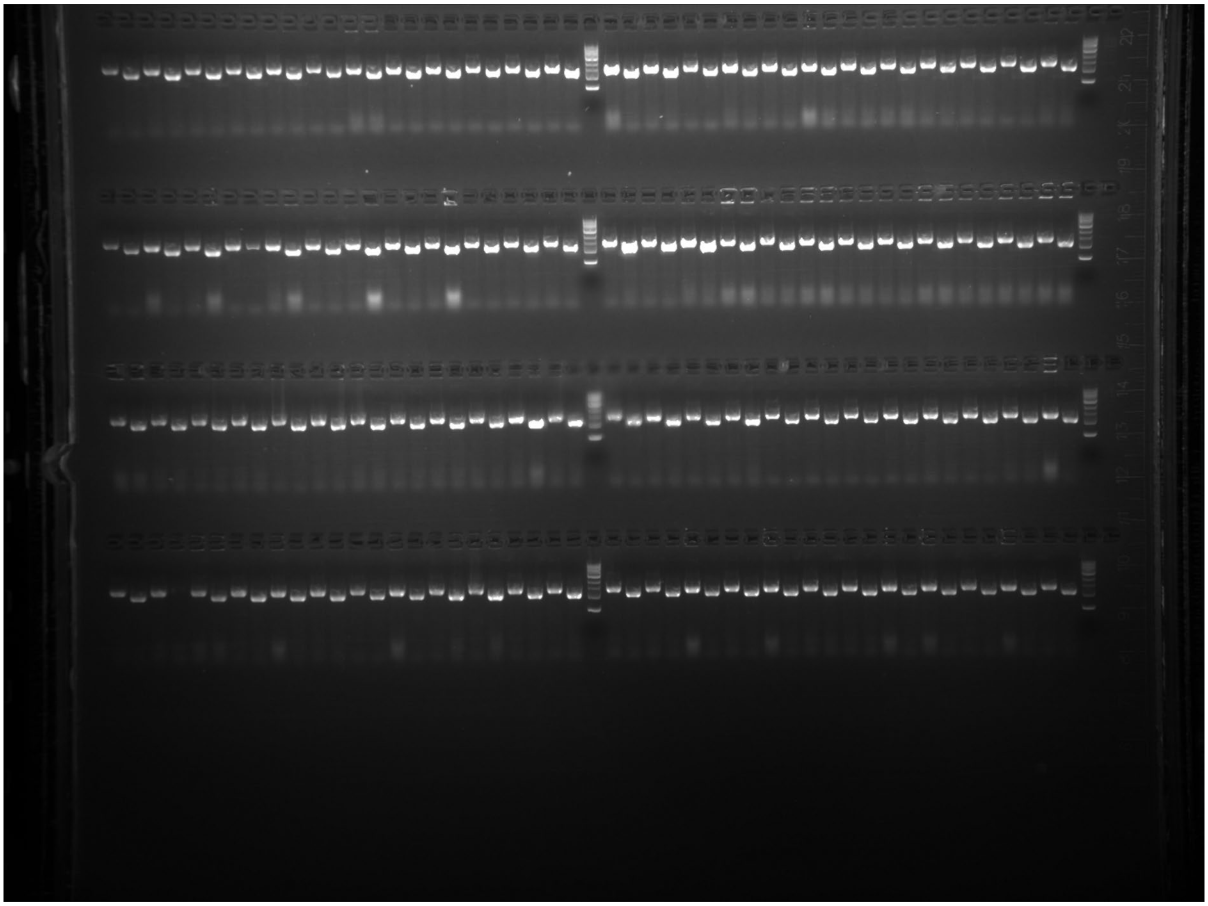

Uncropped gel image of 96 randomly picked colonies from the SynOMICS-based integration of Segment 24, screened using colony-PCR primers 5'-GAAAGTGCCTGGATTGTGACAGT and 5'-GCCTTACCGCCAGAATGATGAAT for the presence of the downstream segment-genome junction[11]; and 5'-ATGTTTGCGCGTTGGGTAAAATC and 5'-CATTCGCCAGATACAGCTCAGTC for the presence of the upstream segment-genome junction.

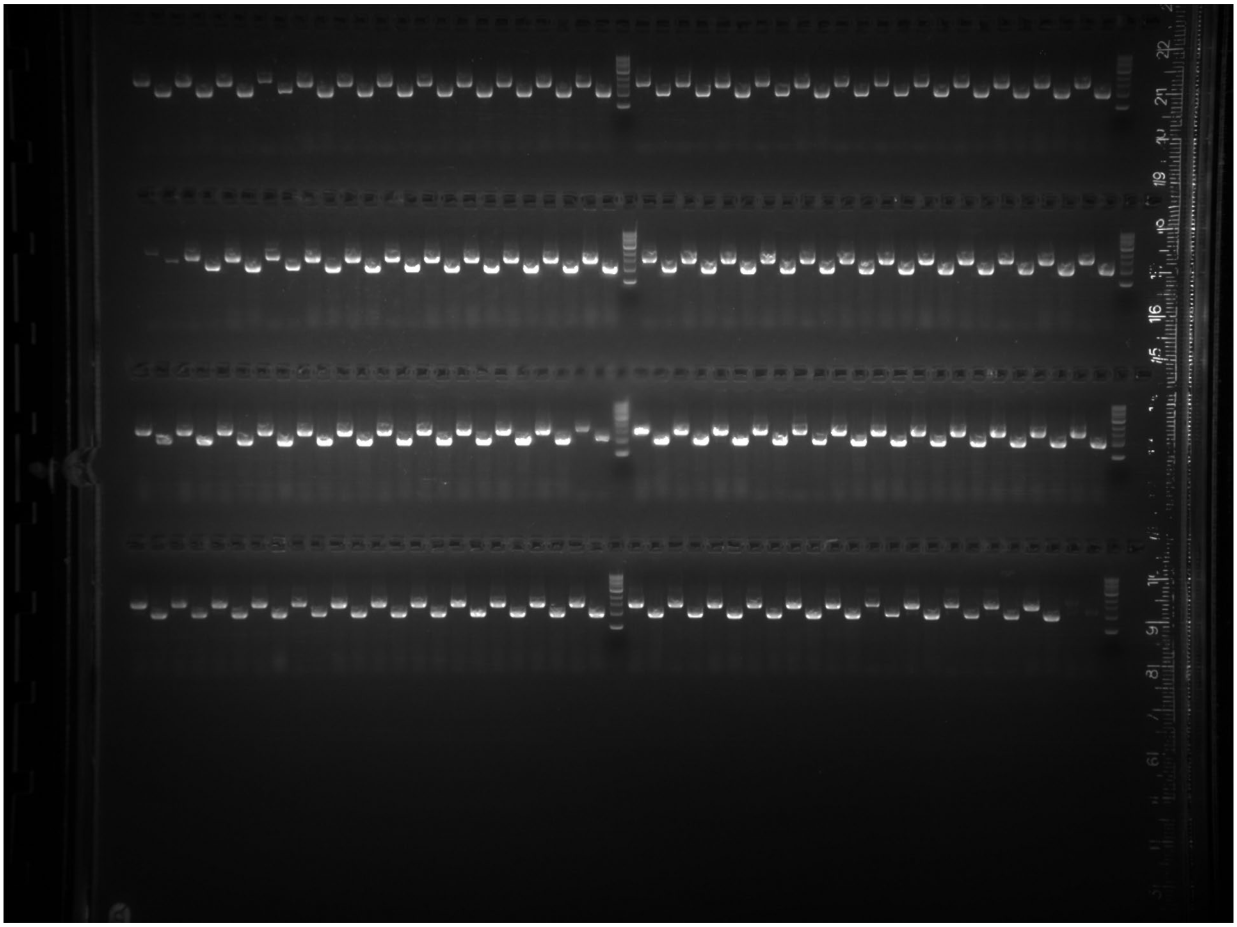

Uncropped gel image of 96 randomly picked colonies from the SynOMICS-based integration of Segment 20, screened using colony-PCR primers 5'-TCCTGTAATTGCCAGCGATTGTT and 5'-CTCCTGTCTGGTCAATCTTTGCC for the presence of the downstream segment-genome junction[11]; and 5'-GTGCTGGCAGTCGATCTGTTAAA and 5'-GGGTACCGATAATCGCCAGAGTA for the presence of the upstream segment-genome junction.

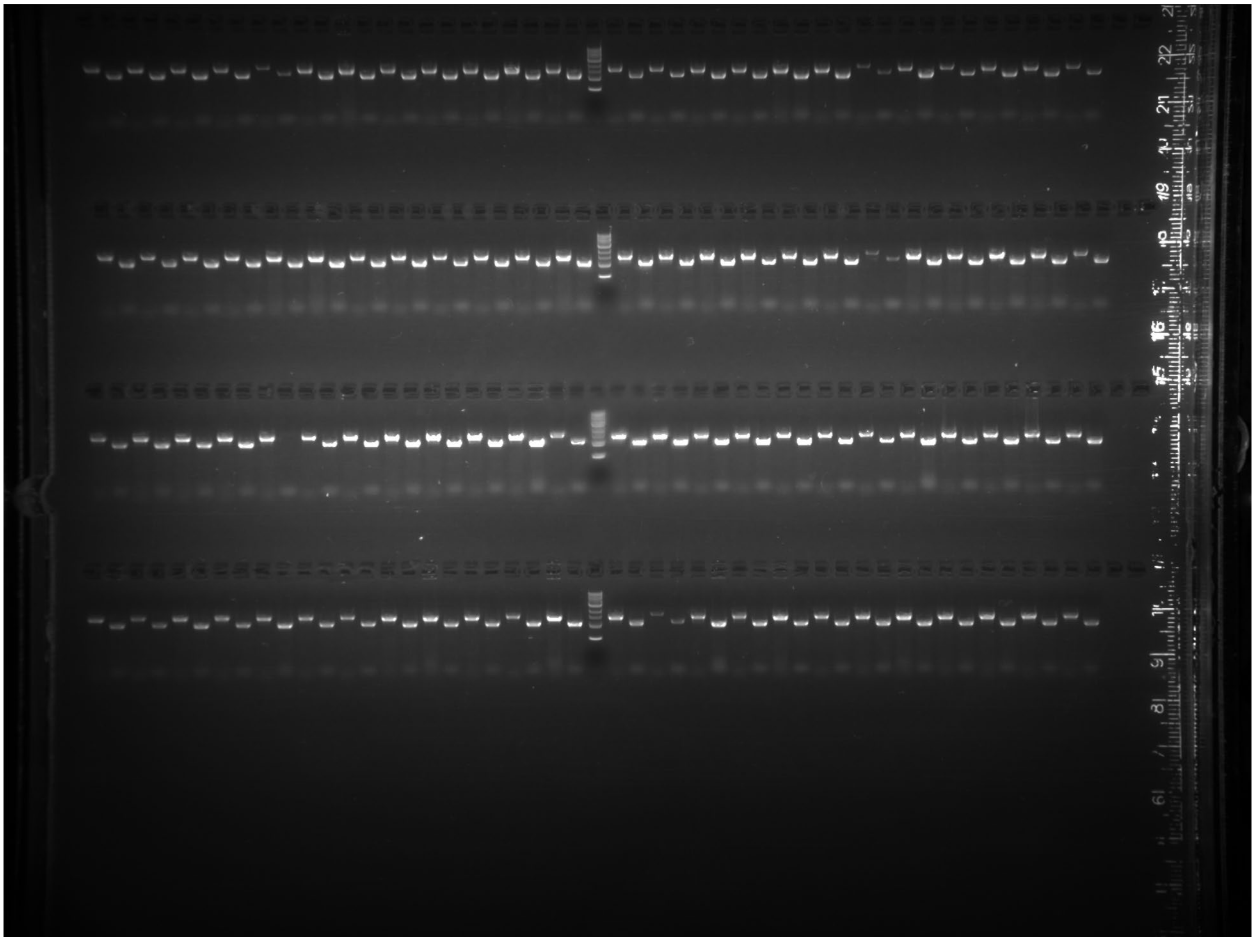

Uncropped gel image of 96 randomly picked colonies from the SynOMICS-based integration of Segment 29, screened using colony-PCR primers 5’-AAATGTCTGACTGGAACCCCTCT and 5’-CGGAAGTGCTGGCTCATTATCTC for the presence of the downstream segment-genome junction[11]; and 5’-AAATGTCTGACTGGAACCCCTCT and 5’-CGGAAGTGCTGGCTCATTATCTC for the presence of the upstream segment-genome junction.

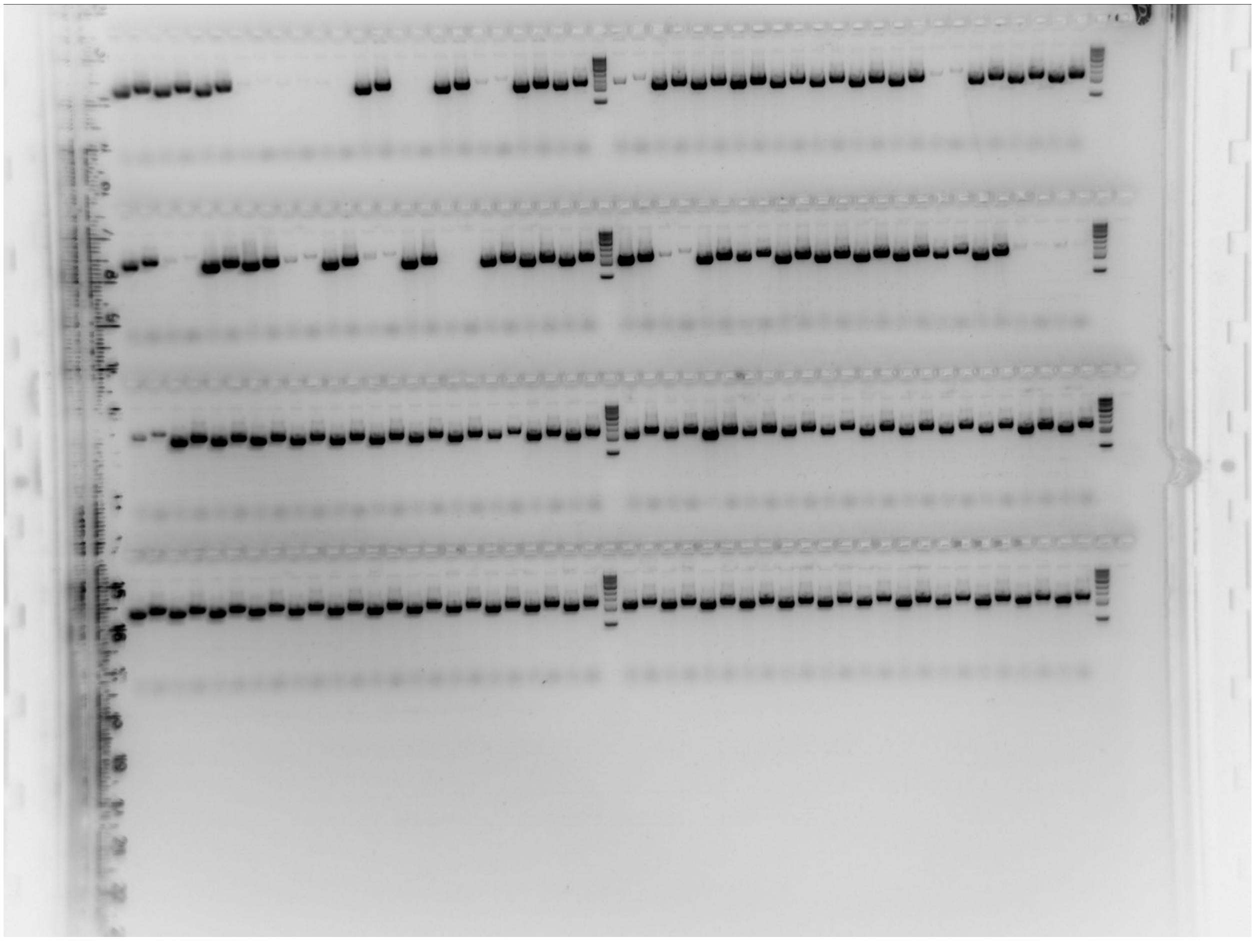

Uncropped gel image of 96 randomly picked colonies from the SynOMICS-based integration of Segment 19, screened using colony-PCR primers 5’-GCCGACGATCGTCACTTTATCAA and 5’-GGGTACCGATAATCGCCAGAGTA for the presence of the downstream segment-genome junction[11]; and 5’-ACGCTGCCTTCTGTCAATGAAAT and 5’-TGGCTGAAATTCAGGAAGACACG for the presence of the upstream segment-genome junction.

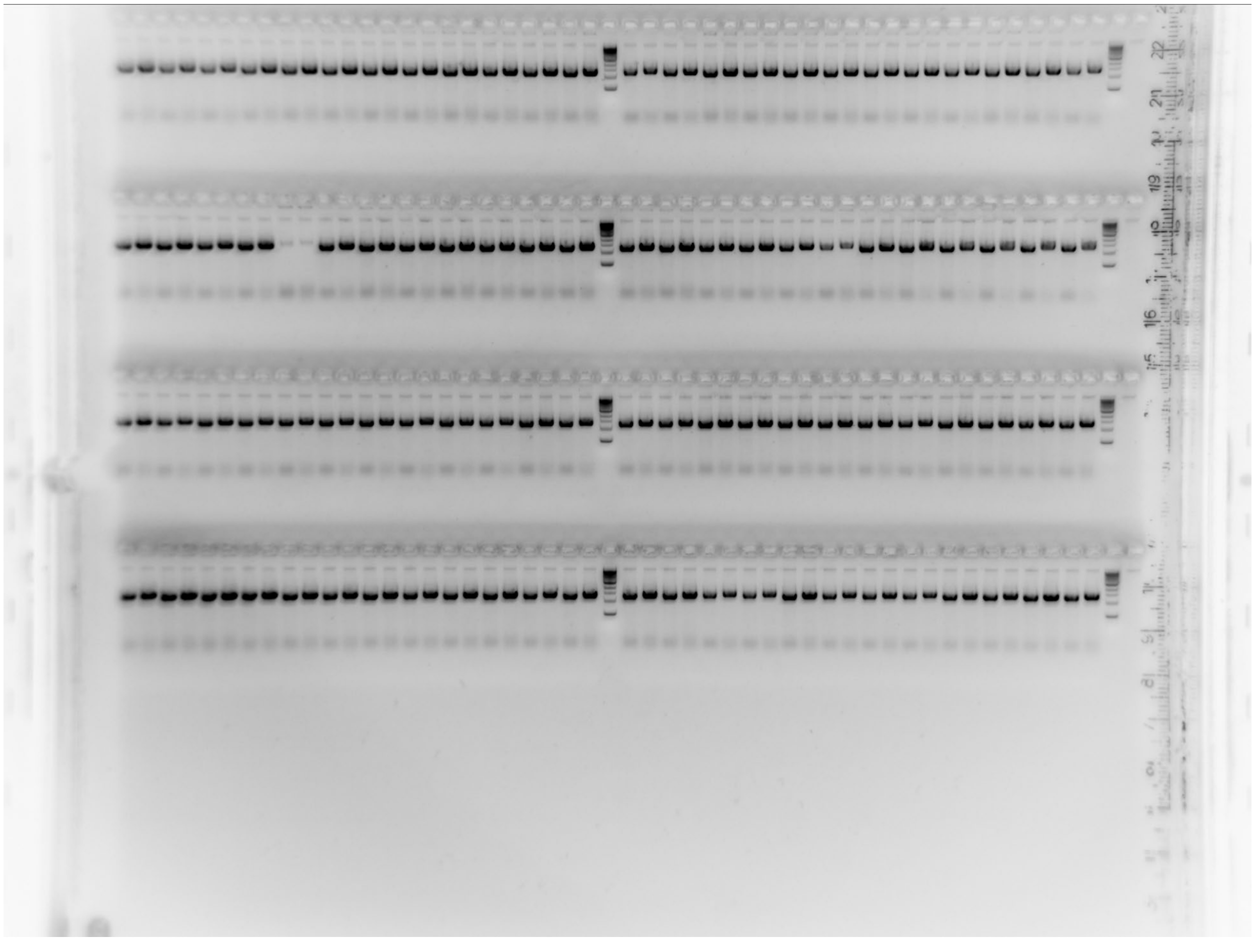

Uncropped gel image of 96 randomly picked colonies from the SynOMICS-based integration of Segment 37, screened using colony-PCR primers 5'-GTGAGGTTCAGGCCACCTTTAAG and 5'-GAATAACGATCGTCGGGTGACTG for the presence of the downstream segment-genome junction[11]; and 5'-GGAGTCTGGCAATCCGTTTATCC and 5'-GCATTGTGCACTTGCGTAAACAT for the presence of the upstream segment-genome junction.

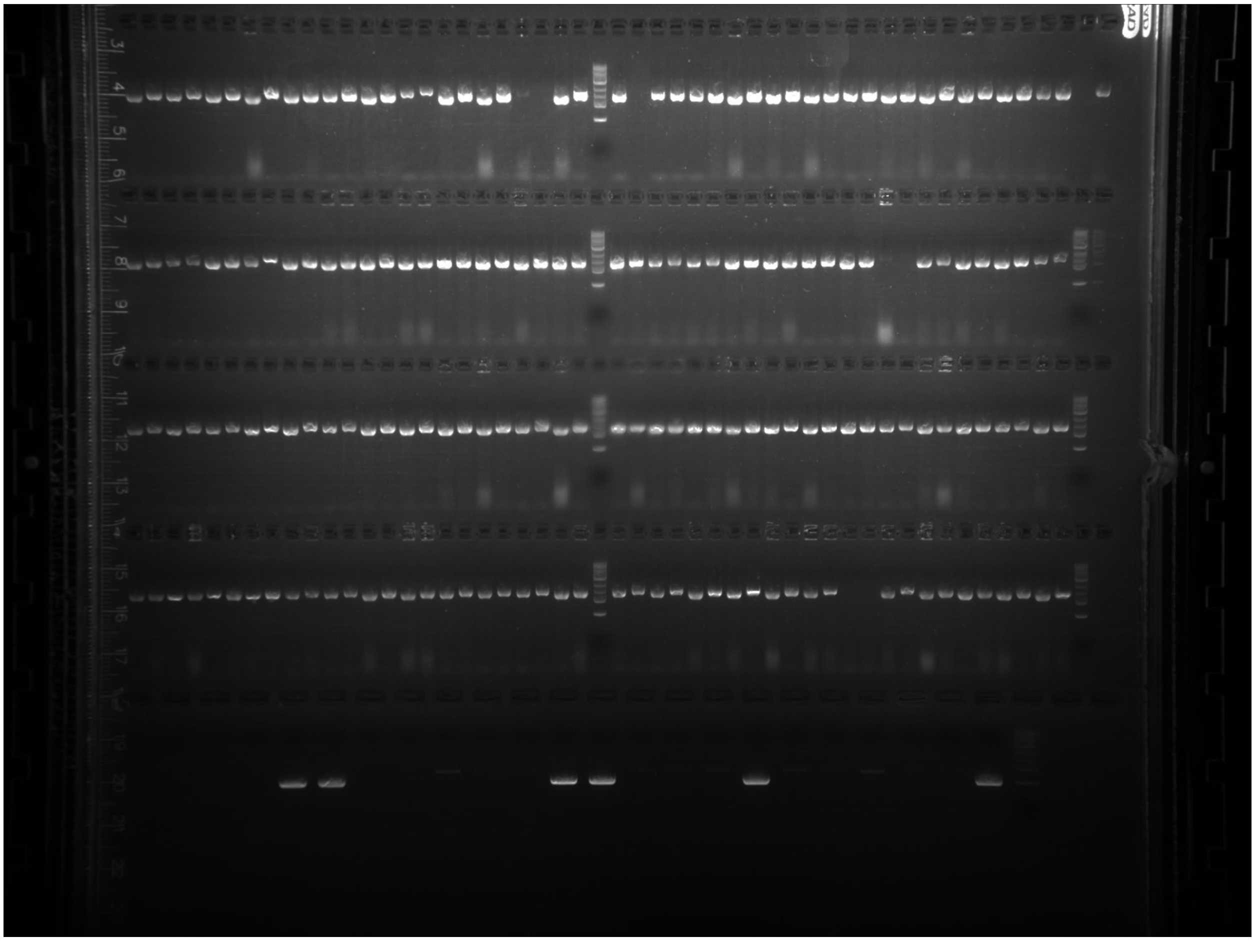

Uncropped gel image of 96 randomly picked colonies from the SynOMICS-based integration of Segment 38, screened using colony-PCR primers 5'-TGTCGTCAGTGACCAGATAACCA and 5'-TGTTCAGAATCACGCATTACCGG for the presence of the downstream segment-genome junction[11]; and 5'-GTCCTGCTGTTTGATGACGTCTT and 5'-GACTTCGCAGTTATCGCCGTATT for the presence of the upstream segment-genome junction.

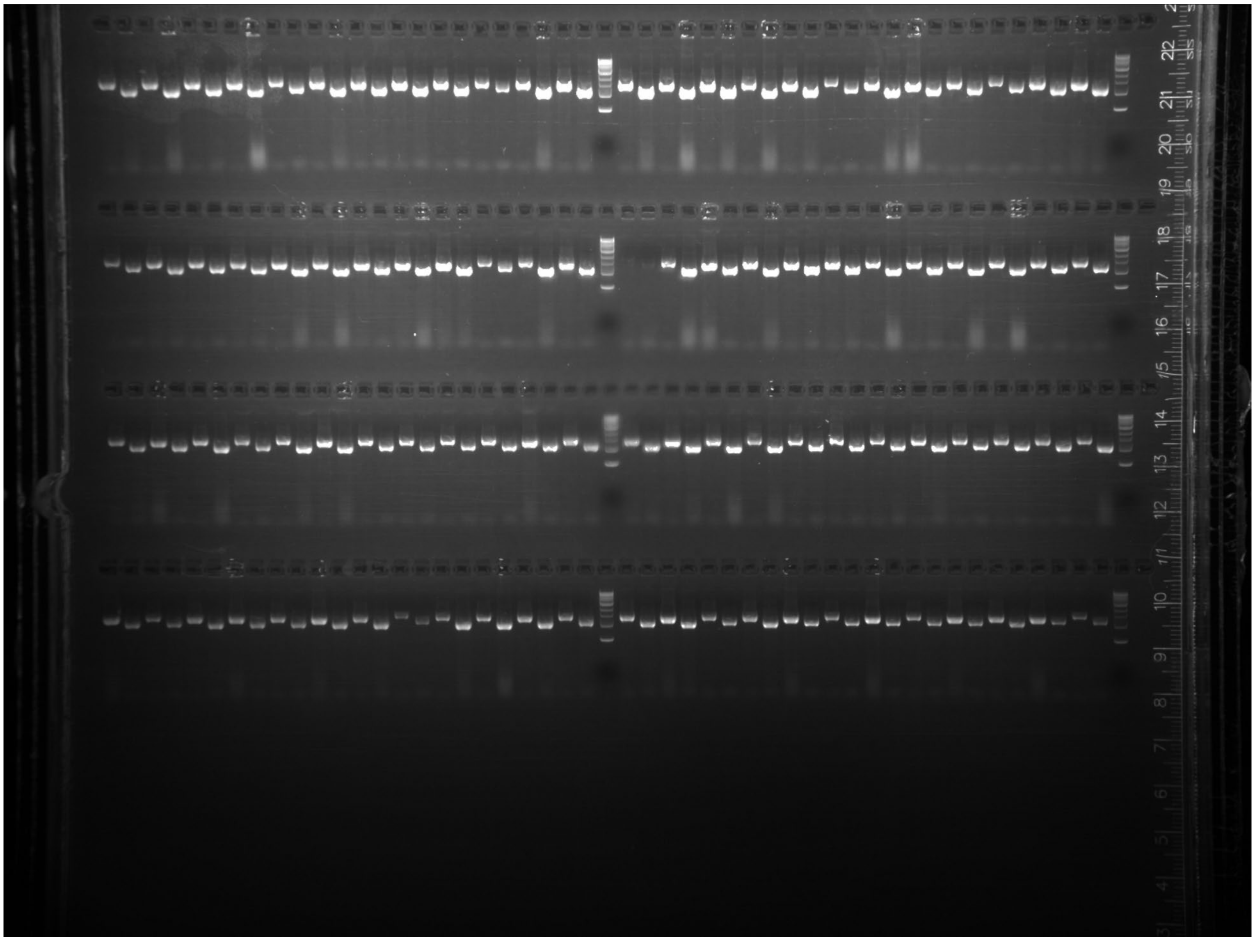

Uncropped gel image of 96 randomly picked colonies from the SynOMICS-based integration of Segment 62, screened using colony-PCR primers 5’-ATTTCCTGCAGAATACCGCCATC and 5’-GAAGAGGTGCATCGTGTTGCTAA for the presence of the downstream segment-genome junction[11]; and 5’-CATGGTCTTTGAGTCTTTCGGCT and 5’-ACTTCGTTTCAAAGTCTCCTCGC for the presence of the upstream segment-genome junction.

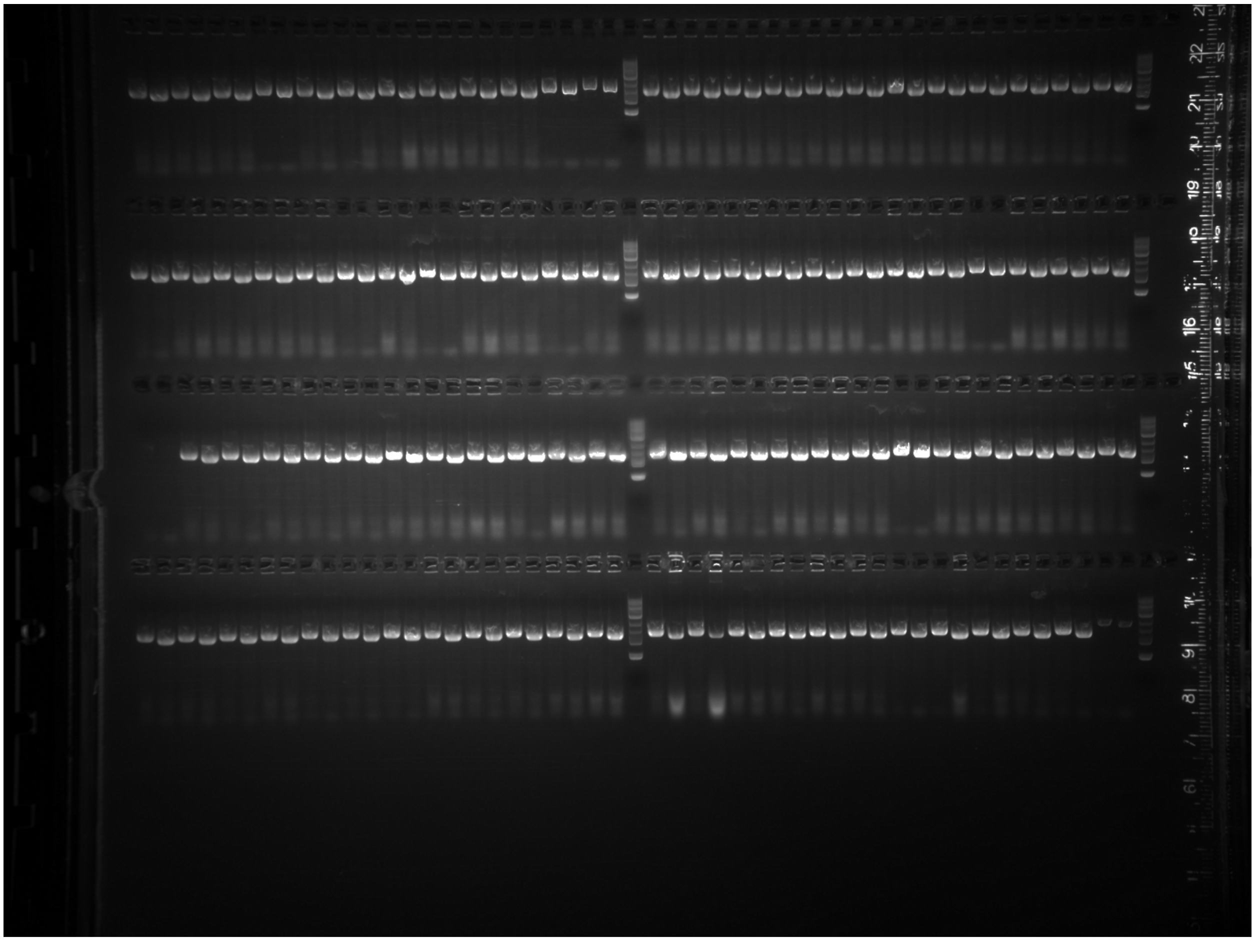

Uncropped gel image of 96 randomly picked colonies from the SynOMICS-based integration of Segment 68, screened using colony-PCR primers 5’-TTTTGCGGTGCCTGATAAGTGAA and 5’-GGTGGAGAAAGATTACACGCTGG for the presence of the downstream segment-genome junction[11]; and 5’-TGTATGCGATTTCACTGGTCGTG and 5’-AACGTTGGTATCTGCGCGATATC for the presence of the upstream segment-genome junction.

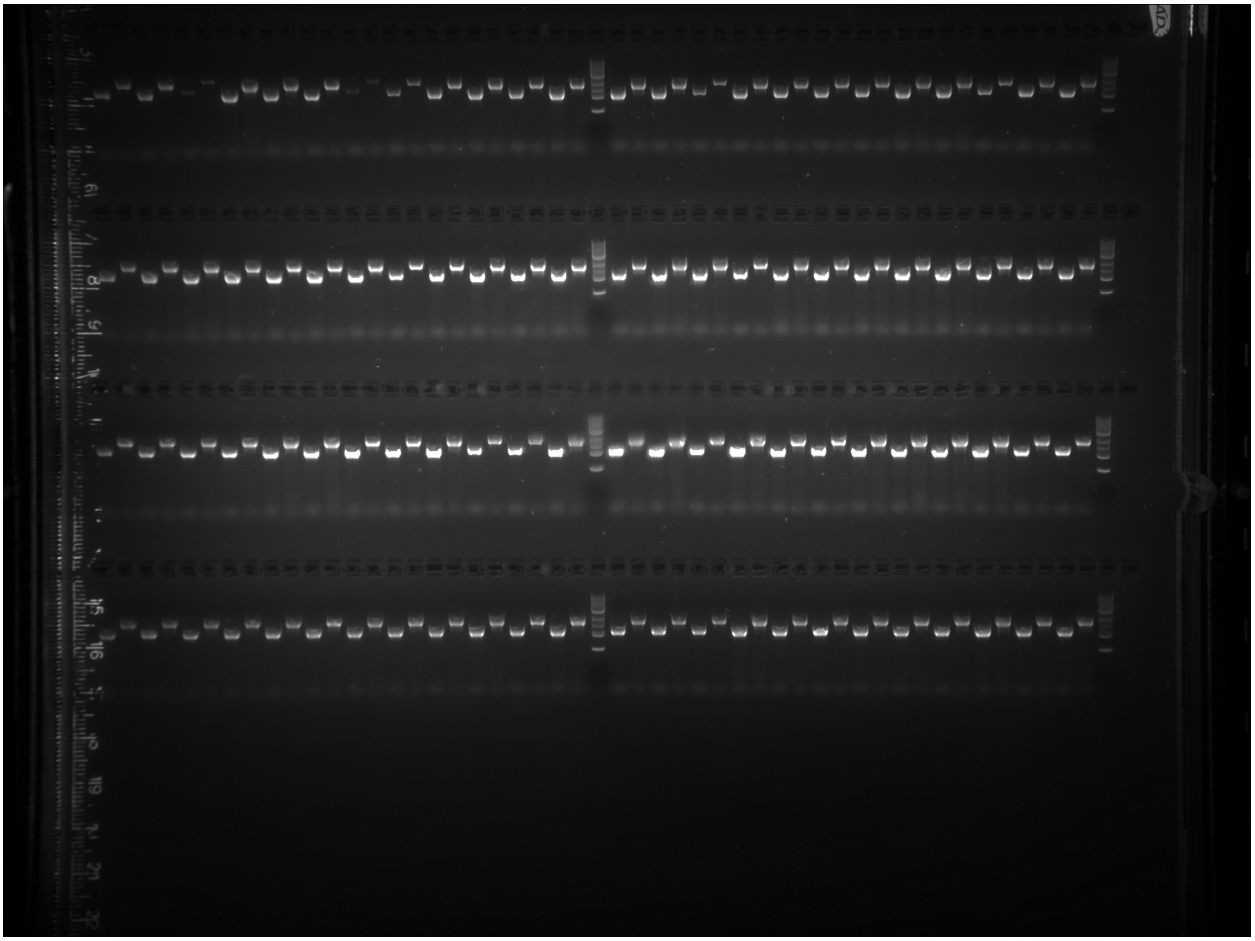

Uncropped gel image of 34 colonies from the SynOMICS-based integration of Segment 85, screened using colony-PCR primers 5’-TTCACTGATGAAGGATGACCGGA and 5’-GAAAATCATTCGCAGCGCTGATC for the presence of the downstream segment-genome junction[11]; and 5’-TCCGAGCCGCTTTCCATATCTAT and 5’-GGCTAACGCAGATCCGATCAAAT for the presence of the upstream segment-genome junction.

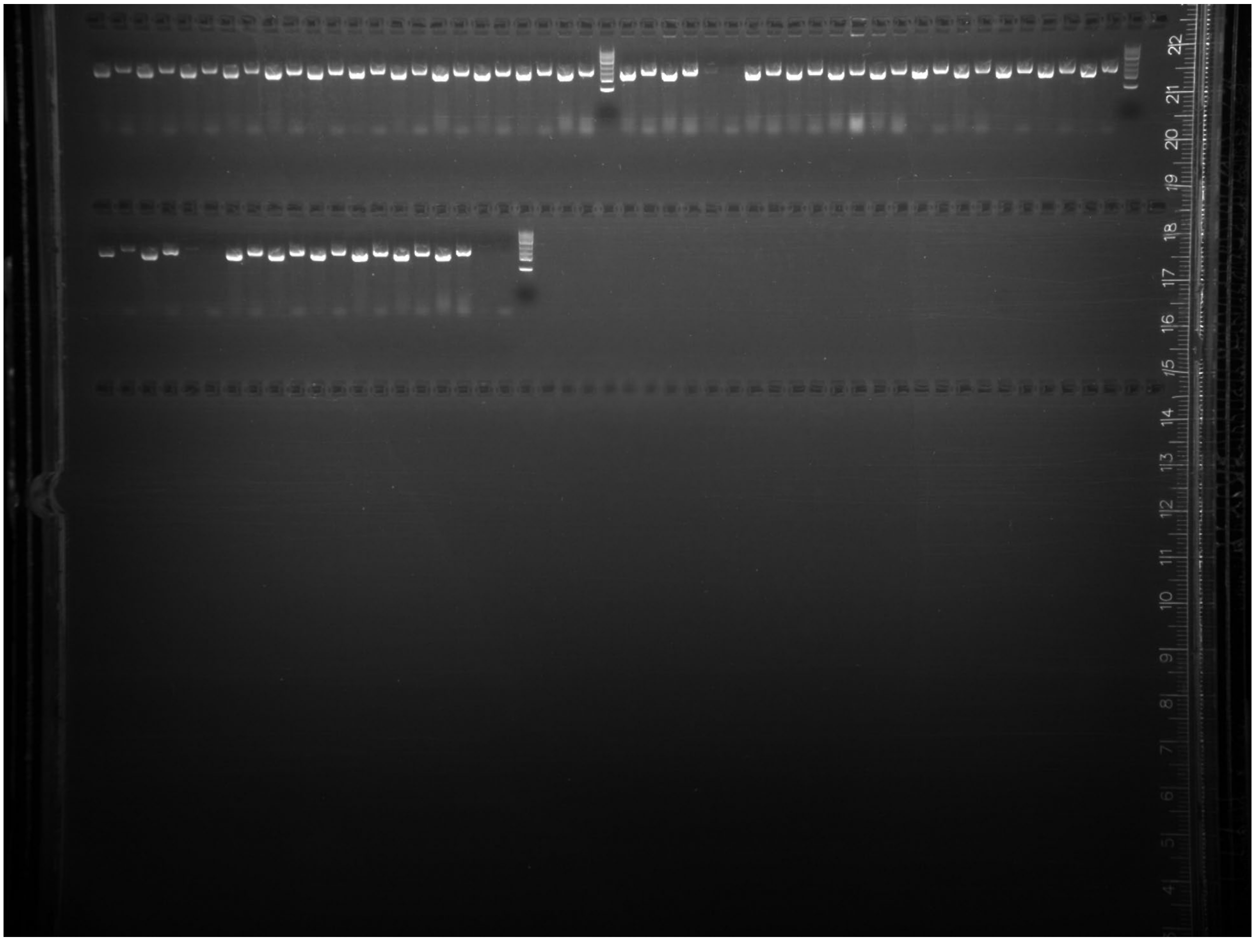

Uncropped gel image of 96 randomly picked colonies from the SynOMICS-based integration of Segment 84, screened using colony-PCR primers 5’-AACCATATGCATACGCCACCTTC and 5’-GGCGCGAGTTGGCTATAATACTC for the presence of the downstream segment-genome junction[11]; and 5’-ACCGAAGTGTCTGAAGCGATCTA and 5’-ACGAGTGGAGTAAGCTTTGAGGA for the presence of the upstream segment-genome junction.

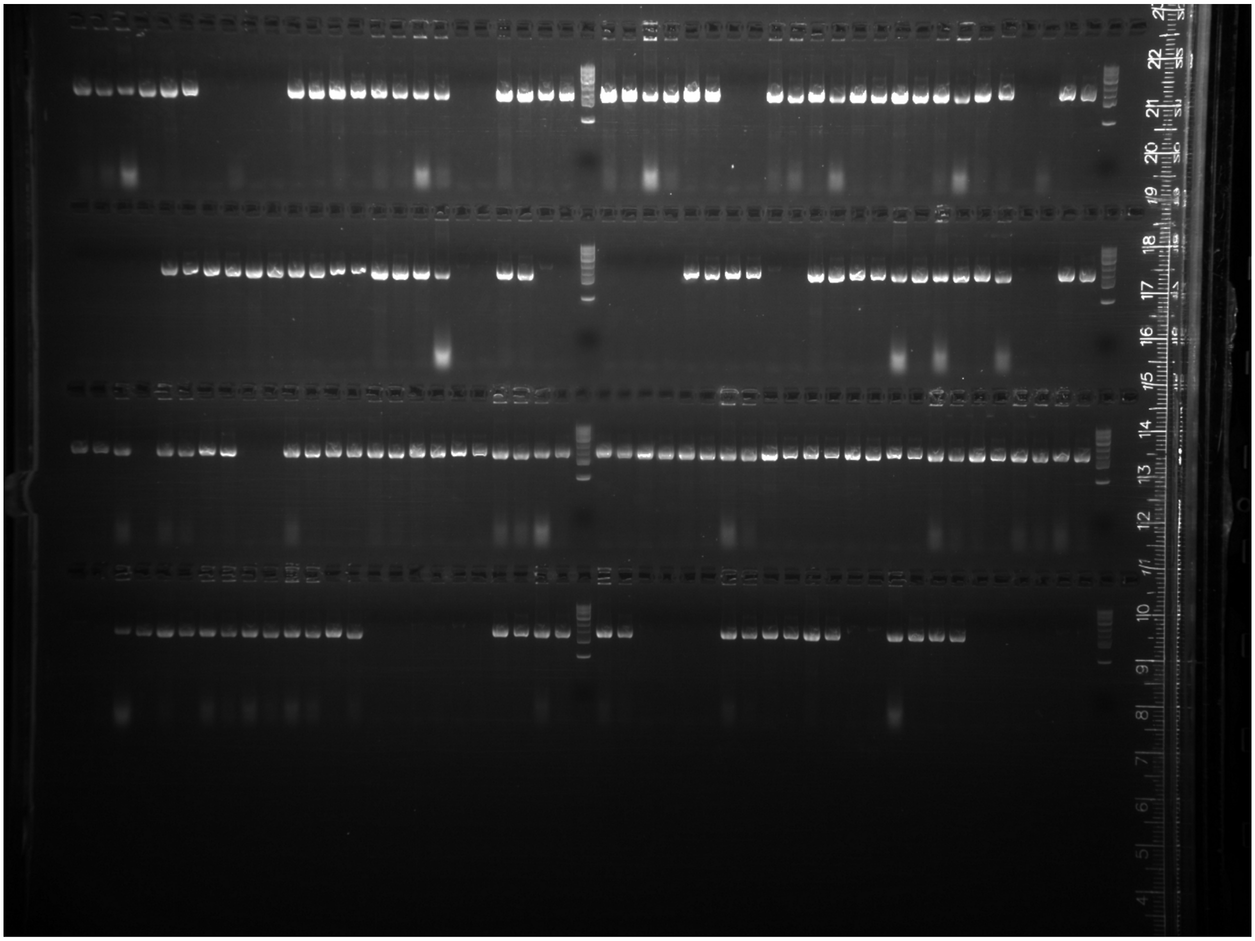
